# Stabilization of the SARS-CoV-2 Spike receptor-binding domain using deep mutational scanning and structure-based design

**DOI:** 10.1101/2021.05.15.444222

**Authors:** Daniel Ellis, Natalie Brunette, Katharine H. D. Crawford, Alexandra C. Walls, Minh N. Pham, Chengbo Chen, Karla-Luise Herpoldt, Brooke Fiala, Michael Murphy, Deleah Pettie, John C. Kraft, Keara D. Malone, Mary Jane Navarro, Cassie Ogohara, Elizabeth Kepl, Rashmi Ravichandran, Claire Sydeman, Maggie Ahlrichs, Max Johnson, Alyssa Blackstone, Lauren Carter, Tyler N. Starr, Allison J. Greaney, Kelly K. Lee, David Veesler, Jesse D. Bloom, Neil P. King

## Abstract

The unprecedented global demand for SARS-CoV-2 vaccines has demonstrated the need for highly effective vaccine candidates that are thermostable and amenable to large-scale manufacturing. Nanoparticle immunogens presenting the receptor-binding domain (RBD) of the SARS-CoV-2 Spike protein (S) in repetitive arrays are being advanced as second-generation vaccine candidates, as they feature robust manufacturing characteristics and have shown promising immunogenicity in preclinical models. Here, we used previously reported deep mutational scanning (DMS) data to guide the design of stabilized variants of the RBD. The selected mutations fill a cavity in the RBD that has been identified as a linoleic acid binding pocket. Screening of several designs led to the selection of two lead candidates that expressed at higher yields than the wild-type RBD. These stabilized RBDs possess enhanced thermal stability and resistance to aggregation, particularly when incorporated into an icosahedral nanoparticle immunogen that maintained its integrity and antigenicity for 28 days at 35-40°C, while corresponding immunogens displaying the wild-type RBD experienced aggregation and loss of antigenicity. The stabilized immunogens preserved the potent immunogenicity of the original nanoparticle immunogen, which is currently being evaluated in a Phase I/II clinical trial. Our findings may improve the scalability and stability of RBD-based coronavirus vaccines in any format and more generally highlight the utility of comprehensive DMS data in guiding vaccine design.

## INTRODUCTION

The rapid development of safe and effective vaccines in response to the SARS-CoV-2 pandemic is a significant achievement of modern vaccinology but has also strained worldwide vaccine manufacturing and distribution capabilities. The success of these pandemic response efforts was made possible by previous prototyping of vaccines against coronaviruses such as MERS-CoV and SARS-CoV (1–3), as well as platform technologies for vaccine delivery (4), emphasizing the importance of continued technology development efforts against possible viral threats. The ability to reliably design, robustly produce, and widely distribute vaccines against SARS-CoV-2 variants and other pandemic threat coronaviruses is a public health priority, and there is room to improve existing vaccines against SARS-CoV-2 in these respects (5). This is particularly true for protein-based vaccines, which are yet to be widely deployed in response to the SARS-CoV-2 pandemic but are likely to become a key part of the global portfolio of SARS-CoV-2 vaccines (6).

Most vaccines against SARS-CoV-2 target epitopes on the Spike protein (S), which is a homotrimeric class I viral fusion protein (7, 8). The S protomer is split into two subunits, S_1_ and S_2_. S_1_ is characterized by an N-terminal domain (NTD) with no known function and a receptor-binding domain (RBD) that facilitates cellular entry by binding the ACE2 receptor, and S_2_ contains the membrane fusion machinery that is held in a metastable prefusion state by the fusion-suppressive S_1_ (9–11). Host receptor identity and mechanisms of recognition vary among coronaviruses, and can make use of either domain in S_1_ depending on the virus (12–15). S is widely considered the most valuable target for protective antibody responses against coronaviruses (2, 16, 17). When used as a vaccine antigen, maintenance of S trimers in their prefusion state is vital for maximizing neutralizing antibody responses (2, 18, 19), and most of the vaccines authorized for emergency use, as well as many current clinical candidates, utilize mutations that stabilize the prefusion conformation (7, 20–22). However, even with stabilizing mutations, most particularly the widely used “2P” mutations (2), protein-based immunogens containing the complete trimeric ectodomain can suffer from instability and low expression yields, which may ultimately limit their development and scalability (23, 24).

The RBD represents a promising alternative to full-length S or trimeric S ectodomains as a target for vaccine development for SARS-CoV-2 and other coronaviruses. The SARS-CoV-2 RBD is more structurally homogeneous than metastable S ectodomains, with far higher expression yields (25, 26). RBD-based immunogens in various oligomeric states have been investigated as genetic or protein-based vaccines for SARS-CoV-2, including monomers (27–31), dimers (32), trimers (33–35) and highly multivalent nanoparticles (25, 36–39), several of which are now being evaluated in clinical trials. Although the RBD comprises only a minority of the total mass and antigenic surface of S and could in principle be more susceptible to escape mutations, this theoretical disadvantage may be mitigated by both functional constraints and the elicitation of antibodies targeting multiple distinct neutralizing epitopes in the RBD by infection or vaccination (25, 40–43). Indeed, the vast majority of neutralizing activity elicited by SARS-CoV-2 infection and currently approved S-based vaccines targets the RBD (40, 44–48), as do most clinical-stage monoclonal antibodies (49). Furthermore, preclinical studies have shown similar reductions in heterologous neutralizing titers against SARS-CoV-2 variants of concern, such as B.1.351, for RBD nanoparticle immunogens and S-based immunogens (37, 50). Finally, RBD antigens can be leveraged to elicit neutralizing responses against the sarbecovirus subgenus through multivalent display on nanoparticles (36, 50) and have recently been shown to be targeted by broadly neutralizing monoclonal antibodies (42, 51–56). In summary, the potential for improved scalability and stability of RBD-based immunogens compared to S-based immunogens, while maintaining similar immunogenicity, affirms the RBD as a *bona fide* antigen for use in protein-based vaccines.

A recent study used deep mutational scanning (DMS) of the SARS-CoV-2 S RBD displayed on the yeast cell surface to identify mutations that enhance RBD expression and stability (57). Such mutations could further improve the manufacturability of RBD-based vaccines and therefore their potential impact on global vaccination efforts. Here, we further combined this DMS data with structure-based design to define stabilizing mutations to the SARS-CoV-2 S RBD, which highlighted the recently-identified linoleic acid-binding pocket as a source of instability in isolated RBDs. Several stabilizing mutations to this region and combinations thereof were found to enhance the thermal and shelf-life stabilities of RBD-containing nanoparticle immunogens and reduce their tendency to aggregate while maintaining their established potent immunogenicity.

## RESULTS

Five mutations previously identified by DMS to strongly improve expression of the SARS-CoV-2 S RBD were considered as starting points for the design of stabilized RBD antigens: I358F, Y365F, Y365W, V367F and F392W (57). A cryo-EM structure of the prefusion S ectodomain trimer (PDB 6VXX) (10) was used to analyze the five mutations in PyMol and Rosetta (58, 59). Only the V367F mutation was found to be exposed to solvent and was therefore not considered for inclusion in stabilized RBD designs to avoid the risk of unfavorably altering antigenicity. The other four mutations were observed to be near or within a recently identified linoleic acid-binding pocket formed between adjacent RBDs in the closed Spike trimer (71, 72), with Y365 identified as a key gating residue for this interaction (**Figure 1A-B**). The improved expression and stability observed by DMS for several mutations in the linoleic acid-binding pocket suggested that this region is structurally suboptimal in the isolated RBD.

**Figure 1.**
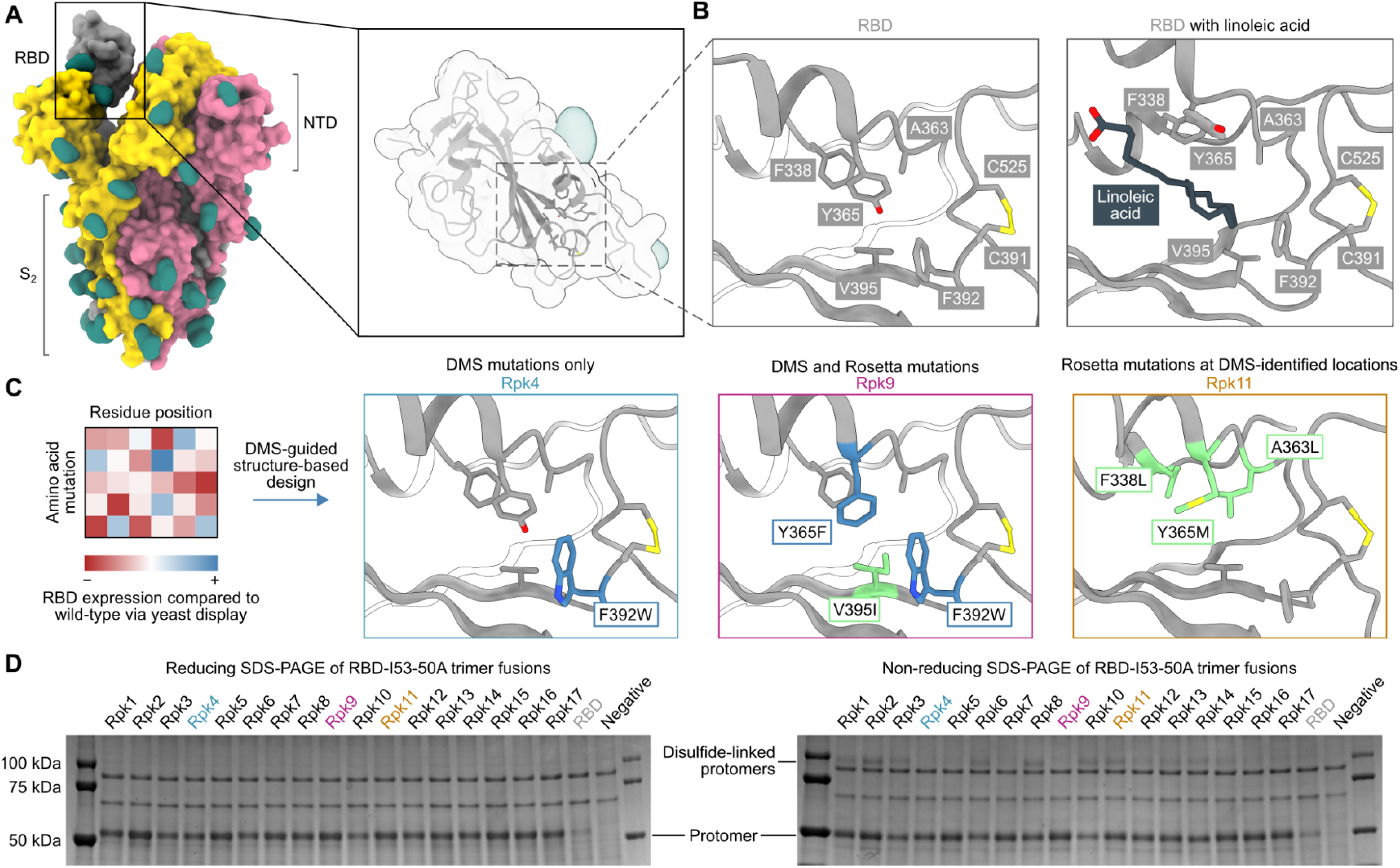
DMS-guided structure-based design of repacked (“Rpk”) SARS-CoV-2 RBDs. **(A)** Molecular surface representation of the SARS-CoV-2 S trimer ectodomain (PDB 6VYB), with a close-up view of the RBD (PDB 6VXX). Each protomer is colored distinctly, and N-linked glycans are rendered dark green. **(B)** The linoleic acid-binding pocket within the RBD, which was targeted for stabilizing mutations. The left panel shows the apo structure (PDB 6VXX) and the right panel shows conformational changes with linoleic acid (black) bound (PDB 6ZB5). **(C)** Mutations that increased RBD expression, identified by DMS of the RBD using yeast display (57) were used to guide Rosetta-based design of stabilized RBDs. Structural models of stabilized RBDs were generated from PDB 6VXX for Rpk4 and Rpk9, and PDB 6YZ5 for Rpk11. All experimentally tested stabilizing mutations are shown in **Supplementary Table 1**. **(D)** Cropped reducing and non-reducing SDS-PAGE of supernatants from HEK293F cells after small-scale expression of stabilized RBD designs genetically fused to the I53-50A trimer. “Negative” refers to a negative control plasmid that does not encode a secreted protein. Uncropped gels are shown in **Supplementary Figure 1**.

We explored whether combinations of these mutations, which were previously reported to individually increase expression and stability (57), could further improve these and other properties of the RBD. We developed computational protocols in Rosetta (59, 73) that modelled one or more of I358F, Y365F, Y365W and/or F392W while also allowing nearby residues to mutate (**Figure 1C, Supplementary Information 1**). Design trajectories that did not force inclusion of any of these four validated mutations within the linoleic acid-binding pocket were also performed, instead allowing Rosetta to design novel sets of stabilizing mutations in the same region. All design trajectories were performed both in the context of the complete S ectodomain (PDB 6VXX) and a crystal structure of an RBD monomer (PDB 6YZ5) that showed a subtly distinct backbone conformation in the region surrounding the linoleic acid-binding pocket. Seventeen repacked designs (abbreviated as “Rpk”) possessing mutations that filled cavities and/or removed buried polar groups were selected for experimental analysis, with some of the DMS-identified individual mutations also included for comparison (**Supplementary Table 1**). The designs were screened in the context of genetic fusions of the Wuhan-Hu-1 RBD to the I53-50A trimer, one component of the two-component icosahedral nanoparticle I53-50 (61), to enable their evaluation as vaccine candidates displaying 60 copies of the RBD (25). Stabilized RBD amino acid sequences were therefore cloned into a vector for mammalian expression with the I53-50A sequence C-terminally fused to the antigen and the two domains joined by a 16-residue flexible Gly-Ser linker (**Supplementary Information 2**).

Stabilized designs and wild-type RBD (“RBD”) were secreted from Expi293F cells while fused to the I53-50A trimer. Reducing SDS-PAGE of cell culture supernatants showed increased expression for all designs compared to wild-type (**Figure 1D, Supplementary Figure 1A**). Furthermore, non-reducing SDS-PAGE showed striking differences in the amounts of disulfide-linked dimers formed by each design (**Figure 1D**). Off-target, intermolecular disulfide formation has been previously observed for both monomeric RBDs and trimeric RBD-I53-50A fusion proteins (25, 57). However, designs including the F392W mutation (Rpk4, Rpk5, Rpk7 Rpk9, Rpk16, Rpk17) yielded noticeably lower levels of disulfide-linked dimers than other designs, consistent with previous experimental analyses of monomeric RBDs bearing individual mutations (57). In addition to F392W partially filling the linoleic acid-binding pocket cavity, the proximity of this mutation to the disulfide between C391 and C525 suggests that it disfavors off-target intermolecular disulfide formation involving these cysteines.

Two designs were selected for more detailed analysis both as monomers and I53-50A-fused trimers (**Supplementary Information 2**): Rpk4, which features F392W alone, and Rpk9, which combines F392W with the DMS-identified Y365F to remove the buried side chain hydroxyl group and the Rosetta-identified V395I to refill the resulting cavity with hydrophobic packing (**Figure 1C**). Scaled-up expression from HEK293F cells and purification by immobilized metal affinity chromatography (IMAC) and size exclusion chromatography (SEC) confirmed increased yields for Rpk4 and Rpk9 both as monomers and as fusions to the I53-50A trimer, with Rpk9 showing a clear advantage for I53-50A trimers (**Figure 2A, Supplementary Figure 2A**). All constructs featured low levels of off-target disulfide-linked dimer formation, highlighting the importance of including F392W in stabilized RBD designs. Melting temperatures (T_m_) were measured for both monomers and trimers by nano differential scanning fluorimetry (nanoDSF), monitoring intrinsic tryptophan fluorescence, which showed increases of 1.9–2.4°C for the Rpk4 proteins and 3.8–5.3°C for the Rpk9 proteins compared to their wild-type counterparts (**Figure 2B**). All of the monomeric RBDs were indistinguishable by circular dichroism and appeared to refold after denaturation at 95°C (**Supplementary Figure 2B**). Moreover, hydrogen/deuterium-exchange mass spectrometry (HDX-MS) of the stabilized RBDs fused to the I53-50A trimer showed decreased deuterium uptake in two distinct peptide segments in the linoleic acid-binding pocket compared to the wild-type RBD (**Figure 2C, Supplementary Figure 3**), suggesting increased local ordering in the stabilized designs. By contrast, peptide segments distant from the linoleic acid-binding pocket, including those in the ACE2 binding motif, showed structural order that was unchanged relative to wild-type. To further assess structural order, all three monomeric RBDs were separately mixed with SYPRO Orange dye to measure the exposure of hydrophobic groups (**Figure 2D**). Both Rpk4 and Rpk9 showed decreased signal compared to the wild-type RBD, with Rpk9 yielding the least fluorescence, suggesting that the improved local order of the linoleic acid-binding pocket in the stabilized RBDs results in less overall hydrophobic exposure. Consistent with the HDX-MS data, neither set of stabilizing mutations impacted the antigenicity of the ACE2 binding motif, as assessed by binding of the antibody CV30 (74) which recognizes an epitope that includes K417 (**Figure 2E**). Affinity to the non-neutralizing antibody CR3022 (75), which binds the RBD core distal to the ACE2 binding motif and closer to the linoleic acid-binding pocket, was slightly decreased (<3.5-fold). In summary, both sets of stabilizing mutations enhanced expression, thermal stability, and structural order of the antigen while minimally impacting antigenicity, with Rpk9 showing superior improvement in all aspects evaluated for both monomeric RBDs and their genetic fusions to the I53-50A trimer.

**Figure 2.**
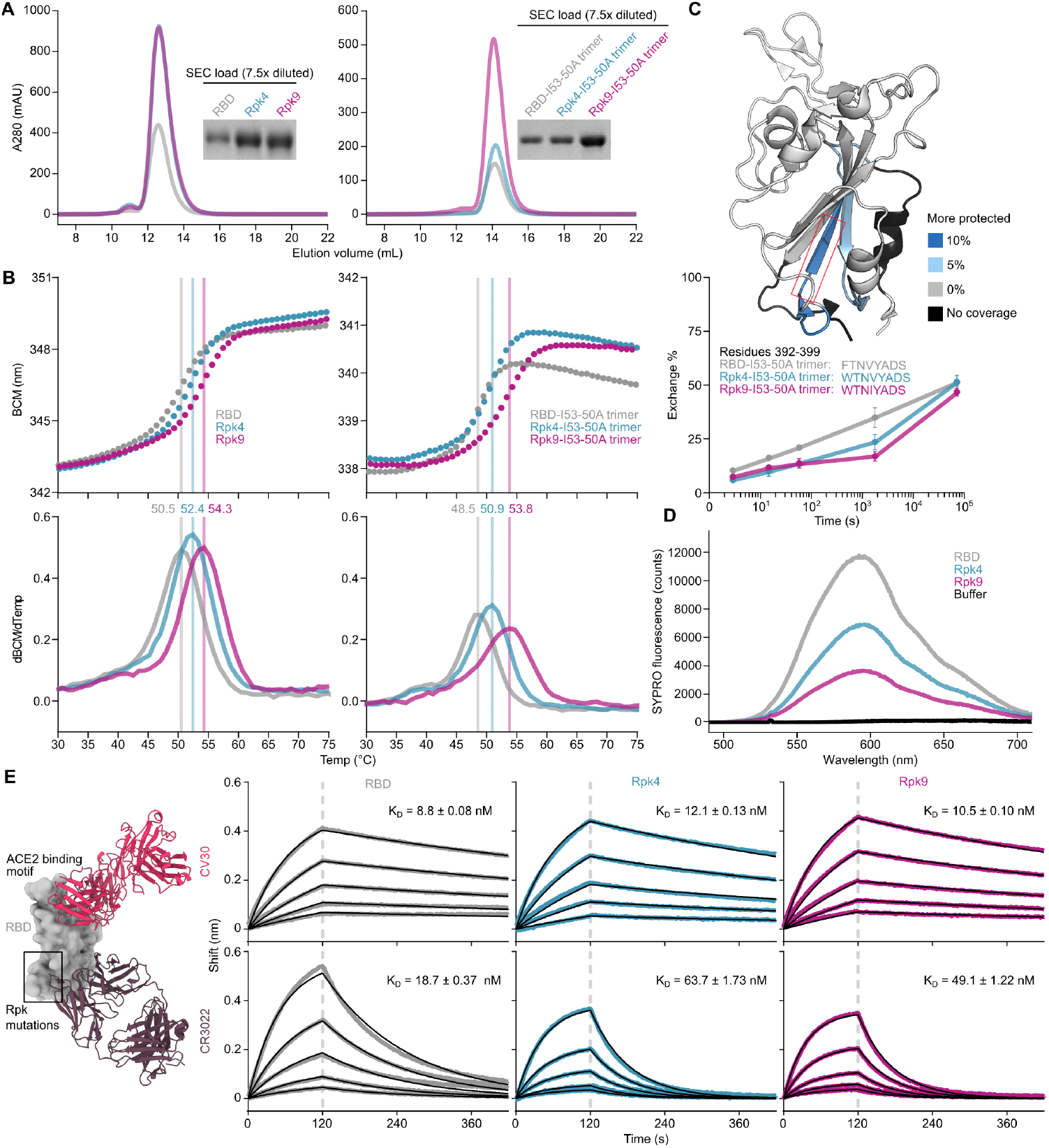
Expression, thermal stability, and structural order of stabilized RBDs is improved while remaining antigenically intact. **(A)** SEC purification of wild-type and stabilized RBDs after expression from equal volumes of HEK293F cultures followed by IMAC purification and concentration. Monomeric RBDs (left) were purified using a Superdex 75 Increase 10/300 GL while fusions to the I53-50A trimer (right) were purified using a Superdex 200 Increase 10/300 GL. Cropped gels show equivalently diluted SEC load samples. Uncropped gels are shown in **Supplementary Figure 2**. **(B)** Thermal denaturation of wild-type and stabilized RBD monomers (left) and fusions to the I53-50A trimer (right), monitored by nanoDSF using intrinsic tryptophan fluorescence. Top panels show the barycentric mean (BCM) of each fluorescence emission spectrum as a function of temperature, while lower panels show smoothed first derivatives used to calculate melting temperatures. **(C)** HDX-MS of wild-type and stabilized RBDs fused to I53-50A trimers. The structural model from PDB 6W41 is shown with differences in deuterium uptake at the 1 minute timepoint highlighted (top). Both Rpk4-I53-50A and Rpk9-I53-50A showed similar increases in exchange protection in similar regions. The red box highlights the peptide segment from residues 392–399, with exchange for this peptide shown at 3 sec, 15 sec, 1 min, 30 min, and 20 h timepoints (bottom). Each point is an average of two measurements. Standard deviations are shown unless smaller than the points plotted. A complete set of plots for all peptide segments is shown in **Supplementary Figure 3**. **(D)** Fluorescence of SYPRO Orange when mixed with equal concentrations of wild-type and stabilized RBD monomers. **(E)** Binding kinetics of immobilized CV30 and CR3022 monoclonal antibodies to monomeric wild-type and stabilized RBDs as assessed by BLI. Experimental data from five concentrations of RBDs in two-fold dilution series (colored traces) were fitted (black lines) with binding equations describing a 1:1 interaction. Structural models (left) were generated by structural alignment of the SARS-CoV-2 bound to CV30 Fab (PDB 6XE1) and CR3022 Fab (PDB 6W41).

Although the stabilizing mutations were designed with isolated RBDs in mind, we also evaluated them in the context of the full S ectodomain. Total yield of the prefusion-stabilized HexaPro antigen fused to T4 fibritin foldon (24) was measured with the Rpk9 mutations (Rpk9-HexaPro-foldon) and compared with the wild-type version (HexaPro-foldon) (**Supplementary Figure 4A-B**). A modest improvement in yield was seen with the Rpk9 mutations, however a slightly earlier SEC elution volume was observed, which could indicate a decrease in stability in the context of S ectodomains (24). Rpk9-HexaPro-foldon showed a similar nanoDSF profile to HexaPro-foldon, although changes in intrinsic fluorescence occurring above 60°C were slightly accelerated with Rpk9-HexaPro-foldon (**Supplementary Figure 4C**). Furthermore, incubation with a 40-fold molar excess of linoleic acid during thermal denaturation revealed substantial changes to the nanoDSF profile of HexaPro-Foldon while Rpk9-HexaPro-foldon appeared less impacted. While not conclusive, these results are consistent with a weakened affinity between Rpk9-HexaPro-foldon and linoleic acid due to the stabilizing mutations. Negative stain electron microscopy (nsEM) revealed that the typical prefusion Spike morphology was maintained with the mutations (**Supplementary Figure 4D**). These data indicate that although it is possible to incorporate mutations to the linoleic acid-binding pocket into prefusion S trimers, the stabilizing effects of the Rpk9 mutations appear unique to isolated RBDs.

While monomeric and trimeric RBDs tend to be stable in solution, multivalent display on self-assembling protein nanoparticles has in some cases exposed a latent tendency of the RBD to aggregate (25, 39). We have previously reported that I53-50-based nanoparticle immunogens displaying the wild-type SARS-CoV-2 RBD elicited potent neutralizing antibody responses and protective immunity in mice and NHPs (25, 37). In those studies, excipients such as glycerol, L-arginine, and the detergent 3-[(3-cholamidopropyl)dimethylammonio]-1-propanesulfonate (CHAPS) were used to stabilize preparations of the nanoparticle immunogens. We next investigated whether the RBD-stabilizing mutations would improve the stability of the nanoparticles in simpler buffers. We assembled the wild-type and stabilized RBD-I53-50A trimers into nanoparticles (RBD-I53-50, Rpk4-I53-50, and Rpk9-I53-50) by addition of the complementary I53-50B.4PT1 pentameric component (61) (**Figure 3A**). Excess residual components were removed by SEC using a mobile phase comprising Tris-buffered saline (TBS) with glycerol, L-arginine, and CHAPS, and the formation of highly monodisperse nanoparticles was confirmed by negative stain electron microscopy (nsEM) (**Figure 3B**). The purified nanoparticles were then dialyzed into buffered solutions with fewer excipients to evaluate solution stability before and after a single freeze/thaw cycle (**Figure 3C-E**). In TBS supplemented with glycerol and L-arginine, the wild-type RBD-I53-50 showed minor indications of aggregation by UV-Vis spectroscopy (**Figure 3C**) and dynamic light scattering (DLS) (**Figure 3D**) that were not observed for Rpk4-I53-50 and Rpk9-I53-50. Differences in solution stability were further magnified after dialysis into TBS with only glycerol: Rpk4-I53-50 and Rpk9-I53-50 were both more resistant to aggregation than RBD-I53-50 and better maintained binding to immobilized human ACE2 (hACE2-Fc) and CR3022 (**Figure 3E**). Dialysis into TBS alone showed clear evidence of aggregation of all samples and loss of antigenicity, with Rpk9-I53-50 retaining slightly better antigenicity than RBD-I53-50 and Rpk4-I53-50. The improved solution stability observed for the stabilized RBDs appears consistent with their enhanced thermal stability and structural order, and offers a subtle but important improvement in formulation stability that is highly relevant to manufacturing of future RBD-based nanoparticle vaccines.

**Figure 3.**
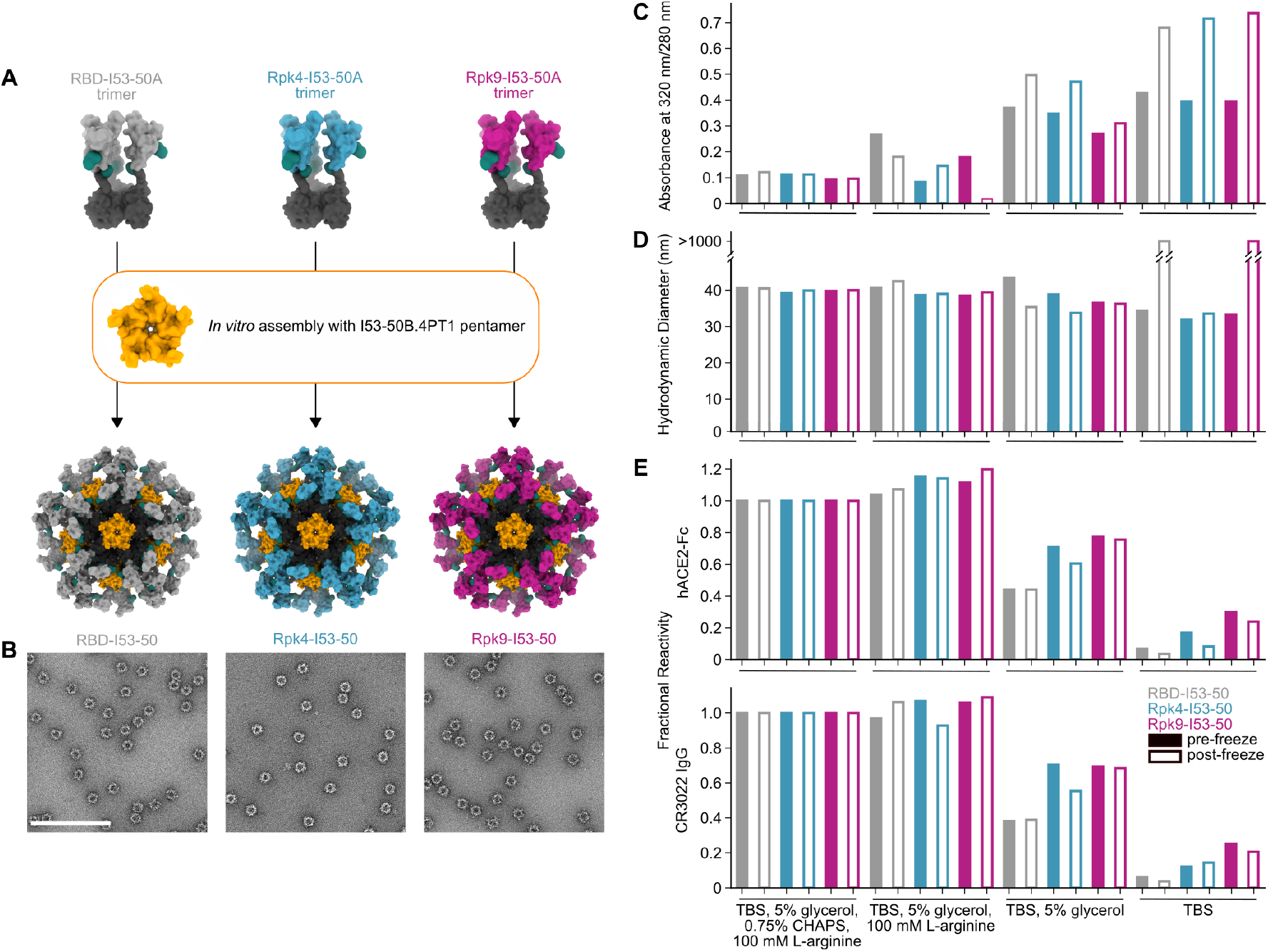
Stabilized RBDs presented on assembled I53-50 nanoparticles enhance solution stability compared to the wild-type RBD. **(A)** Schematic of assembly of I53-50 nanoparticle immunogens displaying RBD antigens. **(B)** nsEM of RBD-I53-50, Rpk4-I53-50, and Rpk9-I53-50 (scale bar, 200 nm). **(C-E)** show summarized quality control results for RBD-I53-50, Rpk4-I53-50, and Rpk9-I53-50 before and after a single freeze/thaw cycle in four different buffers. Complete data available in **Supplementary Information 3**. **(C)** The ratio of absorbance at 320 to 280 nm in UV-Vis spectra, an indicator of the presence of soluble aggregates. **(D)** DLS measurements, which monitor both proper nanoparticle assembly and formation of aggregates. **(E)** Fractional reactivity of I53-50 nanoparticle immunogens against immobilized hACE2-Fc receptor (top) and CR3022 (bottom). The pre-freeze and post-freeze data were separately normalized to the respective CHAPS-containing samples for each nanoparticle.

The immunogenicity of the stabilized RBDs was then evaluated in immunization studies in mice. Immunogens comprising the wild-type and stabilized RBDs were prepared in two formats: I53-50 nanoparticles displaying each antigen (25), and non-assembling controls of nearly equivalent proteins in which the trimeric fusions to I53-50A were mixed with a modified pentameric scaffold lacking the hydrophobic interface that drives nanoparticle assembly (“2OBX”) (76) (**Figure 4A**). In addition to allowing evaluation of the immunogenicity of the different RBDs in trimer and nanoparticle formats, this comparison also directly controls for the effects of nanoparticle assembly. All nanoparticle immunogens were prepared in Tris-buffered saline (TBS) supplemented with glycerol and L-arginine, while the wild-type RBD-I53-50 nanoparticle was also prepared in a buffer that further comprises CHAPS to enable direct comparison to previous immunogenicity studies (25). HexaPro-foldon (featuring the wild-type RBD) was included as a comparator (24). Female BALB/c mice were immunized twice with each immunogen three weeks apart, with serum collection two weeks after each immunization (**Figure 4A**). All doses were administered with equimolar amounts of RBD and included AddaVax adjuvant (77).

**Figure 4.**
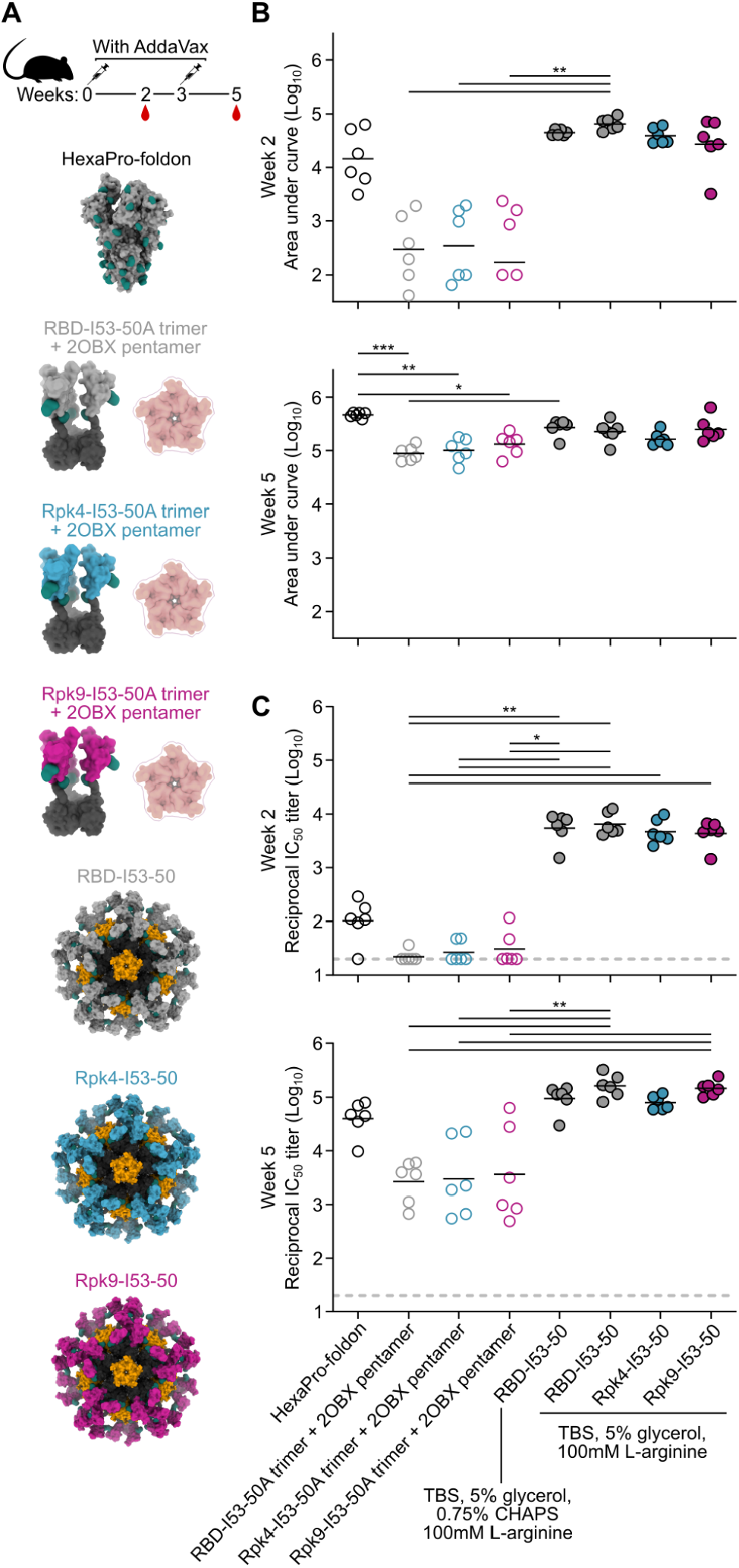
Potent immunogenicity of the parental RBD-I53-50 nanoparticle immunogen is maintained with addition of Rpk mutations. **(A)** Female BALB/c mice (six per group) were immunized at weeks 0 and 3. Each group received equimolar amounts of RBD antigen adjuvanted with AddaVax, which in total antigen equates to 5 μg per dose for HexaPro-foldon and 0.9 μg per dose for all other immunogens. Serum collection was performed at weeks 2 and 5. The RBD-I53-50 immunogen was prepared in two different buffer conditions, with one group including CHAPS as an excipient to bridge to previous studies. **(B)** Binding titers against HexaPro-foldon at weeks 2 and 5, as assessed by AUC from ELISA measurements of serial dilutions of serum. Each circle represents the AUC measurement from an individual mouse and horizontal lines show the geometric mean of each group. One mouse with a near-zero AUC at week 2 for group four was not plotted but still included in the geometric mean calculation. Midpoint titers are shown in **Supplementary Figure 5A**. **(C)** Autologous (D614G) pseudovirus neutralization using a lentivirus backbone. Each circle represents the neutralizing antibody titer at 50% inhibition (IC_50_) for an individual mouse and horizontal lines show the geometric mean of each group. Pseudovirus neutralization titers using an MLV backbone are shown in **Supplementary Figure 5B**. Statistical analysis was performed using one-sided nonparametric Kruskal–Wallis test with Dunn’s multiple comparisons. *, p < 0.05; **, p < 0.01; ***, p < 0.001.

Binding titers were measured against HexaPro-foldon using enzyme-linked immunosorbent assays (ELISA) and analyzed by measuring area under the curve (AUC) (**Figure 4B**) and midpoint titers (**Supplementary Figure 5A**). Sera from all nanoparticle groups showed levels of antigen-specific antibody after the prime that were slightly higher than HexaPro-foldon and markedly higher than the non-assembling controls. Binding signal increased for all groups after the second immunization, with less separation between them. Pseudovirus neutralization using a lentiviral backbone showed similar trends after the prime, with all nanoparticle groups exhibiting significantly higher neutralizing activity than the non-assembling controls and nearly two orders of magnitude more potent neutralization than HexaPro-foldon (**Figure 4C**). Neutralization strongly increased for all groups after the second immunization, with the nanoparticles and HexaPro-foldon showing the highest levels of neutralizing activity. There were no significant differences in neutralizing activity between the various nanoparticle groups or between the various non-assembling control groups at each timepoint. Comparable results were obtained with a different pseudovirus assay using a murine leukemia virus (MLV) backbone (**Supplementary Figure 5B**). These data establish that the stabilized RBDs are similarly immunogenic to the wild-type RBD when presented in either trimeric or particulate formats, with nanoparticle presentation significantly enhancing RBD immunogenicity, most notably after a single immunization.

Improvements to the shelf-life stability of SARS-CoV-2 vaccines have the potential to directly enhance global vaccination efforts by simplifying manufacturing and distribution (78). The stability of the two stabilized RBD nanoparticle immunogens was compared to the wild-type RBD-I53-50 over 28 days of storage at −80°C, 2-8°C, 22-27°C and 35-40°C by DLS (**Figure 5A**), BLI (**Figure 5B**), SDS-PAGE, and nsEM (**Figure 5C, Supplementary Information 4**). No significant deviations from baseline were observed for any immunogen at −80°C, 2-8°C, or 22-27°C over the course of the study. However, storage of the wild-type RBD-I53-50 at 35-40°C for 28 days led to aggregation that was detectable by DLS and nsEM and significant reductions in antigenicity. In contrast, both particle stability and antigenicity were maintained for Rpk4-I53-50 and Rpk9-I53-50 after 28 days of storage at 35-40°C. Collectively, these results establish that the stabilizing mutations we identified in the RBD improve the manufacturability and stability of RBD-based nanoparticle immunogens without compromising their potent immunogenicity.

**Figure 5.**
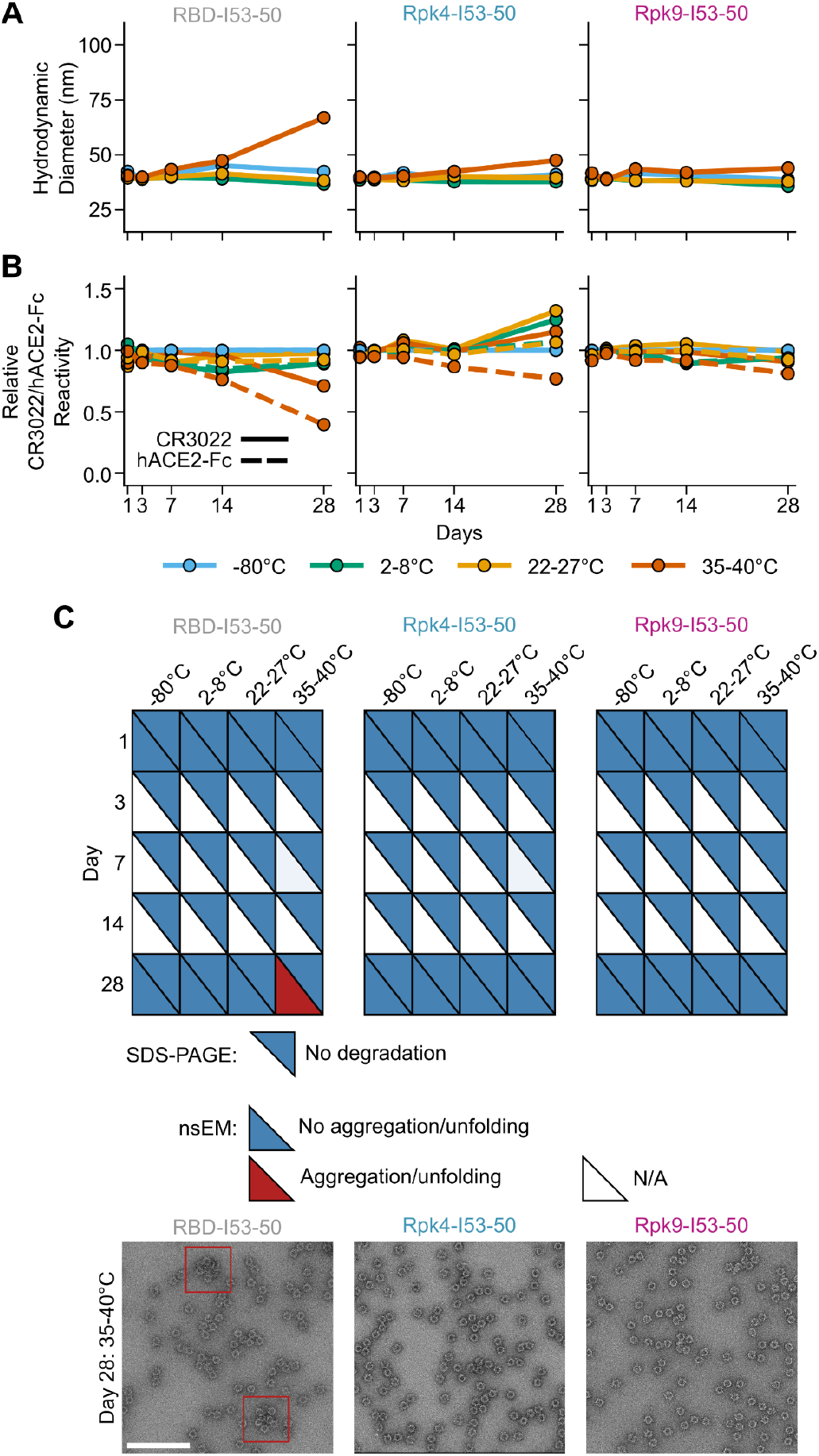
Shelf-life stability of RBD-based nanoparticle immunogens is improved by Rpk mutations. **(A)** Summary of DLS measurements over four weeks. Hydrodynamic diameter remained consistent for all nanoparticles except wild-type RBD-I53-50 at 35-40°C, which showed signs of aggregation after 28 days of storage. **(B)** Binding against immobilized hACE2-Fc receptor (dashed lines) and CR3022 mAb (solid lines) by BLI, normalized to −80°C sample for each time point. Antigenic integrity remained consistent for the stabilized nanoparticle immunogens, while the binding signal of wild-type RBD-I53-50 incubated at 35-40°C decreased by 60% (hACE2-Fc) and 30% (CR3022). **(C)** Summary of SDS-PAGE and nsEM over four weeks. No degradation was observed by SDS-PAGE. Partial aggregation was only observed by nsEM on day 28 for the wild-type nanoparticle stored at 35-40°C. Electron micrographs for day 28 after storage at 35-40°C are shown, with red boxes indicating instances of aggregates (scale bar, 200 nm). All samples were formulated in TBS, 5% glycerol, 100 mM L-arginine. All raw data provided in **Supplementary Figure 4**.

## DISCUSSION

Structure-based protein design can be greatly facilitated by experimental information that narrows the potential design space to particularly valuable regions and mutations. Here, we demonstrate the power of DMS in guiding viral glycoprotein stabilization by characterizing the linoleic acid-binding pocket of the isolated SARS-CoV-2 S RBD as a structurally suboptimal region and identifying individual stabilizing mutations. Guided by these data, structural modeling in Rosetta provided additional stabilizing mutations as well as promising combinations of mutations. All of the experimentally screened designs successfully improved upon the expression of the wild-type RBD, which is an unusually high design efficiency compared to many purely structure-based design experiments. We attribute the high success rate in part to the conservative nature of the mutations tested, but also largely to the prior validation of many of the mutations by DMS. In this case, the DMS data were obtained using yeast display of the RBD which, as opposed to other DMS strategies that measure viral fitness (79, 80), provided a direct readout of RBD expression and folding, making them highly relevant to recombinant forms of the antigen. The stabilizing mutations we identified were distinct from those reported in previous studies focusing on altering surface-exposed positions to improve the expression and stability of the RBD, which were either similarly used to assist in multivalent display of the RBD (39), or alternatively to enhance production in yeast-based expression systems (27, 29–31). Our work instead prioritized stabilizing mutations in a buried, structurally suboptimal region of the RBD. The high success rate of this targeted approach, as well as the substantial improvements we observed in the expression, stability, and solution properties of the Rpk9 variant, suggest that DMS—and particularly the combination of DMS and computational protein design—has considerable promise as a general strategy for identifying stabilizing mutations in glycoprotein antigens.

Our thorough biochemical and biophysical characterization of RBD variants enabled us to select designs that enhanced expression; minimized off-target disulfides; improved local structural order; and increased thermal, solution, and shelf-life stability; all while maintaining the potent immunogenicity of the wild-type RBD displayed on the I53-50 nanoparticle. Of the two mutants studied in detail, Rpk4 (F392W) was more conservative, featuring only a single amino acid change, but less stabilizing compared to Rpk9 (Y365F, F392W, V395I), which included additional mutations that particularly improved expression and thermal stability. We speculate that the improved solution properties of Rpk4- and Rpk9-I53-50 most likely derive from improvements in local structural order and reduced hydrophobic surface area exposure, as indicated by HDX-MS and SYPRO Orange fluorescence. More generally, these results raise the possibility that other RBD antigens may adopt dynamic conformations not observed in existing structures of S ectodomains or isolated RBDs, such as transitions between the open and closed states of the linoleic acid-binding pocket (71, 72).

The similarly potent immunogenicity of Rpk4- and Rpk9-I53-50 compared to the wild-type RBD-I53-50 nanoparticle is consistent with the native-like antigenicity of the ACE2 binding motif, the major focus of neutralizing responses against the RBD (40, 45, 81, 82), and the fact that the stabilizing mutations are not exposed on the surface of the antigen. Our immunogenicity data also clearly demonstrate that high-valency RBD nanoparticle vaccines are far more immunogenic than trimeric forms of RBDs, especially after a single immunization. It is also intriguing that the trimeric S (HexaPro-foldon) elicited higher levels of neutralizing activity than the trimeric non-assembled RBDs. This result demonstrates that removing the RBD from the context of the Spike while maintaining its oligomeric state is not inherently advantageous for improving antibody responses against the RBD, and emphasizes the importance of nanoparticle presentation in RBD-based vaccines.

While the stabilizing mutations reported here are not expected to impact the effectiveness of RBD-based vaccines, the improvements in manufacturability, stability, and solution properties we observe could have a significant impact on the manufacturing and distribution of protein-based vaccines for SARS-CoV-2. As SARS-CoV-2 vaccines are already being updated in response to antigenic drift (83), such improvements could be crucial for maximizing the scale and speed of vaccine production and buffering against unanticipated changes in the stability or solution properties of antigens derived from novel SARS-CoV-2 isolates. Moreover, improved resistance to denaturation and shelf-life stability at various temperatures could be particularly impactful for reliable distribution in developing countries that lack cold chain infrastructure. Finally, as knowledge of the prefusion-stabilizing “2P” mutations prior to the emergence of SARS-CoV-2 proved critical to pandemic response efforts (1, 2, 84), the ability to reliably improve vaccine manufacturability using stabilizing mutations to the RBD may be an important tool for optimizing vaccine designs against other coronaviruses circulating in zoonotic reservoirs that threaten to cross over to humans.

---

**Supplementary Table 1.** Mutations included in each stabilized RBD design. Mutations are separated into previously reported DMS-identified mutations (57) or mutations identified by Rosetta.

| **Mutation set** | **Mutations identified by DMS** | **Mutations identified by Rosetta** |
| --- | --- | --- |
| Rpk1 | Y365W |  |
| Rpk2 | Y365W | F338L |
| Rpk3 | Y365W | L513M |
| Rpk4 | F392W |  |
| Rpk5 | Y365W, F392W |  |
| Rpk6 | Y365F | F338M, A363L, F377V |
| Rpk7 | Y365F, F392W |  |
| Rpk8 | Y365F | V395I |
| Rpk9 | Y365F, F392W | V395I |
| Rpk10 | Y365W | L513I, F515L |
| Rpk11 |  | F338L, A363L, Y365M |
| Rpk12 | I358F, Y365W |  |
| Rpk13 | I358F, Y365W | F338L |
| Rpk14 | I358F, Y365W | L513M |
| Rpk15 | I358F, Y365F | V395I |
| Rpk16 | I358F, Y365W, F392W |  |
| Rpk17 | I358F, Y365F, F392W | V395I |

**Supplementary Figure 1.**
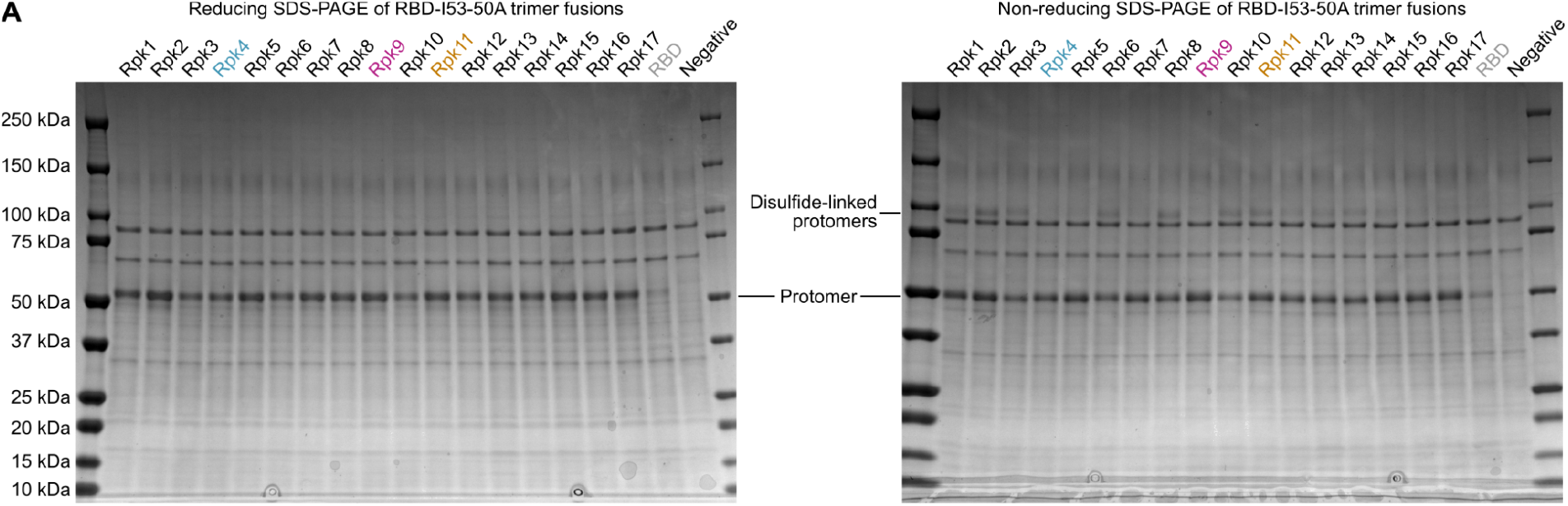
Uncropped reducing and non-reducing SDS-PAGE of supernatants from HEK293F cells during expression of stabilized RBD designs genetically fused to the I53-50A trimer. “Negative” refers to a negative control plasmid that does not encode a secreted protein. Cropped are shown in **Figure 1D**.

**Supplementary Figure 2.**
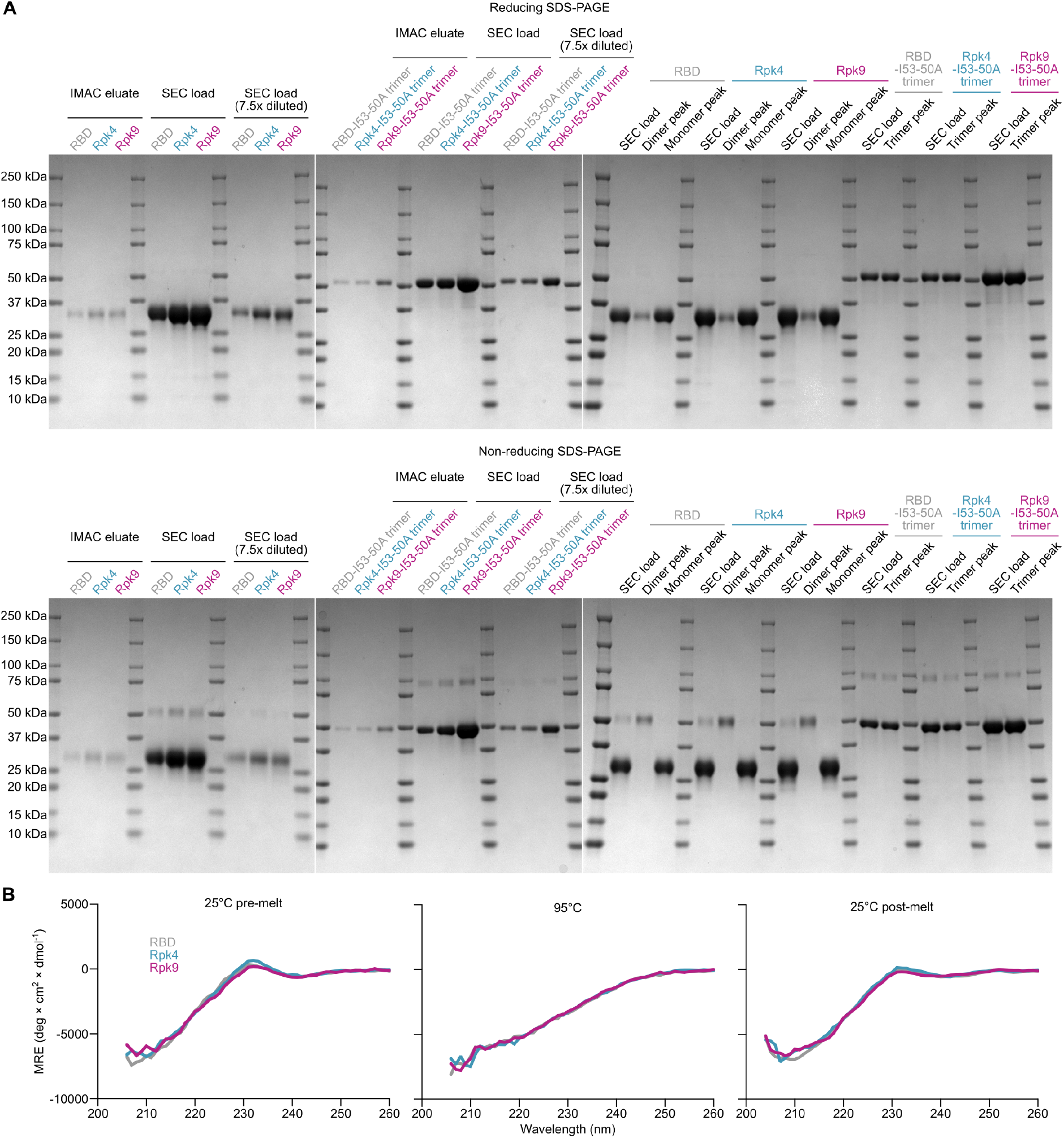
SDS-PAGE of purification of RBD monomers and trimers and circular dichroism of RBD monomers. **(A)** Reducing and non-reducing SDS-PAGE of intermediates and final products during the purification of wild-type and stabilized RBD monomers and genetic fusions to the I53-50A trimer. Selected data are shown cropped in **Figure 2A**. **(B)** Circular dichroism spectra of wild-type and stabilized RBD monomers. Spectra were collected initially at 25°C (left), after raising the temperature to 95°C (center), and again after returning the temperature to 25°C (right).

**Supplementary Figure 3.**
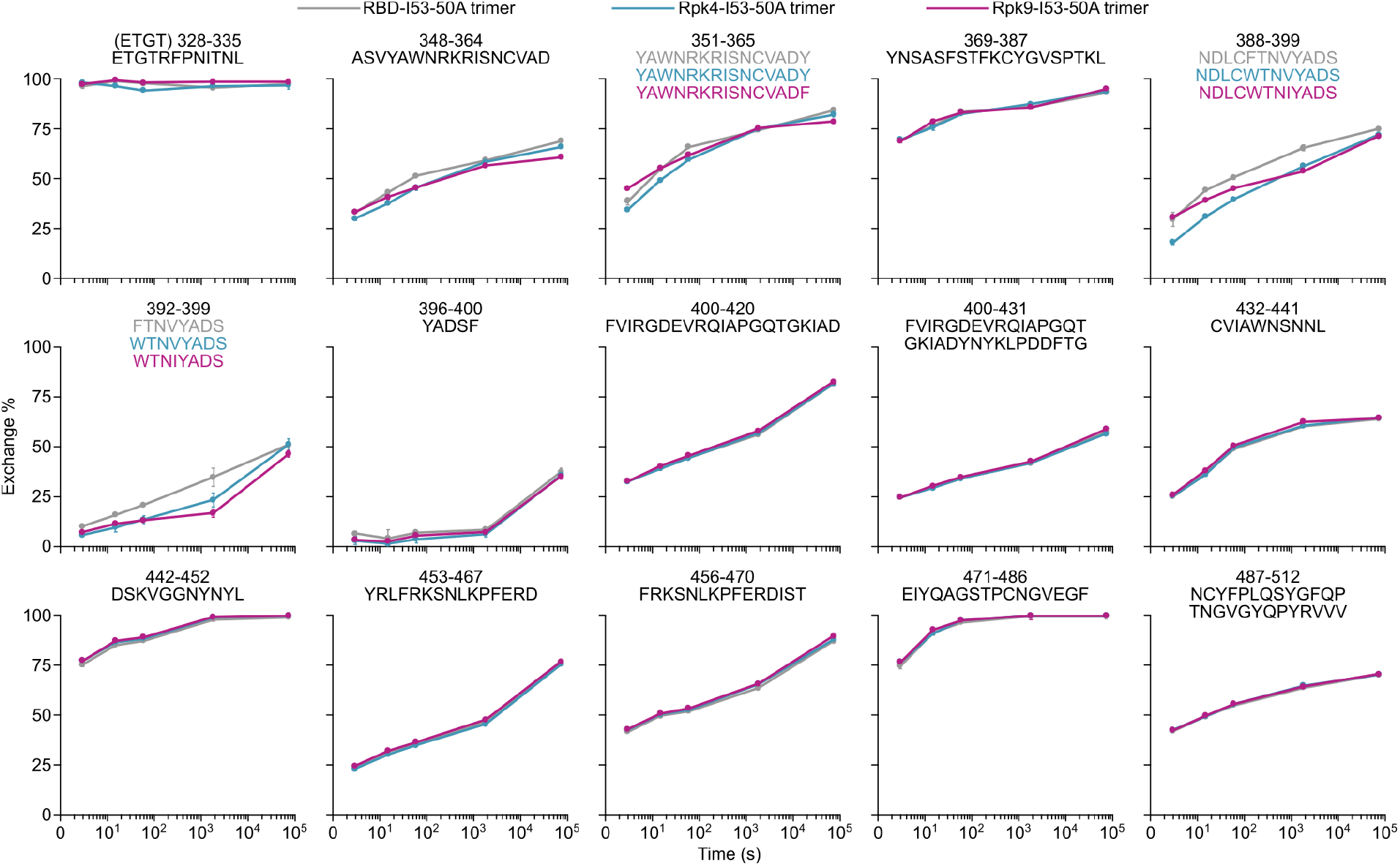
HDX-MS of wild-type and stabilized RBD fusions to I53-50A trimers. Kinetics of deuterium uptake for numbered peptides and leading N-terminal segment (top left plot) in all three constructs are shown for 3 sec, 15 sec, 1 min, 30 min, and 20 h timepoints. Each point is an average of two measurements. Standard deviations are shown unless smaller than the points plotted.

**Supplementary Figure 4.**
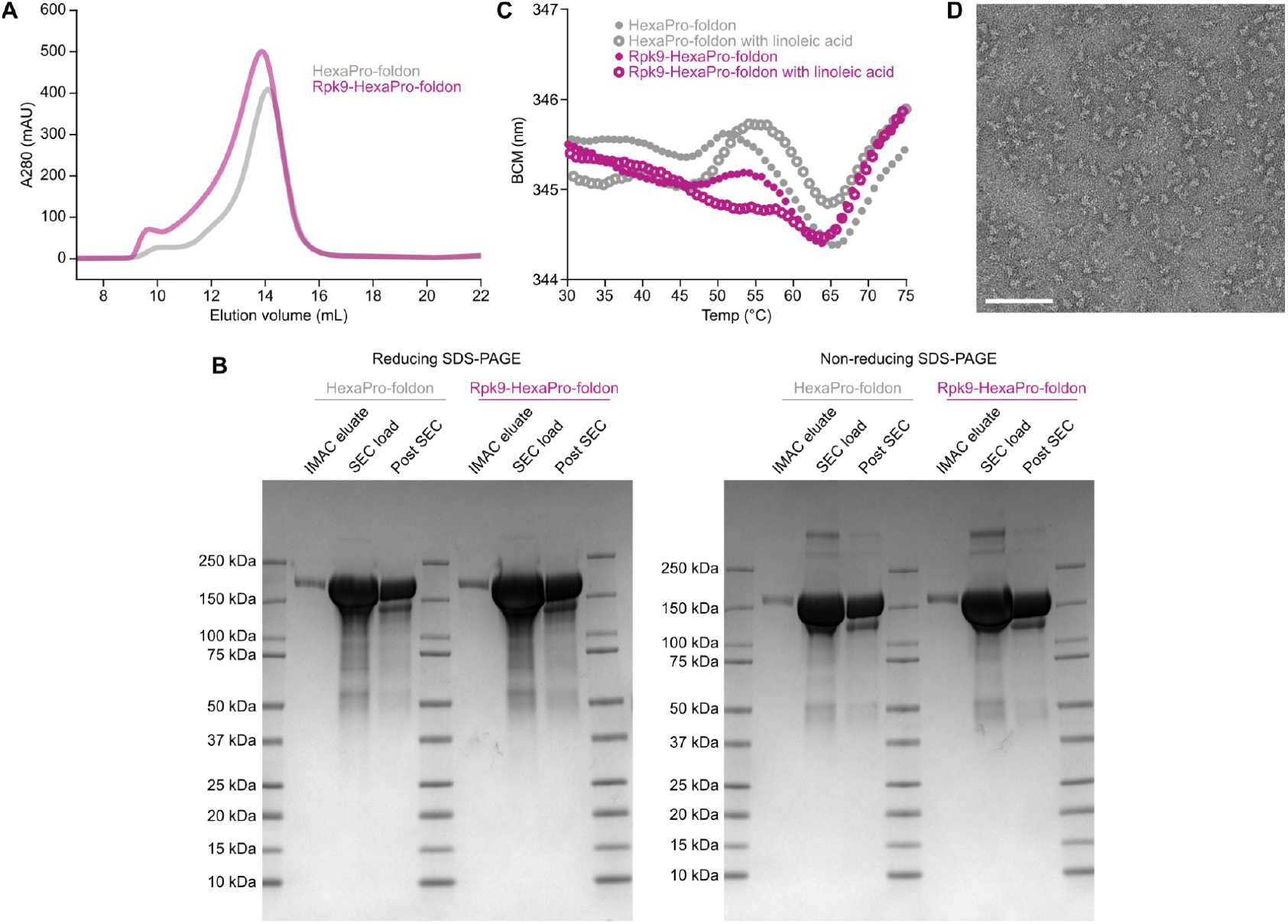
Rpk9 mutations can be incorporated into full length SARS-CoV-2 S ectodomains containing HexaPro mutations. **(A)** SEC purification of wild-type (HexaPro-foldon) and Rpk9 (Rpk9-HexaPro-foldon) prefusion-stabilized S ectodomains after expression from equal volumes of HEK293F cultures followed by IMAC purification and concentration. S ectodomains were purified using a Superose 6 Increase 10/300 GL. **(B)** Reducing and non-reducing SDS-PAGE of intermediates and final products during the purification of HexaPro-foldon and Rpk9-HexaPro-foldon. **(C)** Thermal denaturation of HexaPro-foldon and Rpk9-HexaPro-foldon either in the presence of a 40-fold molar excess (290 μM) of linoleic acid (open circles) or not (closed circles), monitored by nanoDSF using intrinsic tryptophan fluorescence. The barycentric mean (BCM) of the fluorescence emission spectra is plotted as a function of temperature. **(D)** nsEM of Rpk9-HexaPro-foldon (scale bar, 100 nm).

**Supplementary Figure 5.**
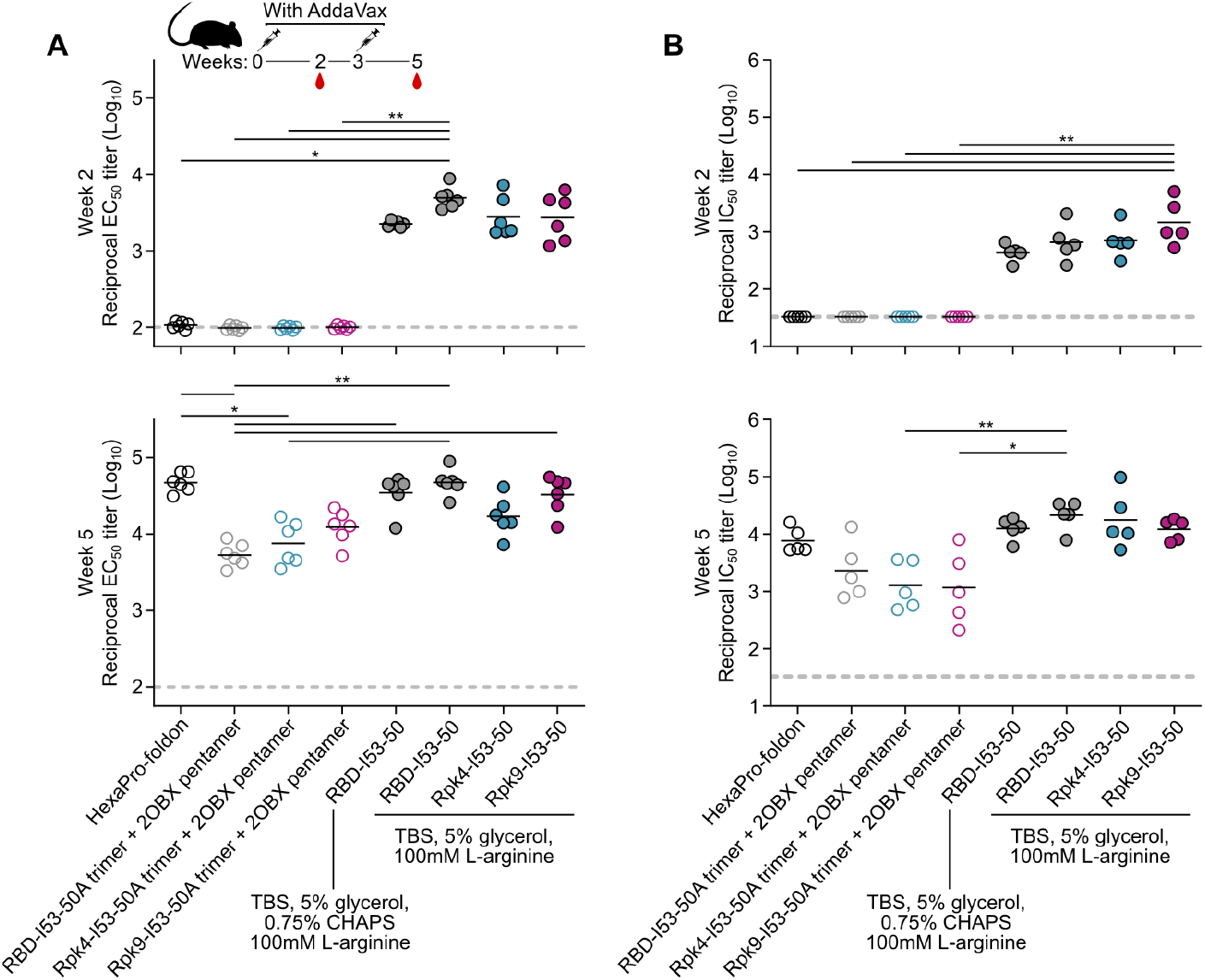
Midpoint binding titers and MLV-based pseudovirus neutralizing titers elicited by RBD immunogens in mice. (A) Serum binding against HexaPro-foldon at weeks 2 and 5 as assessed by midpoint titers (EC_50_) from ELISA measurements. Each circle represents the titer at 50% binding for an individual mouse, and horizontal lines show the geometric mean of each group. (B) Autologous pseudovirus neutralization using an MLV backbone. Each circle represents the neutralizing antibody titer at 50% inhibition (IC_50_) for an individual mouse and horizontal lines show the geometric mean of each group. Five randomly selected mice were analyzed from each group. Statistical analysis was performed using one-sided nonparametric Kruskal–Wallis test with Dunn’s multiple comparisons. *, p < 0.05; **, p < 0.01.

## MATERIALS AND METHODS

### Cell lines

Expi293F cells are derived from the HEK293F cell line, a female human embryonic kidney cell line transformed and adapted to grow in suspension (Life Technologies). Expi293F cells were grown in Expi293 Expression Medium (Life Technologies), cultured at 36.5°C with 8% CO_2_ and shaking at 150 rpm. VeroE6 is a female kidney epithelial cell from African green monkey. The HEK-ACE2 adherent cell line was obtained through BEI Resources, NIAID, NIH: Human Embryonic Kidney Cells (HEK293T) Expressing Human Angiotensin-Converting Enzyme 2, HEK293T-hACE2 Cell Line, NR-52511. All adherent cells were cultured at 37°C with 8% CO_2_ in flasks with DMEM + 10% FBS (Hyclone) + 1% penicillin-streptomycin. Cell lines other than Expi293F were not tested for mycoplasma contamination nor authenticated.

### Mice

Female BALB/c mice four weeks old were obtained from Jackson Laboratory, Bar Harbor, Maine. Animal procedures were performed under the approvals of the Institutional Animal Care and Use Committee of University of Washington, Seattle, WA.

### Design of stabilizing mutations

All calculations in Rosetta were made using version v2020.22-dev61287. All design trajectories assessed the RBD in the closed symmetric trimer conformation observed in a cryo-EM structure of the Spike (PDB 6VXX) and in the context of a crystal structure of the RBD (PDB 6YZ5). The three-fold symmetry axis of PDB 6VXX was aligned with [0,0,1] and a single protomer was saved in .pdb format. An RBD monomer from PDB 6YZ5 was structurally superimposed with the protomer of PDB 6VXX and similarly saved. A design protocol was written using RosettaScripts (58, 59) that takes the aligned protomer and a custom resfile as inputs, with the resfile dictating the side chain identities and conformations sampled during design (**Supplementary Information 1**). Briefly, the protocol applies two rounds of design to a symmetric model based on the input resfile, with side chain minimization applied after each design step. The protocol allows for backbone minimization to be simultaneously performed with side chain minimization, and trajectories were performed either with backbone minimization allowed or disallowed. Both design and minimization steps were allowed to repack or minimize residues within 10 Å of all mutable or packable residues listed in the resfile. Residue positions were manually picked to include positions 358, 365, 392 and surrounding residues based on Spike and RBD structures, and possible residue identities were designated for each position in resfiles using the ‘PIKAA’ option. Resfile inputs were diversified to include various combinations of I358F, Y365F, Y365W, and/or F392W, while also either restricting or allowing mutations to surrounding residues. Further resfile inputs were similarly set up but did not restrict positions 358, 365, and 392 to specific identities. Design models and scores were manually inspected to identify interactions that appeared structurally favorable. Mutations were discarded if they buried polar groups that were natively solvent-exposed or involved in hydrogen bonds. To prevent undesired alterations to antigenicity, mutations to surface-exposed residues were not frequently considered. Favorable sets of mutations were iteratively retested from optimized resfiles and manually refined to finalize a diverse set of designs.

### Plasmid construction

Wild-type and stabilized sequences for the SARS-CoV-2 RBD were genetically fused to the N terminus of the trimeric I53-50A nanoparticle component using linkers of 16 glycine and serine residues. Each I53-50A fusion protein also bore a C-terminal octa-histidine tag, and monomeric sequences contained both Avi and octa-histidine tags. All sequences were cloned into pCMV/R using the XbaI and AvrII restriction sites and Gibson assembly (60). All RBD-bearing components contained an N-terminal mu-phosphatase signal peptide. hACE2-Fc was synthesized and cloned by GenScript with a BM40 signal peptide. The HexaPro-foldon construct used for immunization studies was produced as previously described (24) and placed into pCMV/R with an octa-histidine tag. HexaPro-foldon constructs used for expression and stability comparisons with and without Rpk9 mutations contained a BM40 signal peptide and were placed into pCMV/R. Plasmids were transformed into the NEB 5α strain of *E. coli* (New England Biolabs) for subsequent DNA extraction from bacterial culture (NucleoBond Xtra Midi kit) to obtain plasmid for transient transfection into Expi293F cells. The amino acid sequences of all novel proteins used in this study can be found in **Supplementary Information 2**.

### Transient transfection

SARS-CoV-2 S and ACE2-Fc proteins were produced in Expi293F cells grown in suspension using Expi293F expression medium (Life Technologies) at 33°C, 70% humidity, 8% CO_2_, rotating at 150 rpm. The cultures were transfected using PEI-MAX (Polyscience) with cells grown to a density of 3.0 million cells per mL and cultivated for 3 days. Supernatants were clarified by centrifugation (5 min at 4000 rcf), addition of PDADMAC solution to a final concentration of 0.0375% (Sigma Aldrich, #409014), and a second centrifugation step (5 min at 4000 rcf).

Genes encoding CV30 and CR3022 heavy and light chains were ordered from GenScript and cloned into pCMV/R. Antibodies were expressed by transient co-transfection of both heavy and light chain plasmids in Expi293F cells using PEI MAX (Polyscience) transfection reagent. Cell supernatants were harvested and clarified after 3 or 6 days as described above.

### Purification of glycoproteins

Proteins containing His tags were purified from clarified supernatants via a batch bind method where each clarified supernatant was supplemented with 1 M Tris-HCl pH 8.0 to a final concentration of 45 mM and 5 M NaCl to a final concentration of 313 mM. Talon cobalt affinity resin (Takara) was added to the treated supernatants and allowed to incubate for 15 min with gentle shaking. Resin was collected using vacuum filtration with a 0.45 μm filter and transferred to a gravity column. The resin was washed with 10 column volumes of 20 mM Tris pH 8.0, 300 mM NaCl, and bound protein was eluted with 3 column volumes of 20 mM Tris pH 8.0, 300 mM NaCl, 300 mM imidazole. The batch bind process was then repeated on the same supernatant sample and the first and second elutions were combined. SDS-PAGE was used to assess purity. For quantification of yields of RBD-based constructs, IMAC elutions from comparable cell culture conditions and volumes were supplemented with 100 mM L-arginine and 5% glycerol and concentrated to 1.5 mL. The concentrated samples were subsequently loaded into a 1 mL loop and applied to a Superdex 75 Increase 10/300 GL column (for monomeric RBDs) or a Superdex 200 Increase 10/300 GL column (for RBD fusions to the I53-50A trimer) pre-equilibrated with 50 mM Tris pH 8, 150 mM NaCl, 100 mM L-arginine, 5% glycerol. For quantification of yields of HexaPro-foldon constructs with and without Rpk9 mutations, IMAC elutions from comparable cell culture conditions and volumes were supplemented with 5% glycerol and concentrated to 1.5 mL, which was subsequently loaded into a 1 mL loop and applied to a Superose 6 Increase 10/300 GL column pre-equilibrated with 50 mM Tris pH 8.0, 150 mM NaCl, 0.25% w/v L-histidine, 5% glycerol. HexaPro-foldon for immunization studies was purified by IMAC and dialyzed three times against 50 mM Tris pH 8.0, 150 mM NaCl, 0.25% w/v L-histidine, 5% glycerol for four hours at room temperature.

### Thermal denaturation (nanoDSF)

RBD-based samples were prepared in a buffer containing 50 mM Tris pH 8, 150 mM NaCl, 100 mM L-arginine, 5% glycerol at 1.0 mg/mL for nanoDSF analysis. HexaPro-foldon-based samples were initially prepared in a buffer containing 50 mM Tris pH 8.0, 150 mM NaCl, 0.25% w/v L-histidine, 5% glycerol at 1.1 mg/mL. Prior to nanoDSF, 9 volumes of HexaPro-foldon-based protein was mixed with 1 volume of 50 mM Tris pH 8.0, 150 mM NaCl, 0.25% w/v L-histidine, 5% glycerol, 0.04% Tween-20 that either was or was not further supplemented with 2.9 mM linoleic acid (Sigma), which were lightly shaken at room temperature for 2 hr prior to nanoDSF analysis. Non-equilibrium melting temperatures were determined using an UNcle (UNchained Labs) based on the barycentric mean of intrinsic tryptophan fluorescence emission spectra collected from 20–95°C using a thermal ramp of 1°C per minute. Melting temperatures were defined as the maximum point of the first derivative of the melting curve, with first derivatives calculated using GraphPad Prism software after smoothing with four neighboring points using 2nd order polynomial settings.

### Circular dichroism

Far-ultraviolet CD measurements for monomeric RBD variants were performed on a JASCO-1500 equipped with a temperature-controlled six-cell holder. Wavelength scans were measured from 260–190 nm at 25, 95°C, and again at 25°C after fast refolding (~5 min), on 0.4 mg/mL protein in 25 mM Tris pH 8.0, 150 mM NaCl, 5% glycerol using a 1 mm path-length cuvette.

### SYPRO Orange fluorescence

5000× SYPRO Orange Protein Gel Stain (Thermo Fisher) was diluted into 25 mM Tris pH 8.0, 150 mM NaCl, 5% glycerol and further added to monomeric RBDs prepared in the same buffer, with final concentrations of 1.0 mg/mL for the RBDs and 20× for SYPRO Orange. Samples were loaded into an UNcle Nano-DSF (UNChained Laboratories) and fluorescence emission spectra were collected for all samples 5 min after the addition of SYPRO Orange to the samples.

### Microbial protein expression and purification of I53-50B.4PT1

The complementary pentameric nanoparticle component to RBD-I53-50A, I53-50B.4PT1, was produced as previously described (61), and the same protocol was used for purification of the 2OBX non-assembling control pentamer.

### *In vitro* nanoparticle assembly

Total protein concentration of purified individual nanoparticle components was determined by measuring absorbance at 280 nm using a UV/vis spectrophotometer (Agilent Cary 8454) and calculated extinction coefficients (62). The assembly steps were performed at room temperature with addition in the following order: wild-type or stabilized RBD-I53-50A trimeric fusion protein, followed by q.s. with buffer as needed to achieve desired final concentration, and finally I53-50B.4PT1 pentameric component (in 50 mM Tris pH 8, 500 mM NaCl, 0.75% w/v CHAPS), with a molar ratio of RBD-I53-50A:I53-50B.4PT1 of 1.1:1. q.s. buffer either contained 50 mM Tris pH 7.4, 185 mM NaCl, 100 mM L-arginine, 0.75% CHAPS, 4.5% glycerol (for solution stability studies) or 50 mM Tris pH 8, 150 mM NaCl, 100 mM L-arginine, 5% glycerol. All RBD-I53-50 *in vitro* assemblies were incubated at 2-8°C with gentle rocking for at least 30 min before subsequent purification by SEC in order to remove residual unassembled component. A Superose 6 Increase 10/300 GL column was used for nanoparticle purification. Assembled nanoparticles elute at ~11 mL on the Superose 6 column. Assembled nanoparticles were sterile filtered (0.22 μm) immediately prior to column application and following pooling of SEC fractions.

### Bio-layer interferometry for kinetic analysis of monomeric RBDs

Kinetic measurements were performed using an Octet Red 96 System (Pall FortéBio/Sartorius) at 25°C with shaking at 1000 rpm. All proteins and antibodies were diluted into phosphate buffered saline (PBS) with 0.5% bovine serum albumin and 0.01% Tween-20 in black 96-well Greiner Bio-one microplate at 200 μL per well, with the buffer alone used for baseline and dissociation steps. CV30 or CR3022 IgG at 10ug/mL was loaded onto pre-hydrated Protein A biosensors (Pall FortéBio/Sartorius) for 150 s followed by a 60 s baseline. Biosensors were then transferred into an association step with one of five serial two-fold dilutions of wild-type or stabilized monomeric RBDs for 120 s, with RBD concentrations of 125 μM, 62.5 μM, 31.3 μM, 15.6 μM and 7.8 μM used for CV30 and 31.3 μM, 15.6 μM, 7.8 μM, 3.9 μM and 2.0 μM used for CR3022. After association, biosensors were transferred into buffer for 300 s of dissociation. Data from the association and dissociation steps were baseline subtracted and kinetics measurements were calculated globally across all five serial dilutions of RBD using a 1:1 binding model (FortéBio analysis software, version 12.0).

### Bio-layer interferometry for fractional antigenicity of RBD nanoparticles

Binding of hACE2-Fc (dimerized receptor) and CR3022 IgG to monomeric RBD and RBD-I53-50 nanoparticles was analyzed for real-time stability studies using an Octet Red 96 System (Pall FortéBio/Sartorius) at ambient temperature with shaking at 1000 rpm. Protein samples were diluted to 100 nM in Kinetics buffer (Pall FortéBio/Sartorius). Buffer, antibody, receptor, and immunogen were then applied to a black 96-well Greiner Bio-one microplate at 200 μL per well. Protein A biosensors were first hydrated for 10 min in Kinetics buffer, then dipped into either hACE2-Fc or CR3022 diluted to 10 μg/mL in Kinetics buffer in the immobilization step. After 500 s, the tips were transferred to Kinetics buffer for 90 s to reach a baseline. The association step was performed by dipping the loaded biosensors into the immunogens for 300 s, and the subsequent dissociation steps was performed by dipping the biosensors back into Kinetics buffer for an additional 300 s. The data were baseline subtracted prior to plotting using the FortéBio analysis software (version 12.0).

### Negative stain electron microscopy

Wild-type RBD-I53-50 nanoparticles, Rpk4-I53-50 nanoparticles and Rpk9-I53-50 nanoparticles were first diluted to 75 μg/mL in 50 mM Tris pH 8, 150 mM NaCl, 100 mM L-arginine, 5% v/v glycerol prior to application of 3 μL of sample onto freshly glow-discharged 300-mesh copper grids. Sample was incubated on the grid for 1 minute before the grid was dipped in a 50 μL droplet of water and excess liquid blotted away with filter paper (Whatman). The grids were then dipped into 6 μL of 0.75% w/v uranyl formate stain. Stain was blotted off with filter paper, then the grids were dipped into another 6 μL of stain and incubated for ~90 seconds. Finally, the stain was blotted away and the grids were allowed to dry for 1 minute prior to storage or imaging. Prepared grids were imaged in a Talos model L120C transmission electron microscope using a Gatan camera at 57,000×.

### Dynamic light scattering

Dynamic light scattering (DLS) was used to measure hydrodynamic diameter (Dh) and polydispersity (%Pd) of RBD-I53-50 nanoparticle samples on an UNcle (UNchained Laboratories). Sample was applied to an 8.8 μL quartz capillary cassette (UNi, UNchained Laboratories) and measured with 10 acquisitions of 5 s each, using auto-attenuation of the laser. Increased viscosity due to 5% v/v glycerol in the RBD nanoparticle buffer was accounted for by the UNcle Client software in Dh measurements.

### Endotoxin measurements

Endotoxin levels in protein samples were measured using the EndoSafe Nexgen-MCS System (Charles River). Samples were diluted 1:50 or 1:100 in Endotoxin-free LAL reagent water, and applied into wells of an EndoSafe LAL reagent cartridge. Charles River EndoScan-V software was used to analyze endotoxin content, which automatically back-calculates for the dilution factor. Endotoxin values were reported as EU/mL which were then converted to EU/mg based on UV-Vis measurements. Our threshold for samples suitable for immunization was <100 EU/mg.

### UV-Vis

Ultraviolet-visible spectrophotometry (UV-Vis) measurements were taken using an Agilent Technologies Cary 8454. Samples were applied to a 10 mm, 50 μL quartz cell (Starna Cells, Inc.) and absorbance was measured from 180 to 1000 nm. Net absorbance at 280 nm, obtained from measurement and single reference wavelength baseline subtraction, was used with calculated extinction coefficients and molecular weights to obtain protein concentration. The ratio of absorbance at 320/280 nm was used to determine relative aggregation levels in real-time stability study samples. Samples were diluted with respective blanking buffers to obtain an absorbance between 0.1 and 1.0. All data produced from the UV/vis instrument was processed in the 845x UV/visible System software.

### Hydrogen/deuterium-exchange mass spectrometry

3 mg of RBD-I53-50A, Rpk4-I53-50A, and Rpk9-I53-50A trimers were H/D exchanged (HDX) in the deuteration buffer (pH* 7.5, 85% D_2_O, Cambridge Isotope Laboratories, Inc.) for 3, 15, 60, 1800, and 72000 seconds at 22°C respectively. Exchanged samples were subsequently mixed 1:1 with ice-chilled quench buffer (200 mM tris(2-chlorethyl) phosphate (TCEP), 8 M Urea, 0.2% formic acid (FA)) for a final pH of 2.5 and immediately flash frozen in liquid nitrogen. Samples were analyzed by LC-MS on a Waters Synapt G2-Si mass spectrometer using a custom-built loading system that maintained all columns, loops, valves, and lines at 0°C. Frozen samples were thawed on ice and loaded over a custom packed immobilized pepsin column (2.1 × 50 mm) with a 200 mL/min flow of 0.1% trifluoroacetic acid (TFA) with 2% acetonitrile. Peptides were trapped on a Waters CSH C18 trap cartridge (2.1 × 5 mm) prior to being resolved over a Waters CSH C18 1 × 100 mm 1.7 μm column with a linear gradient from 3% to 40% B over 18 min (A: 98% water, 2% acetonitrile, 0.1% FA, 0.025% TFA; B: 100% acetonitrile, 0.1% FA, flow rate of 40 mL/min). A series of washes were performed between sample runs to minimize carryover (63). All the water and organic solvent used, unless specifically stated, were MS grade (Optima™, Fisher). A fully deuterated control for each sample series was made by LC eluate collection from pepsin-digested undeuterated sample, speedvac drying, incubation in deuteration buffer for 1 hour at 85°C, and quenching the same as all other HDX samples. Internal exchange standards (Pro-Pro-Pro-Ile [PPPI] and Pro-Pro-Pro-Phe [PPPF]) were added in each sample to ensure consistent labeling conditions for all samples (64).

The peptide reference list was updated from wild-type RBD peptide list with addition of new peptides covering mutations (25). These peptides were manually validated using DriftScope™ (Waters) and identified with orthogonal retention time and drift time coordinates. Deuterium uptake analysis was performed with HX-Express v2 (65, 66). Peaks were identified from the peptide spectra based on the peptide m/z values and binomial fitting was applied. The deuterium uptake level was normalized relative to fully deuterated controls.

### Mouse immunizations

Female BALB/c (Stock: 000651) mice were purchased at the age of four weeks from The Jackson Laboratory, Bar Harbor, Maine, and maintained at the Comparative Medicine Facility at the University of Washington, Seattle, WA, accredited by the American Association for the Accreditation of Laboratory Animal Care International (AAALAC). At six weeks of age, 6 mice per dosing group were vaccinated with a prime immunization, and three weeks later mice were boosted with a second vaccination. Prior to inoculation, immunogen suspensions were gently mixed 1:1 vol/vol with AddaVax adjuvant (Invivogen, San Diego, CA) to reach a final concentration of 0.009 or 0.05 mg/mL antigen. Mice were injected intramuscularly into the gastrocnemius muscle of each hind leg using a 27-gauge needle (BD, San Diego, CA) with 50 μL per injection site (100 μL total) of immunogen under isoflurane anesthesia. To obtain sera all mice were bled two weeks after prime and boost immunizations. Blood was collected via submental venous puncture and rested in 1.5 mL plastic Eppendorf tubes at room temperature for 30 min to allow for coagulation. Serum was separated from hematocrit via centrifugation at 2,000 g for 10 min. Complement factors and pathogens in isolated serum were heat-inactivated via incubation at 56°C for 60 min. Serum was stored at 4°C or −80°C until use. All experiments were conducted at the University of Washington, Seattle, WA according to approved Institutional Animal Care and Use Committee protocols.

### ELISA

Enzyme-linked immunosorbent assays (ELISA) were used to determine the binding of mouse sera to the delivered antigens. In brief, Maxisorp (Nunc) ELISA plates were coated overnight at 4°C with 0.08 μg/mL of protein of interest per well in 0.1 M sodium bicarbonate buffer, pH 9.4. Plates were then blocked with a 4% (w/v) solution of dried milk powder (BioRad) in TBS with 0.05% (v/v) Tween 20 (TBST) for 1 hour at room temperature. Serial dilutions of sera were added to the plates and, after washing, antibody binding was revealed using a hydrogen peroxidase coupled horse anti-mouse IgG antibody. Plates were then washed thoroughly in TBST, colorimetric substrate (TMB, Thermo Fisher) was added and absorbance was read at 450 nm. Area under curve (AUC) calculations were generated by addition of trapezoidal areas generated between adjacent pairs of absorbance measurements and baseline. Midpoint titers calculations (EC_50_) were generated based on fitted four point logistic equations using the SciPy library in Python, in which the EC_50_ was the serum dilution at which the curve reached 50% of its maximum.

### Lentivirus-based pseudovirus neutralization assays

Spike-pseudotyped lentivirus neutralization assays were carried out essentially as described in (67). The protocol was modified for this study to use a SARS-CoV-2 Spike with a 21 amino-acid cytoplasmic tail truncation, which increases Spike-pseudotyped lentivirus titers (68), and the D614G mutation, which is now predominant in human SARS-CoV-2 (69). The plasmid encoding this Spike, HDM_Spikedelta21_D614G, is available from Addgene (#158762) or BEI (NR-53765), and the full sequence is at (https://www.addgene.org/158762).

Briefly, 293T-ACE2 cells (BEI NR-52511) were seeded at 1.25×10^4^ cells per well in 50 uL D10 growth media (DMEM with 10% heat-inactivated FBS, 2 mM L-glutamine, 100 U/mL penicillin, and 100 μg/mL streptomycin) in poly-L-lysine coated black-walled clear-bottom 96-well plates (Greiner 655930). The next day, mouse serum samples were heat inactivated for 30 min at 56°C and then serially diluted in D10 growth media. Spike-pseudotyped lentivirus was diluted 1:50 to yield ~200,000 RLUs per well and incubated with the serum dilutions for 1 hr at 37°C. 100 μL of virus-serum mixture was then added to the cells and ~52 hours later luciferase activity was measured using the Bright-Glo Luciferase Assay System (Promega, E2610). Each batch of neutralization assays included a negative control sample of human serum collected in 2017-2018 and a known neutralizing antibody to ensure consistency between batches. Fraction infectivity for each well was calculated compared to two “no-serum” control wells in the same row of the plate. We used the “neutcurve” package (https://jbloomlab.github.io/neutcurve version 0.5.2) to calculate the inhibitory concentration 50% (IC_50_) and the neutralization titer 50% (NT50), which is simply 1/IC_50_, for each serum sample by fitting a Hill curve with the bottom fixed at 0 and the top fixed at 1. All neutralization assay data are available at https://github.com/jbloomlab/RBD_nanoparticle_vaccine.

### MLV-based pseudovirus neutralization assays

MLV-based SARS-CoV-2 S pseudotypes were prepared as previously described (10, 25, 50, 70). Briefly, HEK293T cells were co-transfected using Lipofectamine 2000 (Life Technologies) with an SARS-CoV-2 S-encoding plasmid, an MLV Gag-Pol packaging construct, and the MLV transfer vector encoding a luciferase reporter according to the manufacturer’s instructions. Cells were washed 3× with Opti-MEM and incubated for 5 h at 37°C with transfection medium. DMEM containing 10% FBS was added for 60 h. The supernatants were harvested by spinning at 2,500 g, filtered through a 0.45 μm filter, concentrated with a 100 kDa membrane for 10 min at 2,500 g and then aliquoted and stored at −80°C.

For neutralization assays, HEK-hACE2 cells were cultured in DMEM with 10% FBS (Hyclone) and 1% PenStrep with 8% CO_2_ in a 37°C incubator (ThermoFisher). One day or more prior to infection, 40 μL of poly-lysine (Sigma) was placed into 96-well plates and incubated with rotation for 5 min. Poly-lysine was removed, plates were dried for 5 min then washed 1× with water prior to plating cells HEK-hACE2 cells. The following day, cells were checked to be at 80% confluence. In a half-area 96-well plate, a 1:3 serial dilution of sera was made in DMEM in 22 μL final volume. 22 μL of pseudovirus was then added to the serial dilution and incubated at room temperature for 30-60 min. The mixture was added to cells and 2 hours later 44 μL of DMEM supplemented with 20% FBS and 2% PenStrep was added and cells were incubated for 48 hours. After 48 h 40 μL/well of One-Glo-EX substrate (Promega) was added to the cells and incubated in the dark for 5-10 min prior to reading on a BioTek plate reader. Nonlinear regression of log(inhibitor) versus normalized response was used to determine IC_50_ values from curve fits.

### Quantification and Statistical Analysis

Multi-group comparisons were performed using nonparametric Kruskal-Wallis test with Dunn’s post-hoc analysis in GraphPad Prism 8. Differences were considered significant when P values were less than 0.05. Statistical methods and P value ranges can be found in the Figures and Figure legends.

## DATA AVAILABILITY STATEMENT

The raw data supporting the conclusions of this article will be made available by the authors, without undue reservation.

## ACKNOWLEDGMENTS

We thank Ratika Krishnamurty for program management. This study was supported by the Bill & Melinda Gates Foundation (OPP1156262 to D.V. and N.P.K., and OPP1126258 to K.K.L.), a generous gift from the Audacious Project, a generous gift from the Open Philanthropy Project, a generous gift from Jodi Green and Mike Halperin, a generous gift from the Hanauer family, the Defense Threat Reduction Agency (HDTRA1-18-1-0001 to N.P.K.), the National Institute of Allergy and Infectious Diseases (AI141707 to J.D.B., AI149928 to K.H.D.C, DP1AI158186 and HHSN272201700059C to D.V.), a Pew Biomedical Scholars Award (D.V.), Investigators in the Pathogenesis of Infectious Disease Awards from the Burroughs Wellcome Fund (D.V.) and Fast Grants (D.V.). T.N.S. is an HHMI Fellow of the Damon Runyon Cancer Research Foundation. J.D.B. is an Investigator of the Howard Hughes Medical Institute.

## AUTHOR CONTRIBUTIONS

Conceptualization, D.E. and N.P.K.; Modeling and Design, D.E.; Formal Analysis, D.E., N.B., K.H.D.C., A.C.W., C.C., K.L.H., B.F., J.C.K., L.C., T.S., A.G., K.K.L., D.V., J.B. and N.P.K.; Investigation, D.E., N.B., K.H.D.C., A.C.W., M.N.P., C.C., K.L.H., B.F., M.M., D.P., J.C.K., K.D.M., M.J.N., C.O., E.K., C.S., M.A., M.J., A.B., L.C., T.N.S., A.J.G., K.K.L., D.V., J.B. and N.P.K.; Resources, R.R.; Writing – Original Draft, D.E., N.B., C.C. and B.F.; Writing – Review & Editing, all authors; Visualization, D.E., N.B., C.C., B.F. and J.C.K.; Supervision, L.C., K.K.L., D.V., J.B. and N.P.K.; Funding Acquisition, D.V., J.B. and N.P.K.

## CORRESPONDENCE

Correspondence to N.P.K.

