## Supplementary Information 1 for "Stabilization of the SARS-CoV-2 Spike receptor-binding domain using deep mutational scanning and structure-based design"

```
<ROSETTASCRIPTS>
      <SCOREFXNS>
            <ScoreFunction name="sfx_clean" weights="beta" symmetric="1"
/>
      </SCOREFXNS>

      <TASKOPERATIONS>
                 <ReadResfile name="designed_identities"
filename="%%resfile%%" />
      </TASKOPERATIONS>

      <RESIDUE_SELECTORS>
            <Task name="resfile_muts" designable="true"
task_operations="designed_identities" /> // selects mutations from
resfile
                  <Not name="not_resfile" selector="resfile_muts" /> //
everything else that isn't mutated

            <Neighborhood name="mutant_neighbor" selector="resfile_muts"
distance="10" include_focus_in_subset="false" /> // only operate on other
residues that are 10A from mutation
            <Or name="resfile_or_nbors"
selectors="resfile_muts,mutant_neighbor" />
                  <Not name="not_res_or_nbors" selector="resfile_or_nbors"
/>
            <ResidueName name="cysteines" residue_name3="CYS" /> // will
be used to make sure that disulfides aren't repacked or minimized

      </RESIDUE_SELECTORS>

      <TASKOPERATIONS>
            <IncludeCurrent name="ic" /> // includes input pdb's rotamers
in packing operations
            <LimitAromaChi2 name="limitaro" chi2max="110" chi2min="70" />
      // disallow extreme aromatic rotamers
            <RestrictToRepacking name="repack_only" />                //
for minimize/repack
            <ExtraRotamersGeneric name="ex1_ex2" ex1="1" ex2="1" />
      // use ex1 ex2 rotamers

            <OperateOnResidueSubset name="repack_neighbor"
selector="mutant_neighbor" > // neighbors to mutations are only allowed
to repack
                  <RestrictToRepackingRLT/> </OperateOnResidueSubset>

            <OperateOnResidueSubset name="lock_not_resfile_or_nbors"
selector="not_res_or_nbors" > // everything that isn't a mutation or
neighbor cannot move
                         <PreventRepackingRLT/> </OperateOnResidueSubset>

            <OperateOnResidueSubset name="lock_cysteines"
selector="cysteines" > // freeze cysteines
                         <PreventRepackingRLT/> </OperateOnResidueSubset>
```

```
      </TASKOPERATIONS>

      <MOVERS>
            <Symmetrizer name="gen_docked_config" symm_file="%%symfile%%"
/>
            <SymPackRotamersMover name="design_from_resfile"
scorefxn="sfx_clean"
task_operations="designed_identities,repack_neighbor,lock_not_resfile_or_
nbors,limitaro,ic,ex1_ex2" /> // Adds mutation and repacks the mutation
and neighbors
            <TaskAwareSymMinMover name="rb_min_hard_bb"
scorefxn="sfx_clean" bb="%%bb%%" chi="1" rb="0"
task_operations="repack_neighbor,lock_not_resfile_or_nbors" /> //
minimizes sidechains, and also backbone if bb set to 1
            <SymPackRotamersMover name="repack_hard" scorefxn="sfx_clean"
task_operations="repack_only,repack_neighbor,lock_not_resfile_or_nbors,li
mitaro,ic,ex1_ex2" /> // repacks mutation and neighbors only

      </MOVERS>

      <FILTERS>
                 <SaveResfileToDisk name="save_resfile"
task_operations="designed_identities" designable_only="0"
resfile_prefix="%%outpath%%" resfile_suffix="" resfile_name=""
resfile_general_property="NATRO" selected_resis_property=""
renumber_pdb="0" /> // save a resfile with mutations. Only gives useful
info if allowing Rosetta to make decisions

      </FILTERS>

      <PROTOCOLS>
            // generate docked configuration
            <Add mover_name="gen_docked_config" />

            // design and minimization
            <Add mover_name="design_from_resfile" />
            <Add mover_name="rb_min_hard_bb" />
            <Add mover_name="design_from_resfile" />
            <Add mover_name="rb_min_hard_bb" />

            // save resfile
            <Add filter_name="save_resfile" />

      </PROTOCOLS>

</ROSETTASCRIPTS>
```
