## Supplementary Information 2 for "Stabilization of the SARS-CoV-2 Spike receptor-binding domain using deep mutational scanning and structure-based design"

>RBD-I53-50A trimer (16-GS linker, using wild type RBD from Wuhan-Hu-1)
MGILPSPGMPALLSLVSLLSVLLMGCVAETGTRFPNITNLCPFGEVFNATRFASVYAWNRKRISNCVADY
SVLYNSASFSTFKCYGVSPTKLNDLCFTNVYADSFVIRGDEVRQIAPGQTGKIADYNYKLPDDFTGCVIA
WNSNNLDSKVGGNYNYLYRLFRKSNLKPFERDISTEIYQAGSTPCNGVEGFNCYFPLQSYGFQPTNGVGY
QPYRVVVLSFELLHAPATVCGPKKSTGGSGGSGSGGSGGSGSEKAAKAEEAARKMEELFKKHKIVAVLRA
NSVEEAIEKAVAVFAGGVHLIEITFTVPDADTVIKALSVLKEKGAIIGAGTVTSVEQARKAVESGAEFIV
SPHLDEEISQFAKEKGVFYMPGVMTPTELVKAMKLGHTILKLFPGEVVGPQFVKAMKGPFPNVKFVPTGG
VNLDNVAEWFKAGVLAVGVGSALVKGTPDEVREKAKAFVEKIRGATEGGSHHHHHHHH

>Rpk1-I53-50A trimer
MGILPSPGMPALLSLVSLLSVLLMGCVAETGTRFPNITNLCPFGEVFNATRFASVYAWNRKRISNCVADW
SVLYNSASFSTFKCYGVSPTKLNDLCFTNVYADSFVIRGDEVRQIAPGQTGKIADYNYKLPDDFTGCVIA
WNSNNLDSKVGGNYNYLYRLFRKSNLKPFERDISTEIYQAGSTPCNGVEGFNCYFPLQSYGFQPTNGVGY
QPYRVVVLSFELLHAPATVCGPKKSTGGSGGSGSGGSGGSGSEKAAKAEEAARKMEELFKKHKIVAVLRA
NSVEEAIEKAVAVFAGGVHLIEITFTVPDADTVIKALSVLKEKGAIIGAGTVTSVEQARKAVESGAEFIV
SPHLDEEISQFAKEKGVFYMPGVMTPTELVKAMKLGHTILKLFPGEVVGPQFVKAMKGPFPNVKFVPTGG
VNLDNVAEWFKAGVLAVGVGSALVKGTPDEVREKAKAFVEKIRGATEGGSHHHHHHHH

>Rpk2-I53-50A trimer
MGILPSPGMPALLSLVSLLSVLLMGCVAETGTRFPNITNLCPLGEVFNATRFASVYAWNRKRISNCVADW
SVLYNSASFSTFKCYGVSPTKLNDLCFTNVYADSFVIRGDEVRQIAPGQTGKIADYNYKLPDDFTGCVIA
WNSNNLDSKVGGNYNYLYRLFRKSNLKPFERDISTEIYQAGSTPCNGVEGFNCYFPLQSYGFQPTNGVGY
QPYRVVVLSFELLHAPATVCGPKKSTGGSGGSGSGGSGGSGSEKAAKAEEAARKMEELFKKHKIVAVLRA
NSVEEAIEKAVAVFAGGVHLIEITFTVPDADTVIKALSVLKEKGAIIGAGTVTSVEQARKAVESGAEFIV
SPHLDEEISQFAKEKGVFYMPGVMTPTELVKAMKLGHTILKLFPGEVVGPQFVKAMKGPFPNVKFVPTGG
VNLDNVAEWFKAGVLAVGVGSALVKGTPDEVREKAKAFVEKIRGATEGGSHHHHHHHH

>Rpk3-I53-50A trimer
MGILPSPGMPALLSLVSLLSVLLMGCVAETGTRFPNITNLCPFGEVFNATRFASVYAWNRKRISNCVADW
SVLYNSASFSTFKCYGVSPTKLNDLCFTNVYADSFVIRGDEVRQIAPGQTGKIADYNYKLPDDFTGCVIA
WNSNNLDSKVGGNYNYLYRLFRKSNLKPFERDISTEIYQAGSTPCNGVEGFNCYFPLQSYGFQPTNGVGY
QPYRVVVMSFELLHAPATVCGPKKSTGGSGGSGSGGSGGSGSEKAAKAEEAARKMEELFKKHKIVAVLRA
NSVEEAIEKAVAVFAGGVHLIEITFTVPDADTVIKALSVLKEKGAIIGAGTVTSVEQARKAVESGAEFIV
SPHLDEEISQFAKEKGVFYMPGVMTPTELVKAMKLGHTILKLFPGEVVGPQFVKAMKGPFPNVKFVPTGG
VNLDNVAEWFKAGVLAVGVGSALVKGTPDEVREKAKAFVEKIRGATEGGSHHHHHHHH

>Rpk4-I53-50A trimer

MGILPSPGMPALLSLVSLLSVLLMGCVAETGTRFPNITNLCPFGEVFNATRFASVYAWNRKRISNCVADY
SVLYNSASFSTFKCYGVSPTKLNDLCWTNVYADSFVIRGDEVRQIAPGQTGKIADYNYKLPDDFTGCVIA
WNSNNLDSKVGGNYNYLYRLFRKSNLKPFERDISTEIYQAGSTPCNGVEGFNCYFPLQSYGFQPTNGVGY
QPYRVVVLSFELLHAPATVCGPKKSTGGSGGSGSGGSGGSGSEKAAKAEEAARKMEELFKKHKIVAVLRA
NSVEEAIEKAVAVFAGGVHLIEITFTVPDADTVIKALSVLKEKGAIIGAGTVTSVEQARKAVESGAEFIV
SPHLDEEISQFAKEKGVFYMPGVMTPTELVKAMKLGHTILKLFPGEVVGPQFVKAMKGPFPNVKFVPTGG
VNLDNVAEWFKAGVLAVGVGSALVKGTPDEVREKAKAFVEKIRGATEGGSHHHHHHHH

>Rpk5-I53-50A trimer
MGILPSPGMPALLSLVSLLSVLLMGCVAETGTRFPNITNLCPFGEVFNATRFASVYAWNRKRISNCVADW
SVLYNSASFSTFKCYGVSPTKLNDLCWTNVYADSFVIRGDEVRQIAPGQTGKIADYNYKLPDDFTGCVIA
WNSNNLDSKVGGNYNYLYRLFRKSNLKPFERDISTEIYQAGSTPCNGVEGFNCYFPLQSYGFQPTNGVGY
QPYRVVVLSFELLHAPATVCGPKKSTGGSGGSGSGGSGGSGSEKAAKAEEAARKMEELFKKHKIVAVLRA
NSVEEAIEKAVAVFAGGVHLIEITFTVPDADTVIKALSVLKEKGAIIGAGTVTSVEQARKAVESGAEFIV
SPHLDEEISQFAKEKGVFYMPGVMTPTELVKAMKLGHTILKLFPGEVVGPQFVKAMKGPFPNVKFVPTGG
VNLDNVAEWFKAGVLAVGVGSALVKGTPDEVREKAKAFVEKIRGATEGGSHHHHHHHH

>Rpk6-I53-50A trimer
MGILPSPGMPALLSLVSLLSVLLMGCVAETGTRFPNITNLCPMGEVFNATRFASVYAWNRKRISNCVLDF
SVLYNSASFSTVKCYGVSPTKLNDLCFTNVYADSFVIRGDEVRQIAPGQTGKIADYNYKLPDDFTGCVIA
WNSNNLDSKVGGNYNYLYRLFRKSNLKPFERDISTEIYQAGSTPCNGVEGFNCYFPLQSYGFQPTNGVGY
QPYRVVVLSFELLHAPATVCGPKKSTGGSGGSGSGGSGGSGSEKAAKAEEAARKMEELFKKHKIVAVLRA
NSVEEAIEKAVAVFAGGVHLIEITFTVPDADTVIKALSVLKEKGAIIGAGTVTSVEQARKAVESGAEFIV
SPHLDEEISQFAKEKGVFYMPGVMTPTELVKAMKLGHTILKLFPGEVVGPQFVKAMKGPFPNVKFVPTGG
VNLDNVAEWFKAGVLAVGVGSALVKGTPDEVREKAKAFVEKIRGATEGGSHHHHHHHH

>Rpk7-I53-50A trimer
MGILPSPGMPALLSLVSLLSVLLMGCVAETGTRFPNITNLCPFGEVFNATRFASVYAWNRKRISNCVADF
SVLYNSASFSTFKCYGVSPTKLNDLCWTNVYADSFVIRGDEVRQIAPGQTGKIADYNYKLPDDFTGCVIA
WNSNNLDSKVGGNYNYLYRLFRKSNLKPFERDISTEIYQAGSTPCNGVEGFNCYFPLQSYGFQPTNGVGY
QPYRVVVLSFELLHAPATVCGPKKSTGGSGGSGSGGSGGSGSEKAAKAEEAARKMEELFKKHKIVAVLRA
NSVEEAIEKAVAVFAGGVHLIEITFTVPDADTVIKALSVLKEKGAIIGAGTVTSVEQARKAVESGAEFIV
SPHLDEEISQFAKEKGVFYMPGVMTPTELVKAMKLGHTILKLFPGEVVGPQFVKAMKGPFPNVKFVPTGG
VNLDNVAEWFKAGVLAVGVGSALVKGTPDEVREKAKAFVEKIRGATEGGSHHHHHHHH

>Rpk8-I53-50A trimer
MGILPSPGMPALLSLVSLLSVLLMGCVAETGTRFPNITNLCPFGEVFNATRFASVYAWNRKRISNCVADF
SVLYNSASFSTFKCYGVSPTKLNDLCFTNIYADSFVIRGDEVRQIAPGQTGKIADYNYKLPDDFTGCVIA

WNSNNLDSKVGGNYNYLYRLFRKSNLKPFERDISTEIYQAGSTPCNGVEGFNCYFPLQSYGFQPTNGVGY
QPYRVVVLSFELLHAPATVCGPKKSTGGSGGSGSGGSGGSGSEKAAKAEEAARKMEELFKKHKIVAVLRA
NSVEEAIEKAVAVFAGGVHLIEITFTVPDADTVIKALSVLKEKGAIIGAGTVTSVEQARKAVESGAEFIV
SPHLDEEISQFAKEKGVFYMPGVMTPTELVKAMKLGHTILKLFPGEVVGPQFVKAMKGPFPNVKFVPTGG
VNLDNVAEWFKAGVLAVGVGSALVKGTPDEVREKAKAFVEKIRGATEGGSHHHHHHHH

>Rpk9-I53-50A trimer
MGILPSPGMPALLSLVSLLSVLLMGCVAETGTRFPNITNLCPFGEVFNATRFASVYAWNRKRISNCVADF
SVLYNSASFSTFKCYGVSPTKLNDLCWTNIYADSFVIRGDEVRQIAPGQTGKIADYNYKLPDDFTGCVIA
WNSNNLDSKVGGNYNYLYRLFRKSNLKPFERDISTEIYQAGSTPCNGVEGFNCYFPLQSYGFQPTNGVGY
QPYRVVVLSFELLHAPATVCGPKKSTGGSGGSGSGGSGGSGSEKAAKAEEAARKMEELFKKHKIVAVLRA
NSVEEAIEKAVAVFAGGVHLIEITFTVPDADTVIKALSVLKEKGAIIGAGTVTSVEQARKAVESGAEFIV
SPHLDEEISQFAKEKGVFYMPGVMTPTELVKAMKLGHTILKLFPGEVVGPQFVKAMKGPFPNVKFVPTGG
VNLDNVAEWFKAGVLAVGVGSALVKGTPDEVREKAKAFVEKIRGATEGGSHHHHHHHH

>Rpk10-I53-50A trimer
MGILPSPGMPALLSLVSLLSVLLMGCVAETGTRFPNITNLCPFGEVFNATRFASVYAWNRKRISNCVADW
SVLYNSASFSTFKCYGVSPTKLNDLCFTNVYADSFVIRGDEVRQIAPGQTGKIADYNYKLPDDFTGCVIA
WNSNNLDSKVGGNYNYLYRLFRKSNLKPFERDISTEIYQAGSTPCNGVEGFNCYFPLQSYGFQPTNGVGY
QPYRVVVISLELLHAPATVCGPKKSTGGSGGSGSGGSGGSGSEKAAKAEEAARKMEELFKKHKIVAVLRA
NSVEEAIEKAVAVFAGGVHLIEITFTVPDADTVIKALSVLKEKGAIIGAGTVTSVEQARKAVESGAEFIV
SPHLDEEISQFAKEKGVFYMPGVMTPTELVKAMKLGHTILKLFPGEVVGPQFVKAMKGPFPNVKFVPTGG
VNLDNVAEWFKAGVLAVGVGSALVKGTPDEVREKAKAFVEKIRGATEGGSHHHHHHHH

>Rpk11-I53-50A trimer
MGILPSPGMPALLSLVSLLSVLLMGCVAETGTRFPNITNLCPLGEVFNATRFASVYAWNRKRISNCVLDM
SVLYNSASFSTFKCYGVSPTKLNDLCFTNVYADSFVIRGDEVRQIAPGQTGKIADYNYKLPDDFTGCVIA
WNSNNLDSKVGGNYNYLYRLFRKSNLKPFERDISTEIYQAGSTPCNGVEGFNCYFPLQSYGFQPTNGVGY
QPYRVVVLSFELLHAPATVCGPKKSTGGSGGSGSGGSGGSGSEKAAKAEEAARKMEELFKKHKIVAVLRA
NSVEEAIEKAVAVFAGGVHLIEITFTVPDADTVIKALSVLKEKGAIIGAGTVTSVEQARKAVESGAEFIV
SPHLDEEISQFAKEKGVFYMPGVMTPTELVKAMKLGHTILKLFPGEVVGPQFVKAMKGPFPNVKFVPTGG
VNLDNVAEWFKAGVLAVGVGSALVKGTPDEVREKAKAFVEKIRGATEGGSHHHHHHHH

>Rpk12-I53-50A trimer
MGILPSPGMPALLSLVSLLSVLLMGCVAETGTRFPNITNLCPFGEVFNATRFASVYAWNRKRFSNCVADW
SVLYNSASFSTFKCYGVSPTKLNDLCFTNVYADSFVIRGDEVRQIAPGQTGKIADYNYKLPDDFTGCVIA
WNSNNLDSKVGGNYNYLYRLFRKSNLKPFERDISTEIYQAGSTPCNGVEGFNCYFPLQSYGFQPTNGVGY
QPYRVVVLSFELLHAPATVCGPKKSTGGSGGSGSGGSGGSGSEKAAKAEEAARKMEELFKKHKIVAVLRA

NSVEEAIEKAVAVFAGGVHLIEITFTVPDADTVIKALSVLKEKGAIIGAGTVTSVEQARKAVESGAEFIV
SPHLDEEISQFAKEKGVFYMPGVMTPTELVKAMKLGHTILKLFPGEVVGPQFVKAMKGPFPNVKFVPTGG
VNLDNVAEWFKAGVLAVGVGSALVKGTPDEVREKAKAFVEKIRGATEGGSHHHHHHHH

>Rpk13-I53-50A trimer
MGILPSPGMPALLSLVSLLSVLLMGCVAETGTRFPNITNLCPLGEVFNATRFASVYAWNRKRFSNCVADW
SVLYNSASFSTFKCYGVSPTKLNDLCFTNVYADSFVIRGDEVRQIAPGQTGKIADYNYKLPDDFTGCVIA
WNSNNLDSKVGGNYNYLYRLFRKSNLKPFERDISTEIYQAGSTPCNGVEGFNCYFPLQSYGFQPTNGVGY
QPYRVVVLSFELLHAPATVCGPKKSTGGSGGSGSGGSGGSGSEKAAKAEEAARKMEELFKKHKIVAVLRA
NSVEEAIEKAVAVFAGGVHLIEITFTVPDADTVIKALSVLKEKGAIIGAGTVTSVEQARKAVESGAEFIV
SPHLDEEISQFAKEKGVFYMPGVMTPTELVKAMKLGHTILKLFPGEVVGPQFVKAMKGPFPNVKFVPTGG
VNLDNVAEWFKAGVLAVGVGSALVKGTPDEVREKAKAFVEKIRGATEGGSHHHHHHHH

>Rpk14-I53-50A trimer
MGILPSPGMPALLSLVSLLSVLLMGCVAETGTRFPNITNLCPFGEVFNATRFASVYAWNRKRFSNCVADW
SVLYNSASFSTFKCYGVSPTKLNDLCFTNVYADSFVIRGDEVRQIAPGQTGKIADYNYKLPDDFTGCVIA
WNSNNLDSKVGGNYNYLYRLFRKSNLKPFERDISTEIYQAGSTPCNGVEGFNCYFPLQSYGFQPTNGVGY
QPYRVVVMSFELLHAPATVCGPKKSTGGSGGSGSGGSGGSGSEKAAKAEEAARKMEELFKKHKIVAVLRA
NSVEEAIEKAVAVFAGGVHLIEITFTVPDADTVIKALSVLKEKGAIIGAGTVTSVEQARKAVESGAEFIV
SPHLDEEISQFAKEKGVFYMPGVMTPTELVKAMKLGHTILKLFPGEVVGPQFVKAMKGPFPNVKFVPTGG
VNLDNVAEWFKAGVLAVGVGSALVKGTPDEVREKAKAFVEKIRGATEGGSHHHHHHHH

>Rpk15-I53-50A trimer
MGILPSPGMPALLSLVSLLSVLLMGCVAETGTRFPNITNLCPFGEVFNATRFASVYAWNRKRFSNCVADF
SVLYNSASFSTFKCYGVSPTKLNDLCFTNIYADSFVIRGDEVRQIAPGQTGKIADYNYKLPDDFTGCVIA
WNSNNLDSKVGGNYNYLYRLFRKSNLKPFERDISTEIYQAGSTPCNGVEGFNCYFPLQSYGFQPTNGVGY
QPYRVVVLSFELLHAPATVCGPKKSTGGSGGSGSGGSGGSGSEKAAKAEEAARKMEELFKKHKIVAVLRA
NSVEEAIEKAVAVFAGGVHLIEITFTVPDADTVIKALSVLKEKGAIIGAGTVTSVEQARKAVESGAEFIV
SPHLDEEISQFAKEKGVFYMPGVMTPTELVKAMKLGHTILKLFPGEVVGPQFVKAMKGPFPNVKFVPTGG
VNLDNVAEWFKAGVLAVGVGSALVKGTPDEVREKAKAFVEKIRGATEGGSHHHHHHHH

>Rpk16-I53-50A trimer
MGILPSPGMPALLSLVSLLSVLLMGCVAETGTRFPNITNLCPFGEVFNATRFASVYAWNRKRFSNCVADW
SVLYNSASFSTFKCYGVSPTKLNDLCWTNVYADSFVIRGDEVRQIAPGQTGKIADYNYKLPDDFTGCVIA
WNSNNLDSKVGGNYNYLYRLFRKSNLKPFERDISTEIYQAGSTPCNGVEGFNCYFPLQSYGFQPTNGVGY
QPYRVVVLSFELLHAPATVCGPKKSTGGSGGSGSGGSGGSGSEKAAKAEEAARKMEELFKKHKIVAVLRA
NSVEEAIEKAVAVFAGGVHLIEITFTVPDADTVIKALSVLKEKGAIIGAGTVTSVEQARKAVESGAEFIV

SPHLDEEISQFAKEKGVFYMPGVMTPTELVKAMKLGHTILKLFPGEVVGPQFVKAMKGPFPNVKFVPTGG
VNLDNVAEWFKAGVLAVGVGSALVKGTPDEVREKAKAFVEKIRGATEGGSHHHHHHHH

>Rpk17-I53-50A trimer
MGILPSPGMPALLSLVSLLSVLLMGCVAETGTRFPNITNLCPFGEVFNATRFASVYAWNRKRFSNCVADF
SVLYNSASFSTFKCYGVSPTKLNDLCWTNIYADSFVIRGDEVRQIAPGQTGKIADYNYKLPDDFTGCVIA
WNSNNLDSKVGGNYNYLYRLFRKSNLKPFERDISTEIYQAGSTPCNGVEGFNCYFPLQSYGFQPTNGVGY
QPYRVVVLSFELLHAPATVCGPKKSTGGSGGSGSGGSGGSGSEKAAKAEEAARKMEELFKKHKIVAVLRA
NSVEEAIEKAVAVFAGGVHLIEITFTVPDADTVIKALSVLKEKGAIIGAGTVTSVEQARKAVESGAEFIV
SPHLDEEISQFAKEKGVFYMPGVMTPTELVKAMKLGHTILKLFPGEVVGPQFVKAMKGPFPNVKFVPTGG
VNLDNVAEWFKAGVLAVGVGSALVKGTPDEVREKAKAFVEKIRGATEGGSHHHHHHHH

>RBD monomer (with Avi and hexa-histidine tags)
MGILPSPGMPALLSLVSLLSVLLMGCVAETGTRFPNITNLCPFGEVFNATRFASVYAWNRKRISNCVADY
SVLYNSASFSTFKCYGVSPTKLNDLCFTNVYADSFVIRGDEVRQIAPGQTGKIADYNYKLPDDFTGCVIA
WNSNNLDSKVGGNYNYLYRLFRKSNLKPFERDISTEIYQAGSTPCNGVEGFNCYFPLQSYGFQPTNGVGY
QPYRVVVLSFELLHAPATVCGPKKSTGLNDIFEAQKIEWHEHHHHHHHH

>Rpk4 monomer (with Avi and hexa-histidine tags)
MGILPSPGMPALLSLVSLLSVLLMGCVAETGTRFPNITNLCPFGEVFNATRFASVYAWNRKRISNCVADY
SVLYNSASFSTFKCYGVSPTKLNDLCWTNVYADSFVIRGDEVRQIAPGQTGKIADYNYKLPDDFTGCVIA
WNSNNLDSKVGGNYNYLYRLFRKSNLKPFERDISTEIYQAGSTPCNGVEGFNCYFPLQSYGFQPTNGVGY
QPYRVVVLSFELLHAPATVCGPKKSTGLNDIFEAQKIEWHEHHHHHHHH

>Rpk9 monomer (with Avi and hexa-histidine tags)
MGILPSPGMPALLSLVSLLSVLLMGCVAETGTRFPNITNLCPFGEVFNATRFASVYAWNRKRISNCVADF
SVLYNSASFSTFKCYGVSPTKLNDLCWTNIYADSFVIRGDEVRQIAPGQTGKIADYNYKLPDDFTGCVIA
WNSNNLDSKVGGNYNYLYRLFRKSNLKPFERDISTEIYQAGSTPCNGVEGFNCYFPLQSYGFQPTNGVGY
QPYRVVVLSFELLHAPATVCGPKKSTGLNDIFEAQKIEWHEHHHHHHHH

>I53-50B.4PT1 pentamer
MNQHSHKDHETVRIAVVRARWHAEIVDACVSAFEAAMRDIGGDRFAVDVFDVPGAYEIPLHARTLAETGR
YGAVLGTAFVVNGGIYRHEFVASAVINGMMNVQLNTGVPVLSAVLTPHNYDKSKAHTLLFLALFAVKGME
AARACVEILAAREKIAAGSLEHHHHHH

>2OBX pentamer

MNQHSHKDYETVRIAVVRARWHADIVDQCVSAFEAEMADIGGDRFAVDVFDVPGAYEIPLHARTLAETGR
YGAVLGTAFVVNGGIYRHEFVASAVIDGMMNVQLSTGVPVLSAVLTPHNYHDSAEHHRFFFEHFTVKGKE
AARACVEILAAREKIAAGSLEHHHHHH

>Hexapro-foldon, used for immunizations (Wuhan-Hu-1)
MFVFLVLLPLVSSQCVNLTTRTQLPPAYTNSFTRGVYYPDKVFRSSVLHSTQDLFLPFFSNVTWFHAIHV
SGTNGTKRFDNPVLPFNDGVYFASTEKSNIIRGWIFGTTLDSKTQSLLIVNNATNVVIKVCEFQFCNDPF
LGVYYHKNNKSWMESEFRVYSSANNCTFEYVSQPFLMDLEGKQGNFKNLREFVFKNIDGYFKIYSKHTPI
NLVRDLPQGFSALEPLVDLPIGINITRFQTLLALHRSYLTPGDSSSGWTAGAAAYYVGYLQPRTFLLKYN
ENGTITDAVDCALDPLSETKCTLKSFTVEKGIYQTSNFRVQPTESIVRFPNITNLCPFGEVFNATRFASV
YAWNRKRISNCVADYSVLYNSASFSTFKCYGVSPTKLNDLCFTNVYADSFVIRGDEVRQIAPGQTGKIAD
YNYKLPDDFTGCVIAWNSNNLDSKVGGNYNYLYRLFRKSNLKPFERDISTEIYQAGSTPCNGVEGFNCYF
PLQSYGFQPTNGVGYQPYRVVVLSFELLHAPATVCGPKKSTNLVKNKCVNFNFNGLTGTGVLTESNKKFL
PFQQFGRDIADTTDAVRDPQTLEILDITPCSFGGVSVITPGTNTSNQVAVLYQDVNCTEVPVAIHADQLT
PTWRVYSTGSNVFQTRAGCLIGAEHVNNSYECDIPIGAGICASYQTQTNSPGSASSVASQSIIAYTMSLG
AENSVAYSNNSIAIPTNFTISVTTEILPVSMTKTSVDCTMYICGDSTECSNLLLQYGSFCTQLNRALTGI
AVEQDKNTQEVFAQVKQIYKTPPIKDFGGFNFSQILPDPSKPSKRSPIEDLLFNKVTLADAGFIKQYGDC
LGDIAARDLICAQKFNGLTVLPPLLTDEMIAQYTSALLAGTITSGWTFGAGPALQIPFPMQMAYRFNGIG
VTQNVLYENQKLIANQFNSAIGKIQDSLSSTPSALGKLQDVVNQNAQALNTLVKQLSSNFGAISSVLNDI
LSRLDPPEAEVQIDRLITGRLQSLQTYVTQQLIRAAEIRASANLAATKMSECVLGQSKRVDFCGKGYHLM
SFPQSAPHGVVFLHVTYVPAQEKNFTTAPAICHDGKAHFPREGVFVSNGTHWFVTQRNFYEPQIITTDNT
FVSGNCDVVIGIVNNTVYDPLQPELDSFKEELDKYFKNHTSPDVDLGDISGINASVVNIQKEIDRLNEVA
KNLNESLIDLQELGKYEQGSGYIPEAPRDGQAYVRKDGEWVLLSTFLGRSLEVLFQGPGHHHHHHHHSAW
SHPQFEKGGGSGGGGSGGSAWSHPQFEK

>Hexapro-foldon, used for expression and stability comparisons with
Rpk9-Hexapro-foldon (Wuhan-Hu-1)
MARAWIFFLLCLAGRALAQCVNLTTRTQLPPAYTNSFTRGVYYPDKVFRSSVLHSTQDLFLPFFSNVTWF
HAIHVSGTNGTKRFDNPVLPFNDGVYFASTEKSNIIRGWIFGTTLDSKTQSLLIVNNATNVVIKVCEFQF
CNDPFLGVYYHKNNKSWMESEFRVYSSANNCTFEYVSQPFLMDLEGKQGNFKNLREFVFKNIDGYFKIYS
KHTPINLVRDLPQGFSALEPLVDLPIGINITRFQTLLALHRSYLTPGDSSSGWTAGAAAYYVGYLQPRTF
LLKYNENGTITDAVDCALDPLSETKCTLKSFTVEKGIYQTSNFRVQPTESIVRFPNITNLCPFGEVFNAT
RFASVYAWNRKRISNCVADYSVLYNSASFSTFKCYGVSPTKLNDLCFTNVYADSFVIRGDEVRQIAPGQT
GKIADYNYKLPDDFTGCVIAWNSNNLDSKVGGNYNYLYRLFRKSNLKPFERDISTEIYQAGSTPCNGVEG
FNCYFPLQSYGFQPTNGVGYQPYRVVVLSFELLHAPATVCGPKKSTNLVKNKCVNFNFNGLTGTGVLTES
NKKFLPFQQFGRDIADTTDAVRDPQTLEILDITPCSFGGVSVITPGTNTSNQVAVLYQDVNCTEVPVAIH
ADQLTPTWRVYSTGSNVFQTRAGCLIGAEHVNNSYECDIPIGAGICASYQTQTNSPGSASSVASQSIIAY
TMSLGAENSVAYSNNSIAIPTNFTISVTTEILPVSMTKTSVDCTMYICGDSTECSNLLLQYGSFCTQLNR

ALTGIAVEQDKNTQEVFAQVKQIYKTPPIKDFGGFNFSQILPDPSKPSKRSPIEDLLFNKVTLADAGFIK
QYGDCLGDIAARDLICAQKFNGLTVLPPLLTDEMIAQYTSALLAGTITSGWTFGAGPALQIPFPMQMAYR
FNGIGVTQNVLYENQKLIANQFNSAIGKIQDSLSSTPSALGKLQDVVNQNAQALNTLVKQLSSNFGAISS
VLNDILSRLDPPEAEVQIDRLITGRLQSLQTYVTQQLIRAAEIRASANLAATKMSECVLGQSKRVDFCGK
GYHLMSFPQSAPHGVVFLHVTYVPAQEKNFTTAPAICHDGKAHFPREGVFVSNGTHWFVTQRNFYEPQII
TTDNTFVSGNCDVVIGIVNNTVYDPLQPELDSFKEELDKYFKNHTSPDVDLGDISGINASVVNIQKEIDR
LNEVAKNLNESLIDLQELGKYEQGSGYIPEAPRDGQAYVRKDGEWVLLSTFLGRSLEVLFQGPGHHHHHH
HH

>Rpk9-Hexapro-foldon (Wuhan-Hu-1)
MARAWIFFLLCLAGRALAQCVNLTTRTQLPPAYTNSFTRGVYYPDKVFRSSVLHSTQDLFLPFFSNVTWF
HAIHVSGTNGTKRFDNPVLPFNDGVYFASTEKSNIIRGWIFGTTLDSKTQSLLIVNNATNVVIKVCEFQF
CNDPFLGVYYHKNNKSWMESEFRVYSSANNCTFEYVSQPFLMDLEGKQGNFKNLREFVFKNIDGYFKIYS
KHTPINLVRDLPQGFSALEPLVDLPIGINITRFQTLLALHRSYLTPGDSSSGWTAGAAAYYVGYLQPRTF
LLKYNENGTITDAVDCALDPLSETKCTLKSFTVEKGIYQTSNFRVQPTESIVRFPNITNLCPFGEVFNAT
RFASVYAWNRKRISNCVADFSVLYNSASFSTFKCYGVSPTKLNDLCWTNIYADSFVIRGDEVRQIAPGQT
GKIADYNYKLPDDFTGCVIAWNSNNLDSKVGGNYNYLYRLFRKSNLKPFERDISTEIYQAGSTPCNGVEG
FNCYFPLQSYGFQPTNGVGYQPYRVVVLSFELLHAPATVCGPKKSTNLVKNKCVNFNFNGLTGTGVLTES
NKKFLPFQQFGRDIADTTDAVRDPQTLEILDITPCSFGGVSVITPGTNTSNQVAVLYQDVNCTEVPVAIH
ADQLTPTWRVYSTGSNVFQTRAGCLIGAEHVNNSYECDIPIGAGICASYQTQTNSPGSASSVASQSIIAY
TMSLGAENSVAYSNNSIAIPTNFTISVTTEILPVSMTKTSVDCTMYICGDSTECSNLLLQYGSFCTQLNR
ALTGIAVEQDKNTQEVFAQVKQIYKTPPIKDFGGFNFSQILPDPSKPSKRSPIEDLLFNKVTLADAGFIK
QYGDCLGDIAARDLICAQKFNGLTVLPPLLTDEMIAQYTSALLAGTITSGWTFGAGPALQIPFPMQMAYR
FNGIGVTQNVLYENQKLIANQFNSAIGKIQDSLSSTPSALGKLQDVVNQNAQALNTLVKQLSSNFGAISS
VLNDILSRLDPPEAEVQIDRLITGRLQSLQTYVTQQLIRAAEIRASANLAATKMSECVLGQSKRVDFCGK
GYHLMSFPQSAPHGVVFLHVTYVPAQEKNFTTAPAICHDGKAHFPREGVFVSNGTHWFVTQRNFYEPQII
TTDNTFVSGNCDVVIGIVNNTVYDPLQPELDSFKEELDKYFKNHTSPDVDLGDISGINASVVNIQKEIDR
LNEVAKNLNESLIDLQELGKYEQGSGYIPEAPRDGQAYVRKDGEWVLLSTFLGRSLEVLFQGPGHHHHHH
HH

>hACE2-FC
MARAWIFFLLCLAGRALASTIEEQAKTFLDKFNHEAEDLFYQSSLASWNYNTNITEENVQNMNNAGDKWS
AFLKEQSTLAQMYPLQEIQNLTVKLQLQALQQNGSSVLSEDKSKRLNTILNTMSTIYSTGKVCNPDNPQE
CLLLEPGLNEIMANSLDYNERLWAWESWRSEVGKQLRPLYEEYVVLKNEMARANHYEDYGDYWRGDYEVN
GVDGYDYSRGQLIEDVEHTFEEIKPLYEHLHAYVRAKLMNAYPSYISPIGCLPAHLLGDMWGRFWTNLYS
LTVPFGQKPNIDVTDAMVDQAWDAQRIFKEAEKFFVSVGLPNMTQGFWENSMLTDPGNVQKAVCHPTAWD
LGKGDFRILMCTKVTMDDFLTAHHEMGHIQYDMAYAAQPFLLRNGANEGFHEAVGEIMSLSAATPKHLKS
IGLLSPDFQEDNETEINFLLKQALTIVGTLPFTYMLEKWRWMVFKGEIPKDQWMKKWWEMKREIVGVVEP

VPHDETYCDPASLFHVSNDYSFIRYYTRTLYQFQFQEALCQAAKHEGPLHKCDISNSTEAGQKLFNMLRL
GKSEPWTLALENVVGAKNMNVRPLLNYFEPLFTWLKDQNKNSFVGWSTDWSPYADPLVPRGSGGGGDPEP
KSCDKTHTCPPCPAPELLGGPSVFLFPPKPKDTLMISRTPEVTCVVVDVSHEDPEVKFNWYVDGVEVHNA
KTKPREEQYNSTYRVVSVLTVLHQDWLNGKEYKCKVSNKALPAPIEKTISKAKGQPREPQVYTLPPSRDE
LTKNQVSLTCLVKGFYPSDIAVEWESNGQPENNYKTTPPVLDSDGSFFLYSKLTVDKSRWQQGNVFSCSV
MHEALHNHYTQKSLSLSPGK
