## Supplementary Information 3 for "Stabilization of the SARS-CoV-2 Spike receptor-binding domain using deep mutational scanning and structure-based design"

### Supplementary Solution Stability Data

### SDS-PAGE for nanoparticles in TBS, 5% glycerol, 0.75% CHAPS, 100 mM L-arginine

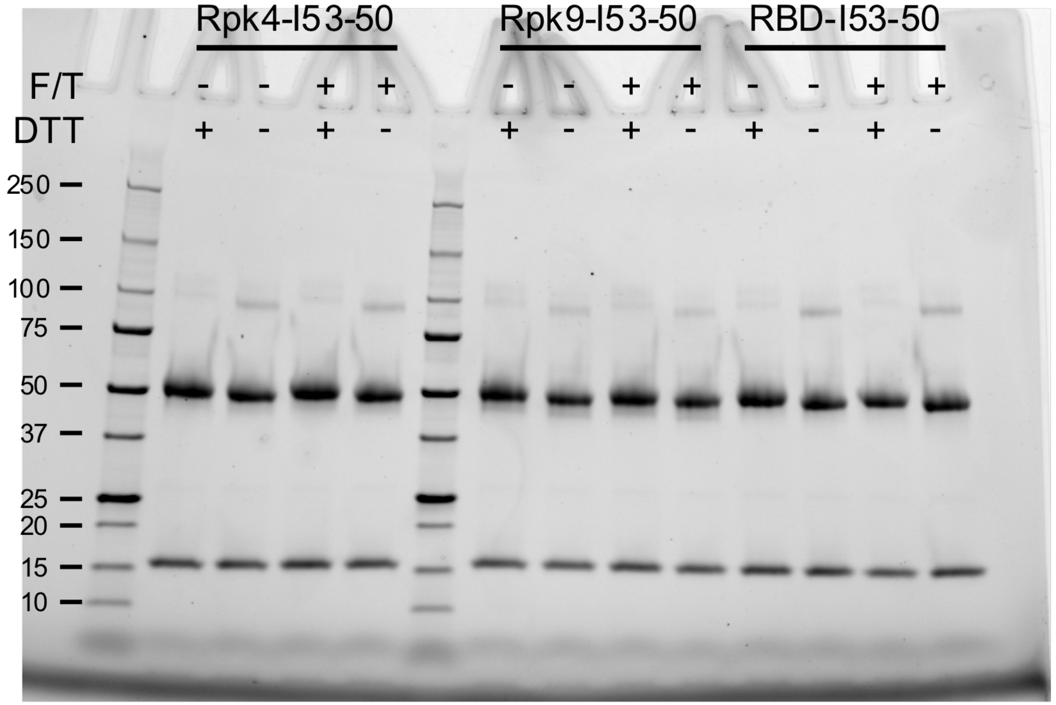


The integrity of samples in 50 mM Tris pH 7.4, 185 mM NaCl, 4.5% glycerol, 0.75% CHAPS, 100 mM L-arginine was analyzed by SDS-PAGE. Molecular weights of the standard are noted in kDa. Each sample was analyzed +/- reducing agent (DTT), pre- and post-freeze thaw (F/T).

### hACE2-Fc binding for nanoparticles in TBS, 5% glycerol, 0.75% CHAPS, 100 mM L-arginine

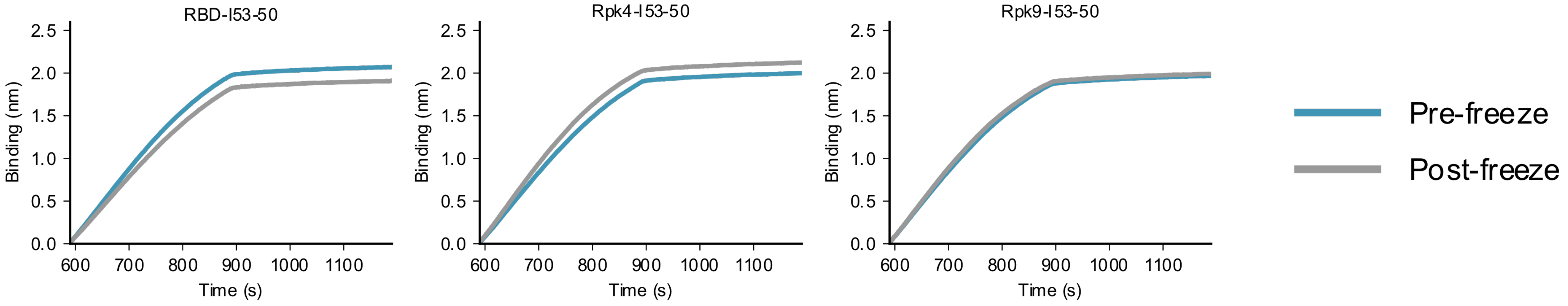


hACE2-Fc binding of antigen in 50 mM Tris pH 7.4, 185 mM NaCl, 4.5% glycerol, 0.75% CHAPS, 100 mM L-arginine was analyzed by bio-layer interferometry (BLI). Protein A biosensors loaded with hACE2-Fc were incubated with immunogen (association, x = 590–889 s) and then buffer (dissociation, x = 890–1190 s).

### CR3022 binding for nanoparticles in TBS, 5% glycerol, 0.75% CHAPS, 100 mM L-arginine

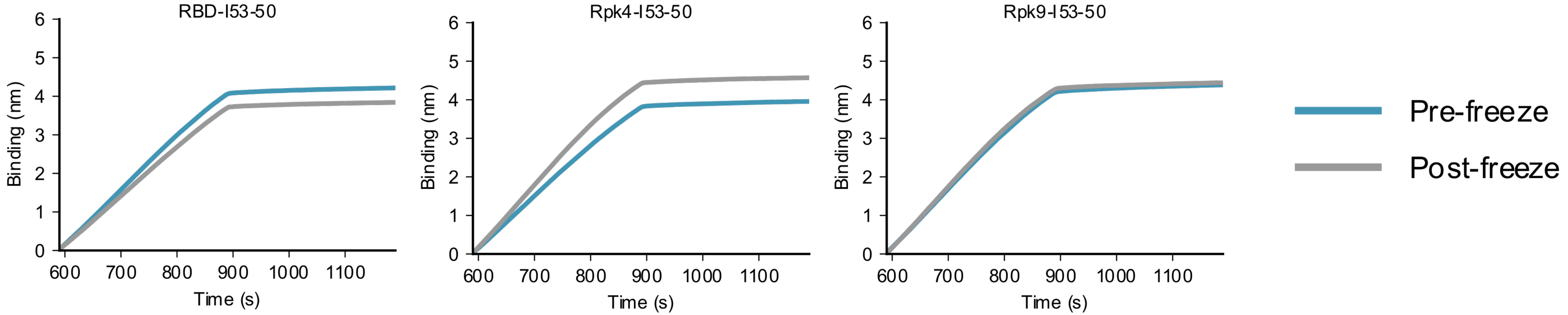


CR3022 IgG binding of antigen in 50 mM Tris pH 7.4, 185 mM NaCl, 4.5% glycerol, 0.75% CHAPS, 100 mM L-arginine was analyzed by bio-layer interferometry (BLI). Protein A biosensors loaded with CR3022 IgG were incubated with immunogen (association, x = 590–889 s) and then buffer (dissociation, x = 890–1190 s).

### Dynamic light scattering for nanoparticles in TBS, 5% glycerol, 0.75% CHAPS, 100 mM L-arginine

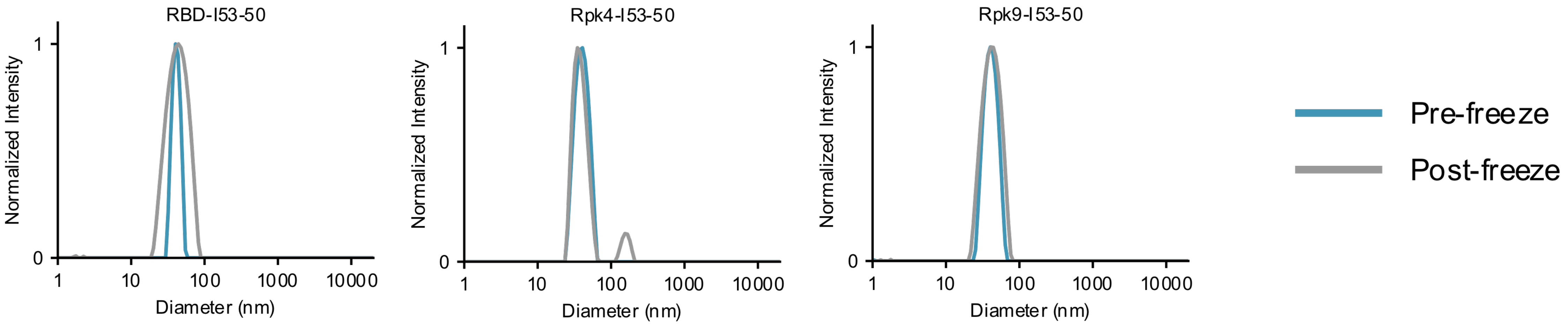


Hydrodynamic diameter (nm) for each sample in 50 mM Tris pH 7.4, 185 mM NaCl, 4.5% glycerol, 0.75% CHAPS, 100 mM L-arginine, plotted as normalized intensity.

### UV-Vis for nanoparticles in TBS, 5% glycerol, 0.75% CHAPS, 100 mM L-arginine

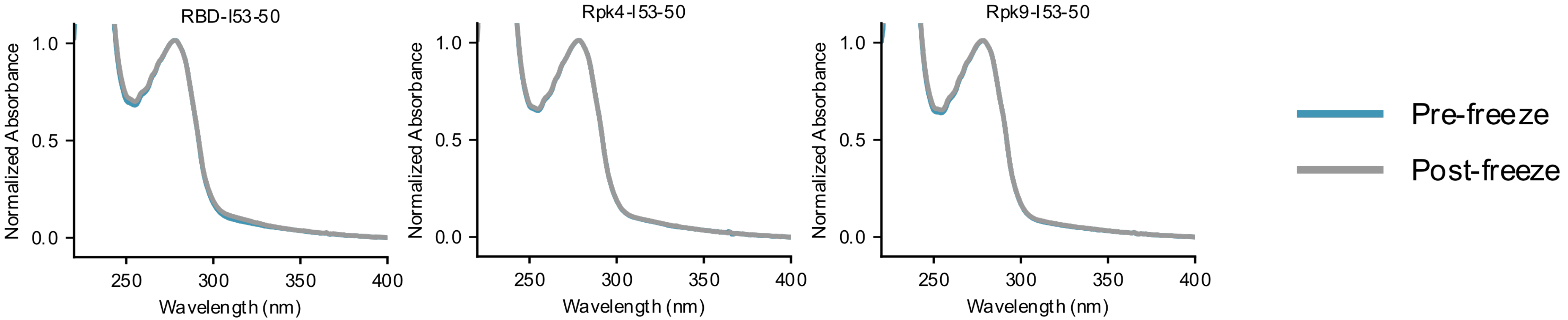


UV-Vis spectra (nm) for each sample in 50 mM Tris pH 7.4, 185 mM NaCl, 4.5% glycerol, 0.75% CHAPS, 100 mM L-arginine, plotted as normalized absorbance.

### SDS-PAGE for nanoparticles in TBS, 5% glycerol, 100 mM L-arginine

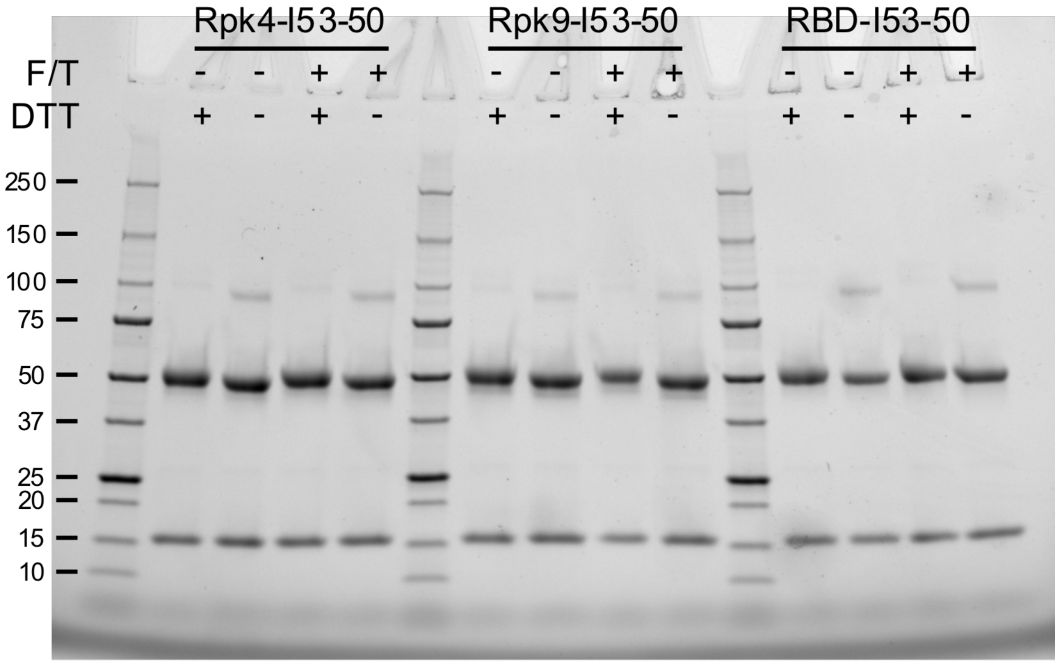


The integrity of samples in 50 mM Tris pH 8, 150 mM NaCl, 5% glycerol, 100 mM L-arginine was analyzed by SDS-PAGE. Molecular weights of the standard are noted in kDa. Each sample was analyzed +/- reducing agent (DTT), pre- and post-freeze thaw (F/T).

### hACE2-Fc binding for nanoparticles in TBS, 5% glycerol, 100 mM L-arginine

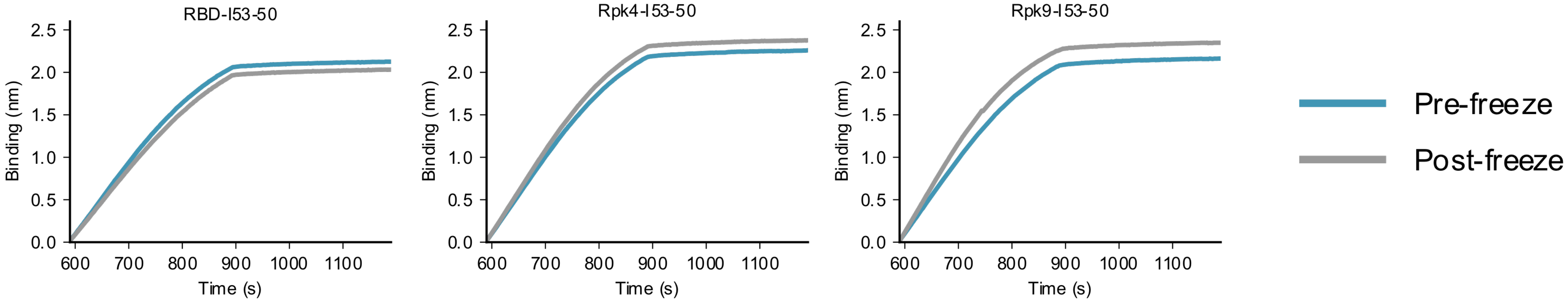


hACE2-Fc binding of antigen in 50 mM Tris pH 8, 150 mM NaCl, 5% glycerol, 100 mM L-arginine was analyzed by bio-layer interferometry (BLI). Protein A biosensors loaded with hACE2-Fc were incubated with immunogen (association, x = 590–889 s) and then buffer (dissociation, x = 890–1190 s).

### CR3022 binding for nanoparticles in TBS, 5% glycerol, 100 mM L-arginine

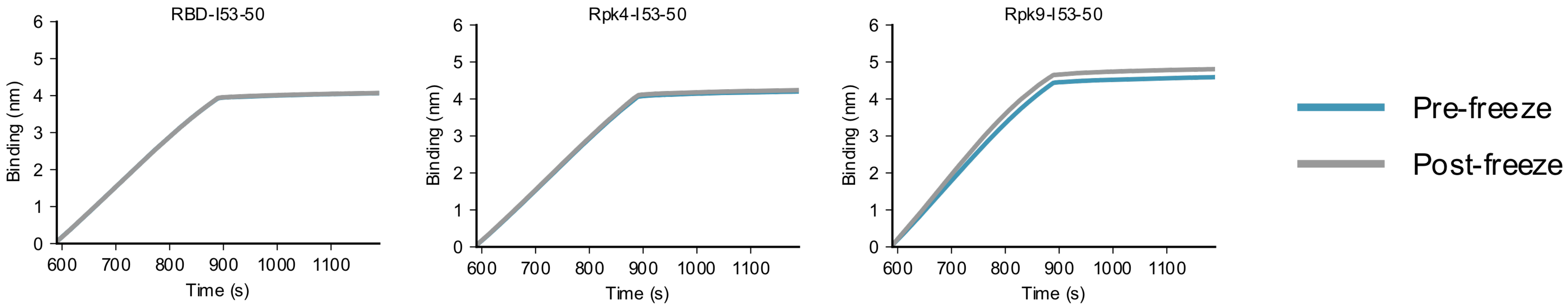


CR3022 IgG binding of antigen in 50 mM Tris pH 8, 150 mM NaCl, 5% glycerol, 100 mM L-arginine was analyzed by bio-layer interferometry (BLI). Protein A biosensors loaded with CR3022 IgG were incubated with immunogen (association, x = 590–889 s) and then buffer (dissociation, x = 890–1190 s).

### Dynamic light scattering for nanoparticles in TBS, 5% glycerol, 100 mM L-arginine

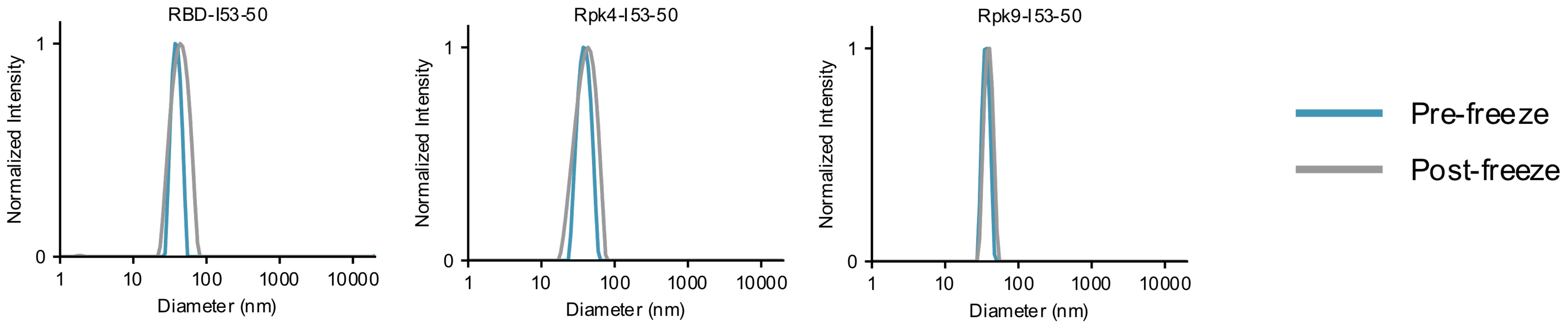


Hydrodynamic diameter (nm) for each sample in 50 mM Tris pH 8, 150 mM NaCl, 5% glycerol, 100 mM L-arginine, plotted as normalized intensity.

### UV-Vis for nanoparticles in TBS, 5% glycerol, 100 mM L-arginine

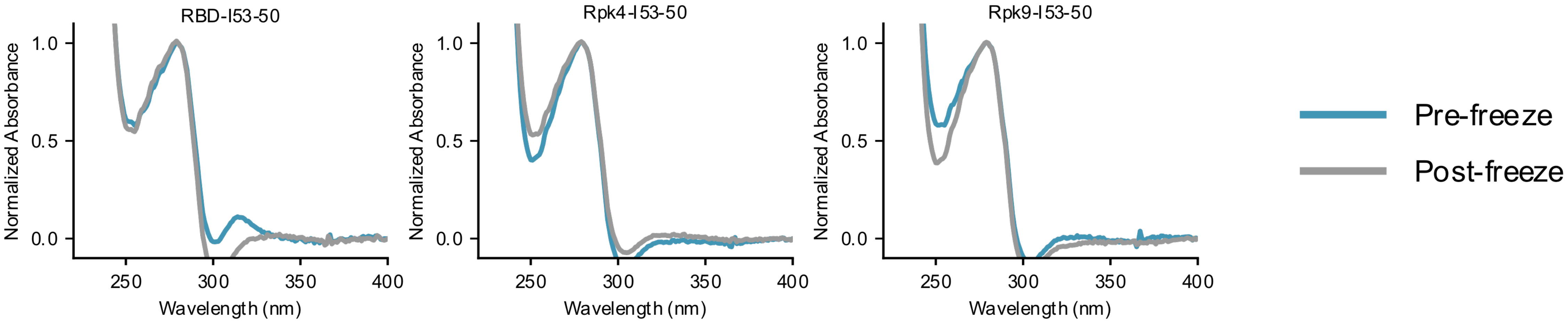


UV-Vis spectra (nm) for each sample in 50 mM Tris pH 8, 150 mM NaCl, 5% glycerol, 100 mM L-arginine, plotted as normalized absorbance.

### SDS-PAGE for nanoparticles in TBS, 5% glycerol

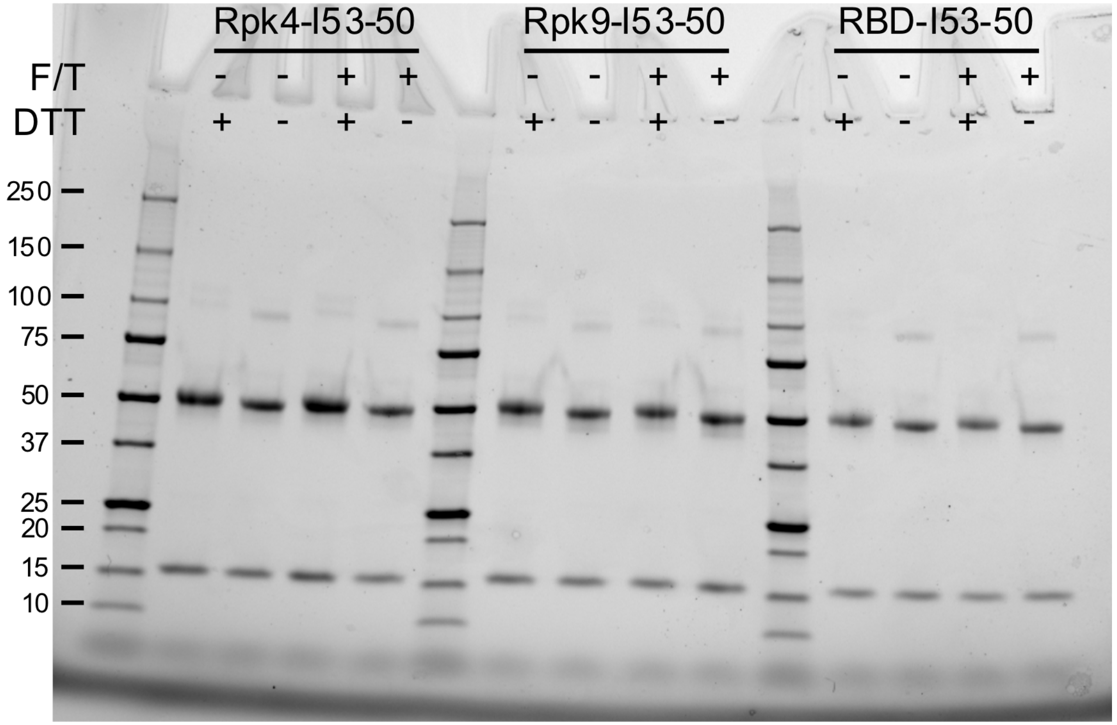


The integrity of samples in 50 mM Tris pH 8, 150 mM NaCl, 5% glycerol was analyzed by SDS-PAGE. Molecular weights of the standard are noted in kDa. Each sample was analyzed +/- reducing agent (DTT), pre- and post-freeze thaw (F/T).

### hACE2-Fc binding for nanoparticles in TBS, 5% glycerol

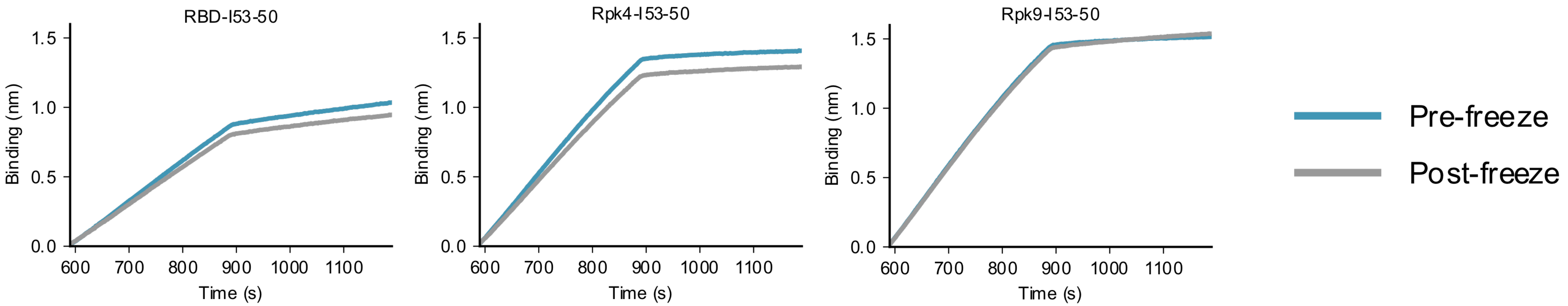


hACE2-Fc binding of antigen in 50 mM Tris pH 8, 150 mM NaCl, 5% glycerol was analyzed by bio-layer interferometry (BLI). Protein A biosensors loaded with hACE2-Fc were incubated with immunogen (association, x = 590–889 s) and then buffer (dissociation, x = 890–1190 s).

### CR3022 binding for nanoparticles in TBS, 5% glycerol

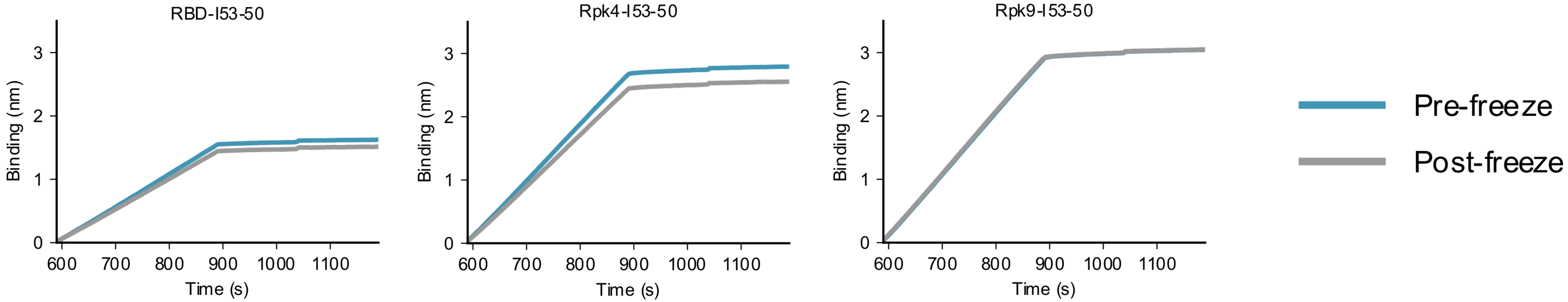


CR3022 IgG binding of antigen in 50 mM Tris pH 8, 150 mM NaCl, 5% glycerol was analyzed by bio-layer interferometry (BLI). Protein A biosensors loaded with CR3022 IgG were incubated with immunogen (association, x = 590–889 s) and then buffer (dissociation, x = 890–1190 s).

### Dynamic light scattering for nanoparticles in TBS, 5% glycerol

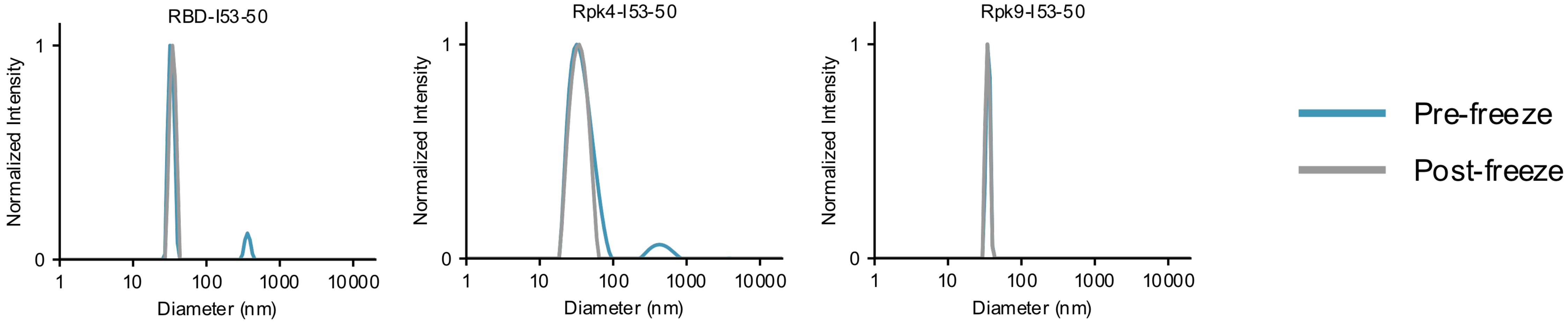


Hydrodynamic diameter (nm) for each sample in 50 mM Tris pH 8, 150 mM NaCl, 5% glycerol, plotted as normalized intensity.

### UV-Vis for nanoparticles in TBS, 5% glycerol

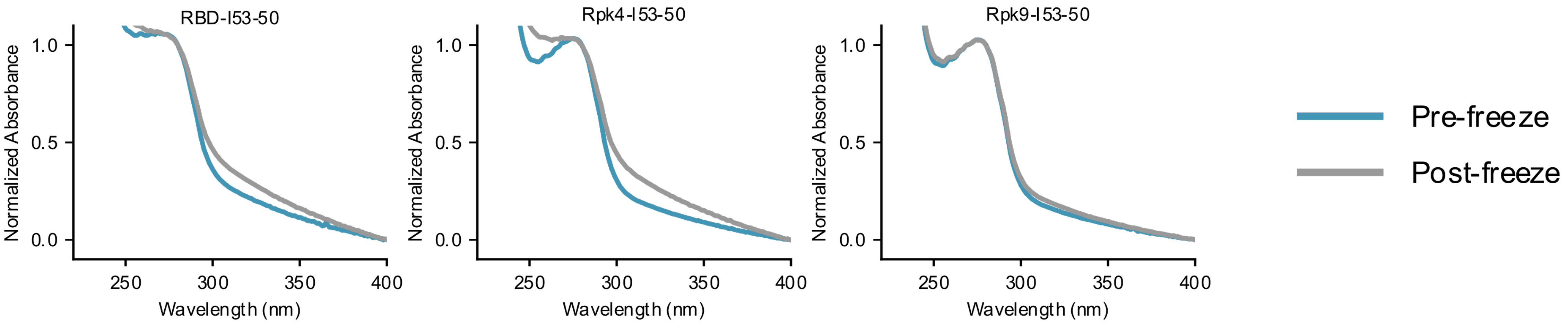


UV-Vis spectra (nm) for each sample in 50 mM Tris pH 8, 150 mM NaCl, 5% glycerol, plotted as normalized absorbance.

### SDS-PAGE for nanoparticles in TBS

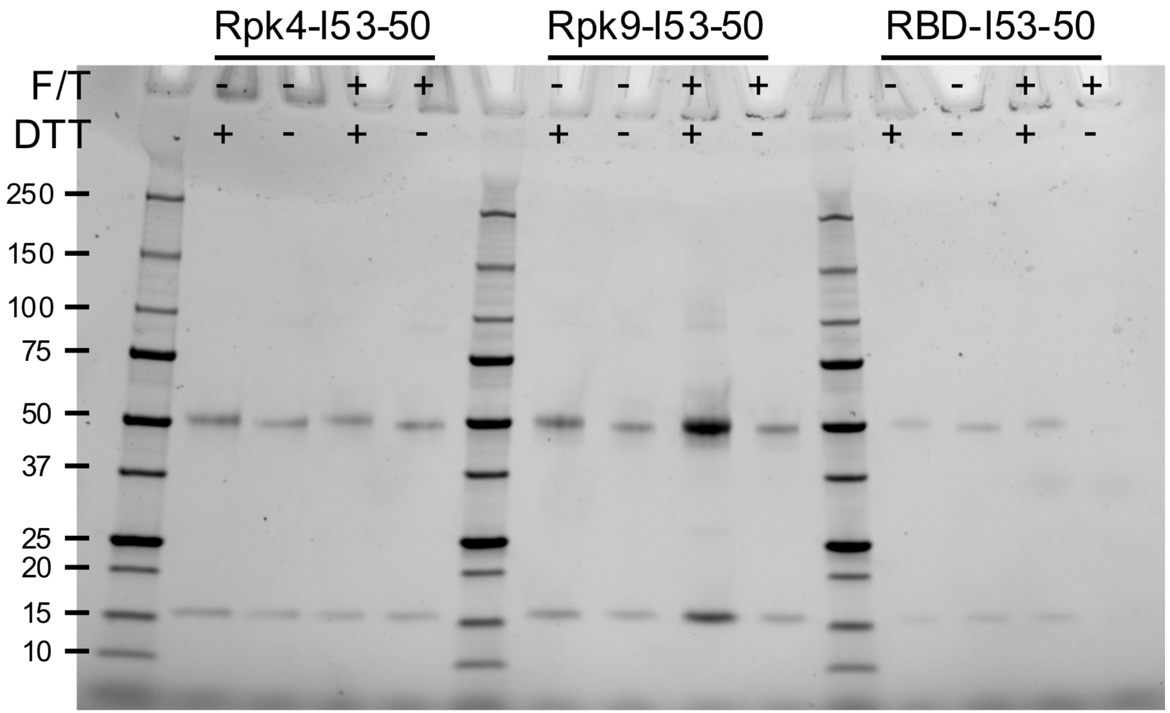


The integrity of samples in 50 mM Tris pH 8, 150 mM NaCl was analyzed by SDS-PAGE. Molecular weights of the standard are noted in kDa. Each sample was analyzed +/- reducing agent (DTT), pre- and post-freeze thaw (F/T).

### hACE2-Fc binding for nanoparticles in TBS

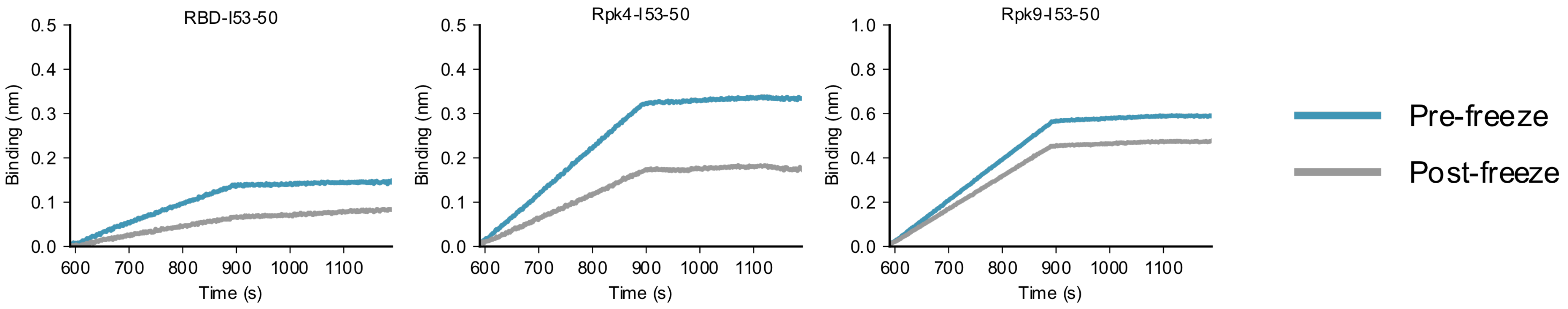


hACE2-Fc binding of antigen in 50 mM Tris pH 8, 150 mM NaCl was analyzed by bio-layer interferometry (BLI). Protein A biosensors loaded with hACE2-Fc were incubated with immunogen (association, x = 590–889 s) and then buffer (dissociation, x = 890–1190 s).

### CR3022 binding for nanoparticles in TBS

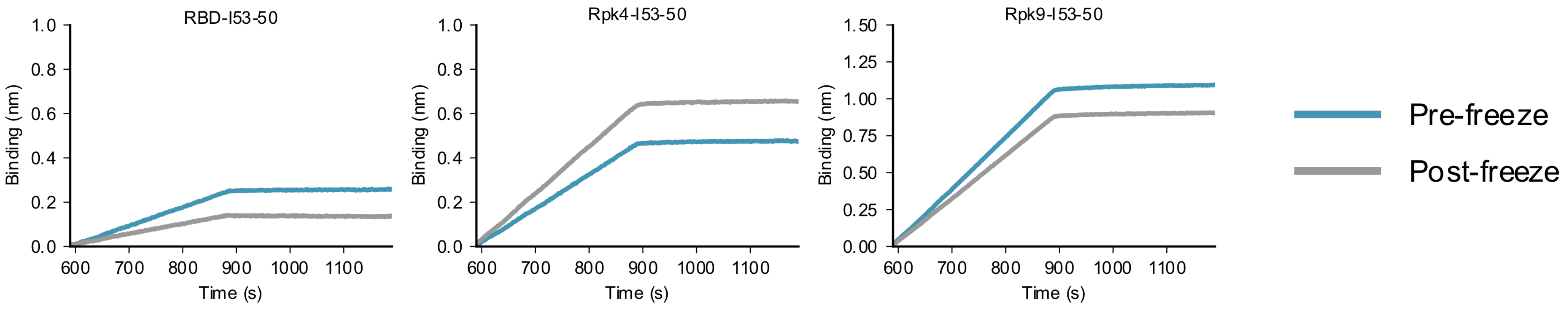


CR3022 IgG binding of antigen in 50 mM Tris pH 8, 150 mM NaCl was analyzed by bio-layer interferometry (BLI). Protein A biosensors loaded with CR3022 IgG were incubated with immunogen (association, x = 590–889 s) and then buffer (dissociation, x = 890–1190 s).

### Dynamic light scattering for nanoparticles in TBS

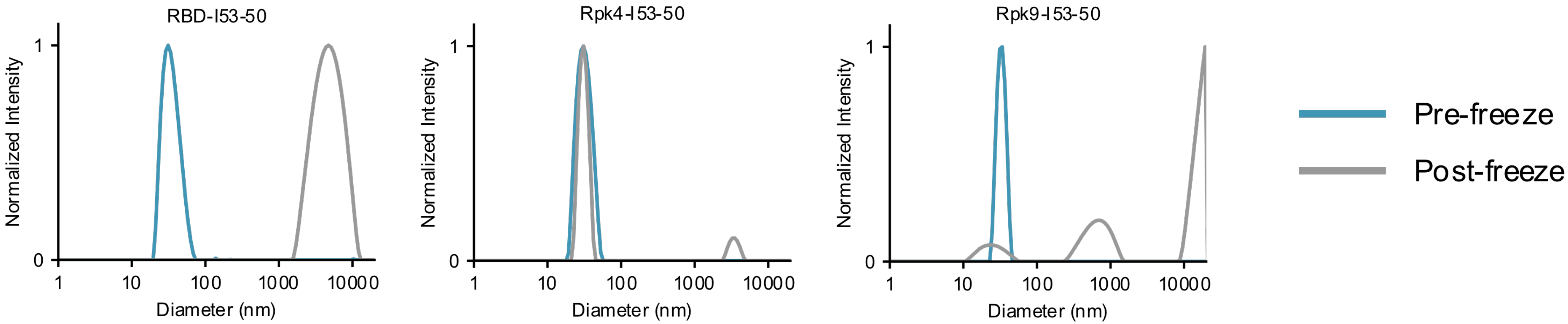


Hydrodynamic diameter (nm) for each sample in 50 mM Tris pH 8, 150 mM NaCl, plotted as normalized intensity.

### UV-Vis for nanoparticles in TBS

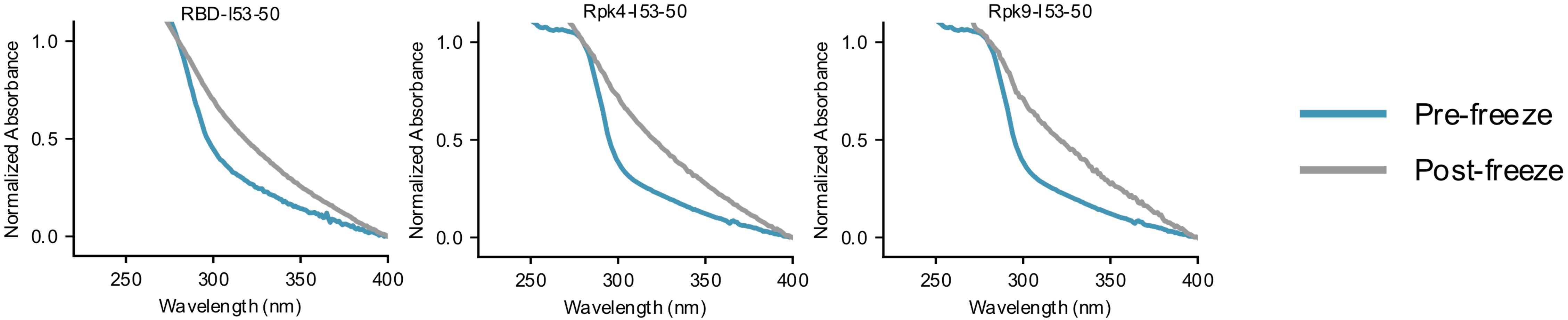


UV-vis spectra (nm) for each sample in 50 mM Tris pH 8, 150 mM NaCl, plotted as normalized absorbance.
