## Supplementary Information 4 for "Stabilization of the SARS-CoV-2 Spike receptor-binding domain using deep mutational scanning and structure-based design"

#### Supplementary Shelf-life Stability Data

### SDS-PAGE for RBD-I53-50 nanoparticle

D1

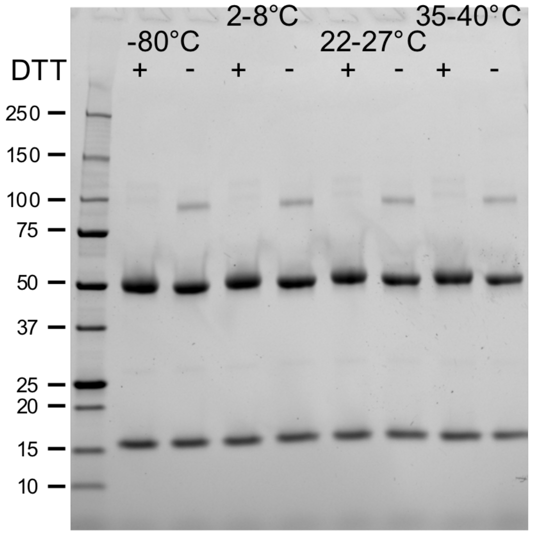


D3

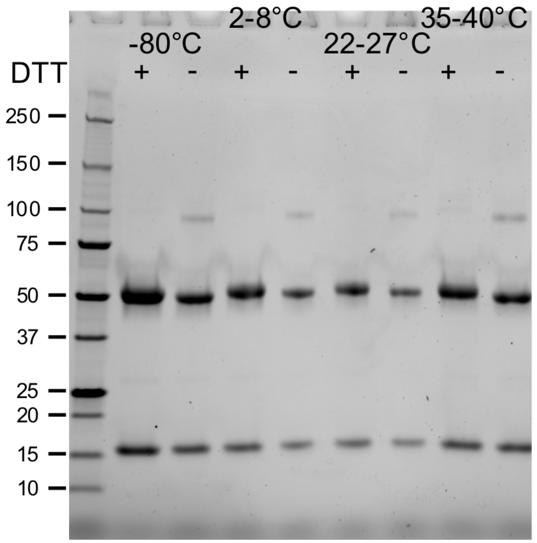


D7

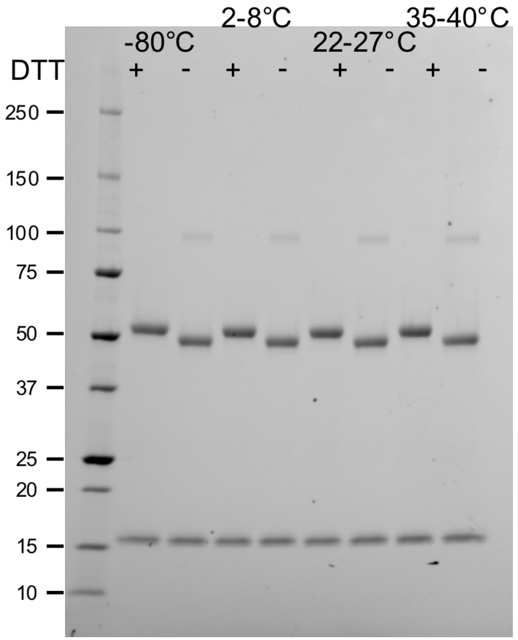


D14

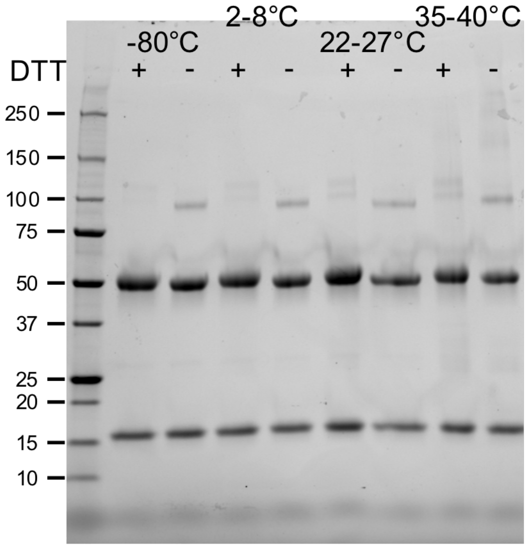


D28

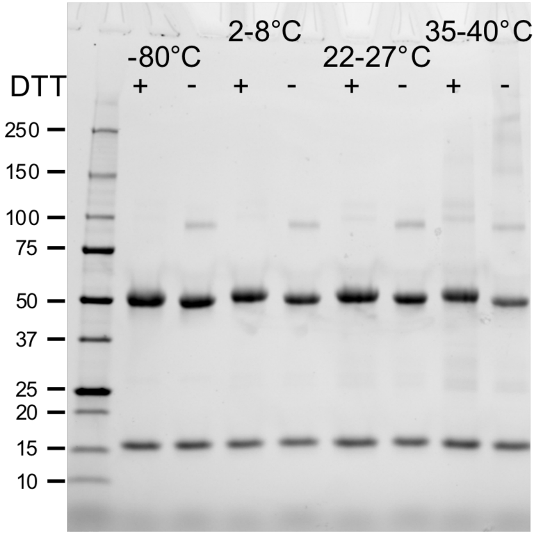


The integrity of samples after incubation at four temperatures over a 28 day (D) study was analyzed by SDS-PAGE. Molecular weights of the standard are noted in kDa. Each sample was analyzed +/- reducing agent (DTT).

### hACE2-Fc binding for RBD-I53-50 nanoparticle

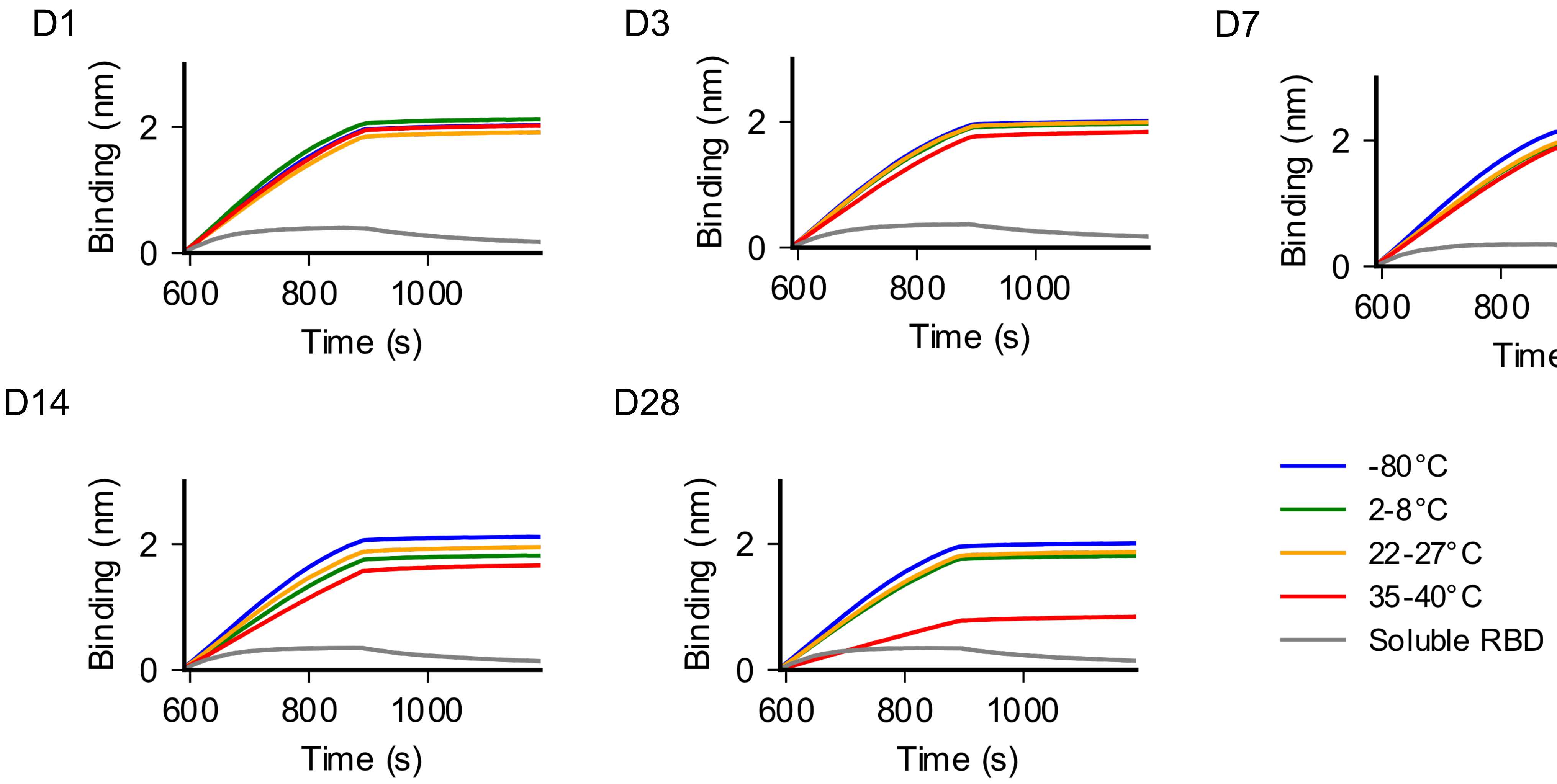


hACE2-Fc binding of antigen incubated at four different temperatures for 28 days (D) was analyzed by bio-layer Interferometry (BLI). Protein A biosensors loaded with hACE2-Fc were incubated with immunogen (association, x = 590–889 s) and then buffer (dissociation, x = 890–1190 s).

### CR3022 binding for RBD-I53-50 nanoparticle

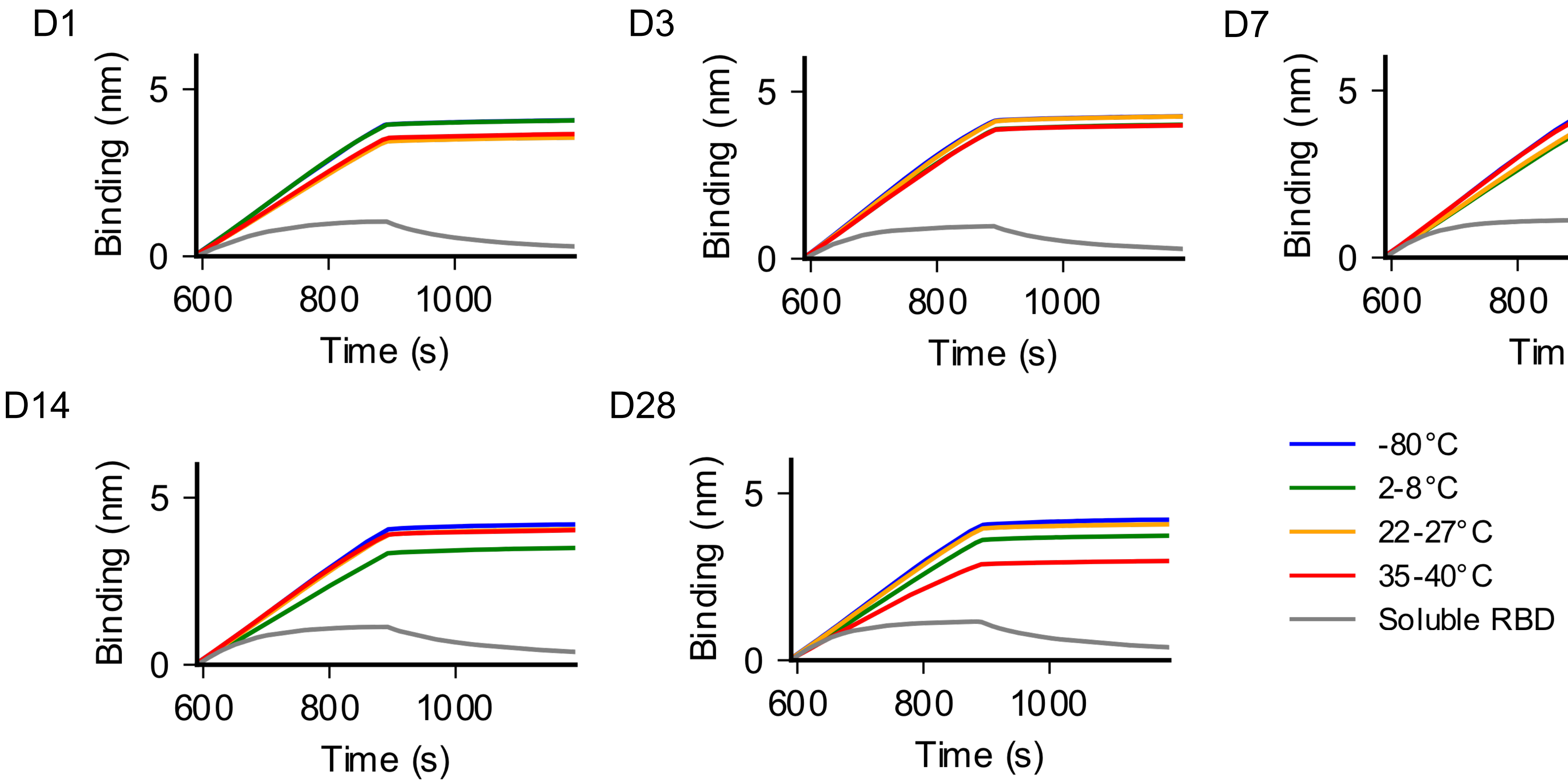


CR3022 IgG binding of antigen incubated at four different temperatures for 28 days (D) was analyzed by bio-layer Interferometry (BLI). Protein A biosensors loaded with CR3022 were incubated with immunogen (association, x = 590–889 s) and then buffer (dissociation, x = 890–1190 s).

### nsEM for RBD-I53-50 nanoparticle

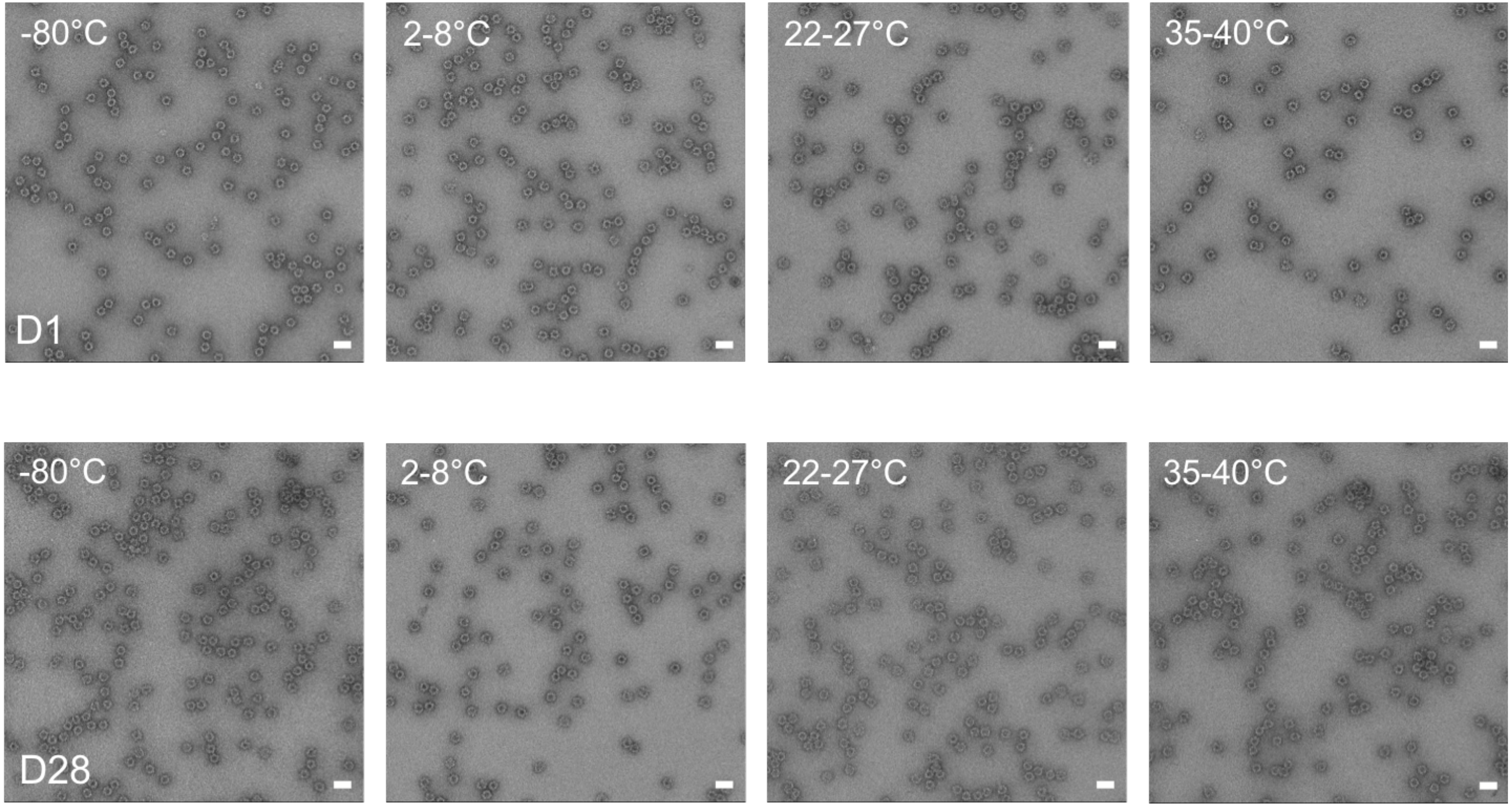


Representative negative stain electron micrographs for each sample at days (D) 1 and 28 following incubation at four temperatures. Scale bar, 50 nm.

### Dynamic light scattering for RBD-I53-50 nanoparticle

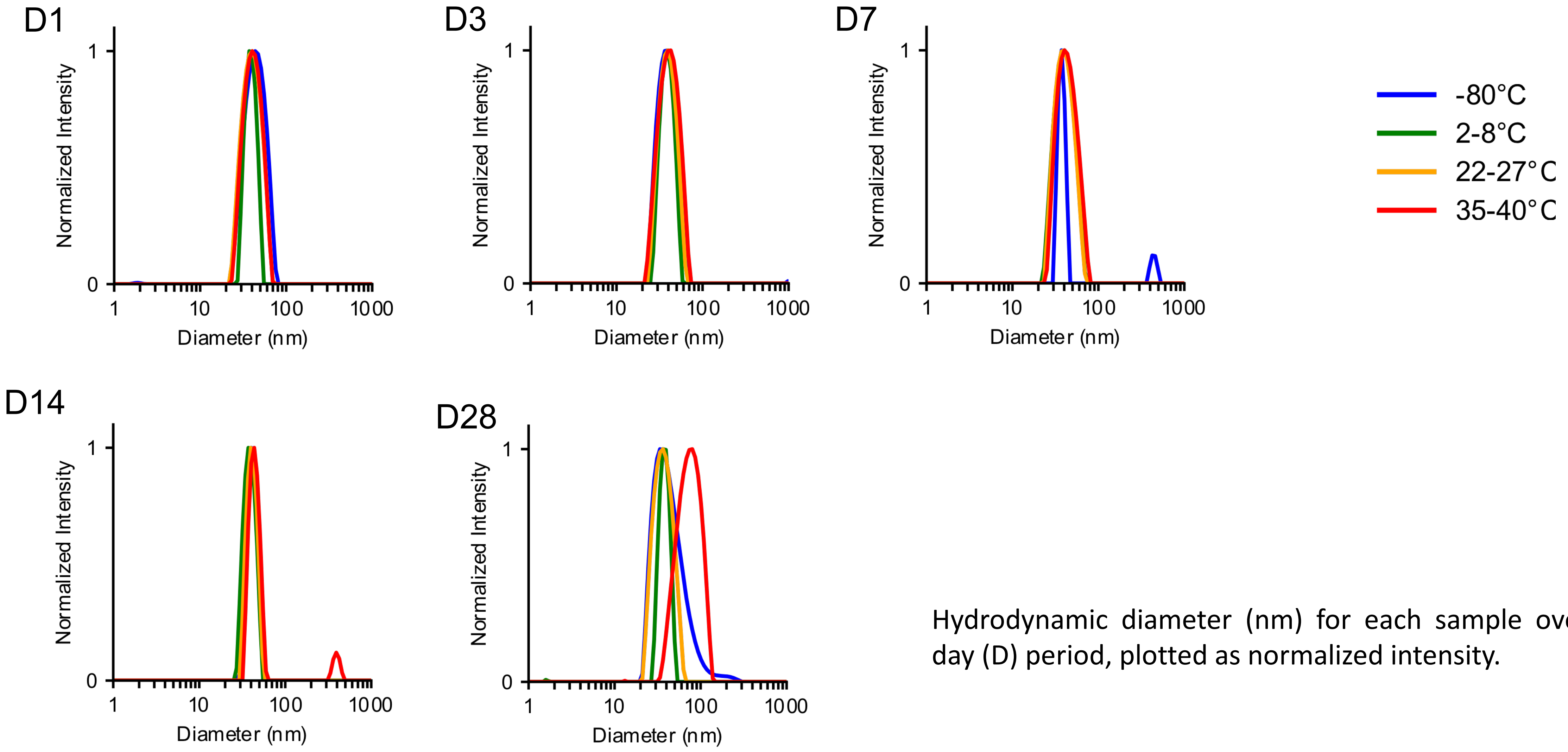


Hydrodynamic diameter (nm) for each sample over a 28 day (D) period, plotted as normalized intensity.

### SDS-PAGE for Rpk4-I53-50 nanoparticle

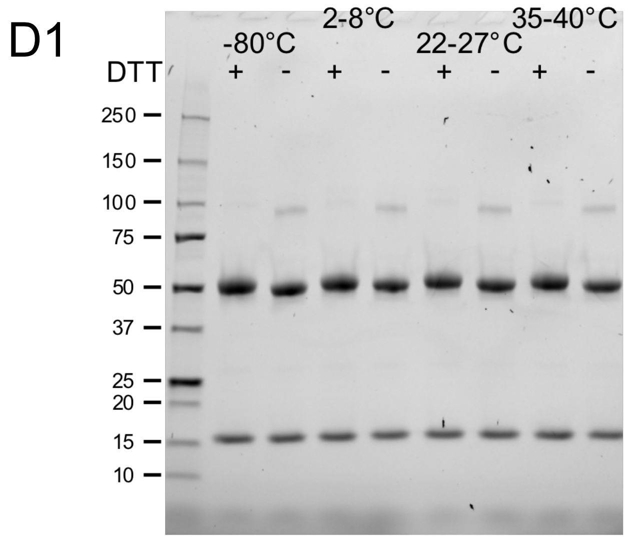


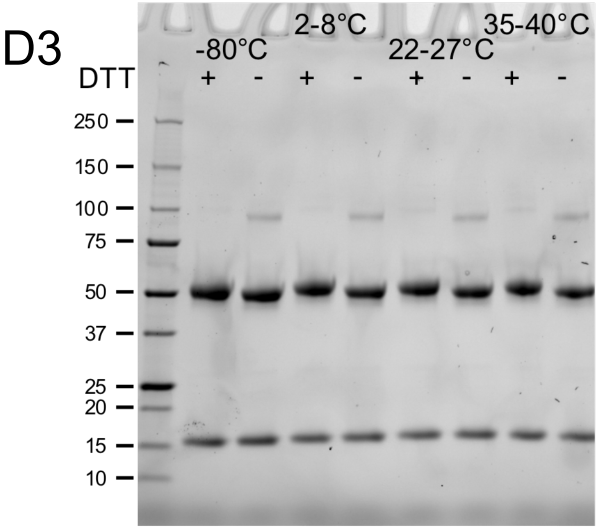


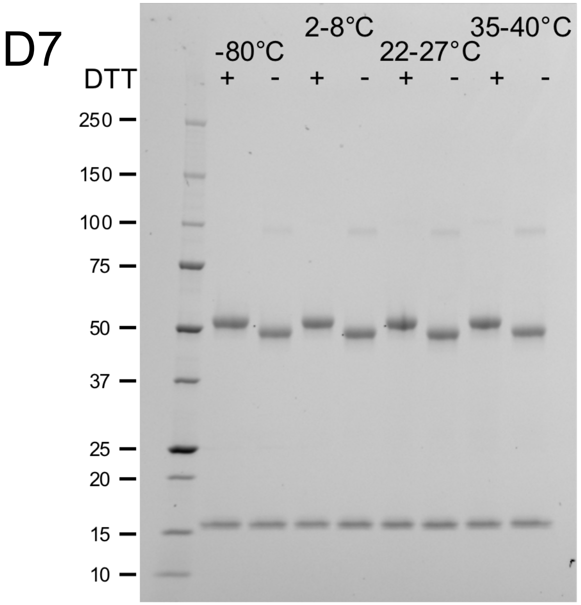


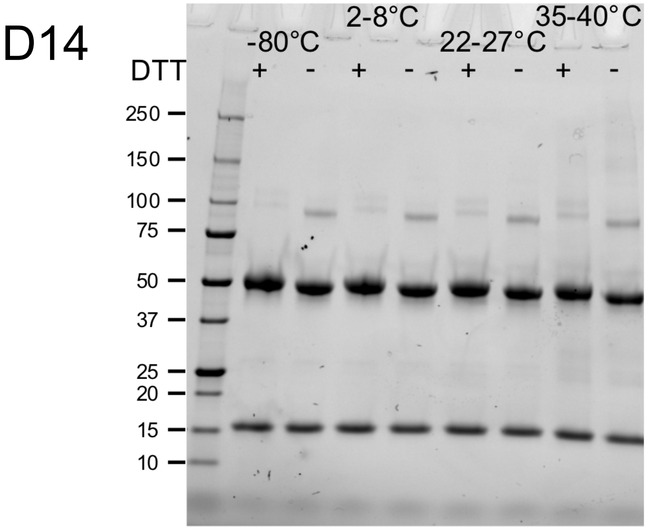


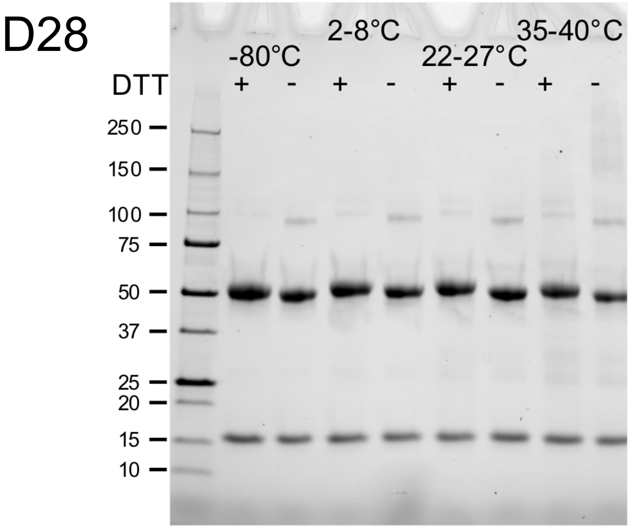


The integrity of samples after incubation at four temperatures over a 28 day (D) study was analyzed by SDS-PAGE. Molecular weights of the standard are noted in kDa. Each sample was analyzed +/- reducing agent (DTT).

### hACE2-Fc binding for Rpk4-I53-50 nanoparticle

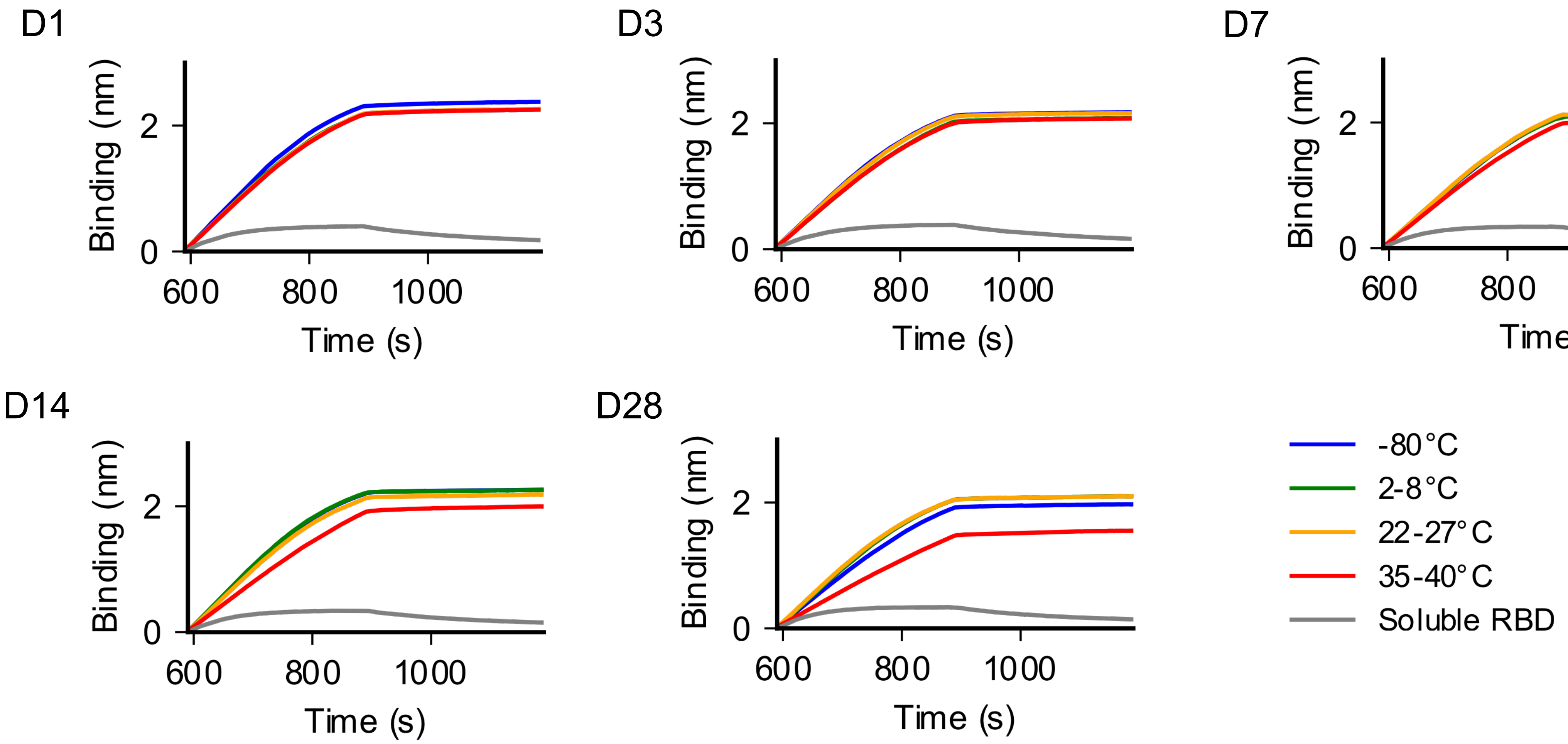


hACE2-Fc binding of antigen incubated at four different temperatures for 28 days (D) was analyzed by bio-layer Interferometry (BLI). Protein A biosensors loaded with hACE2-Fc were incubated with immunogen (association, x = 590–889 s) and then buffer (dissociation, x = 890–1190 s).

### CR3022 binding for Rpk4-I53-50 nanoparticle

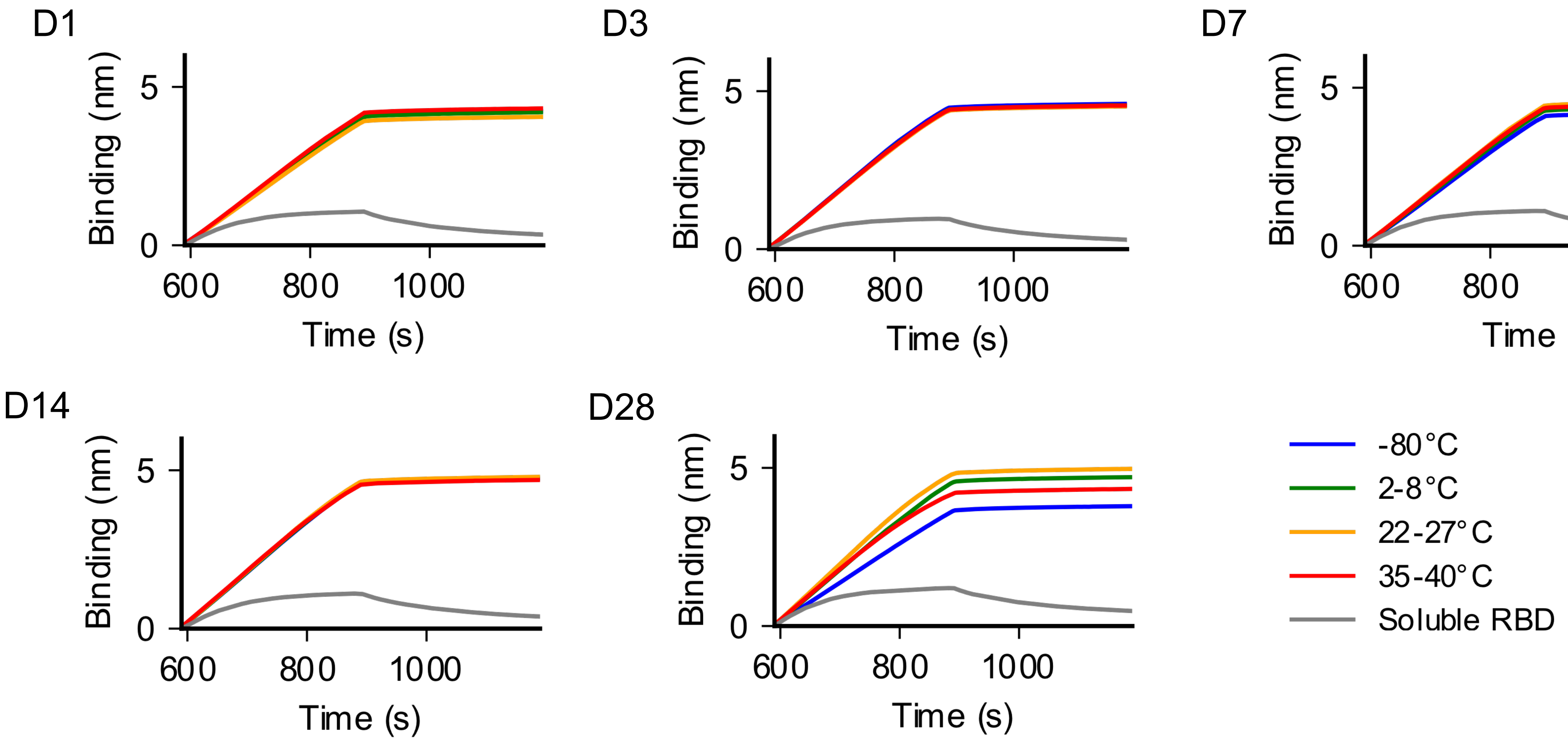


CR3022 IgG binding of antigen incubated at four different temperatures for 28 days (D) was analyzed by bio-layer Interferometry (BLI). Protein A biosensors loaded with CR3022 were incubated with immunogen (association, x = 590–889 s) and then buffer (dissociation, x = 890–1190 s).

### nsEM for Rpk4-I53-50 nanoparticle

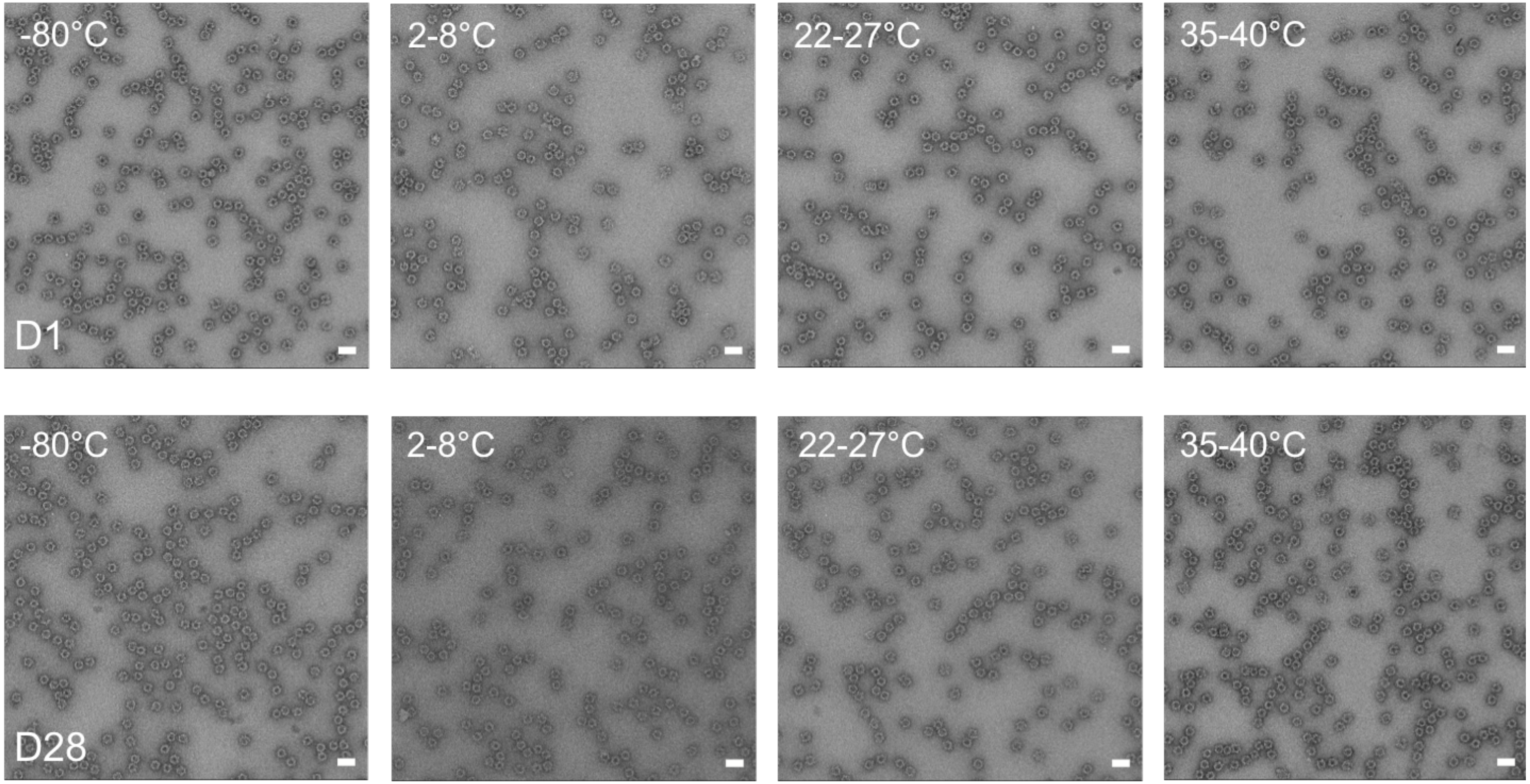


Representative negative stain electron micrographs for each sample at days (D) 1 and 28 following incubation at four temperatures. Scale bar, 50 nm.

### Dynamic light scattering for Rpk4-I53-50 nanoparticle

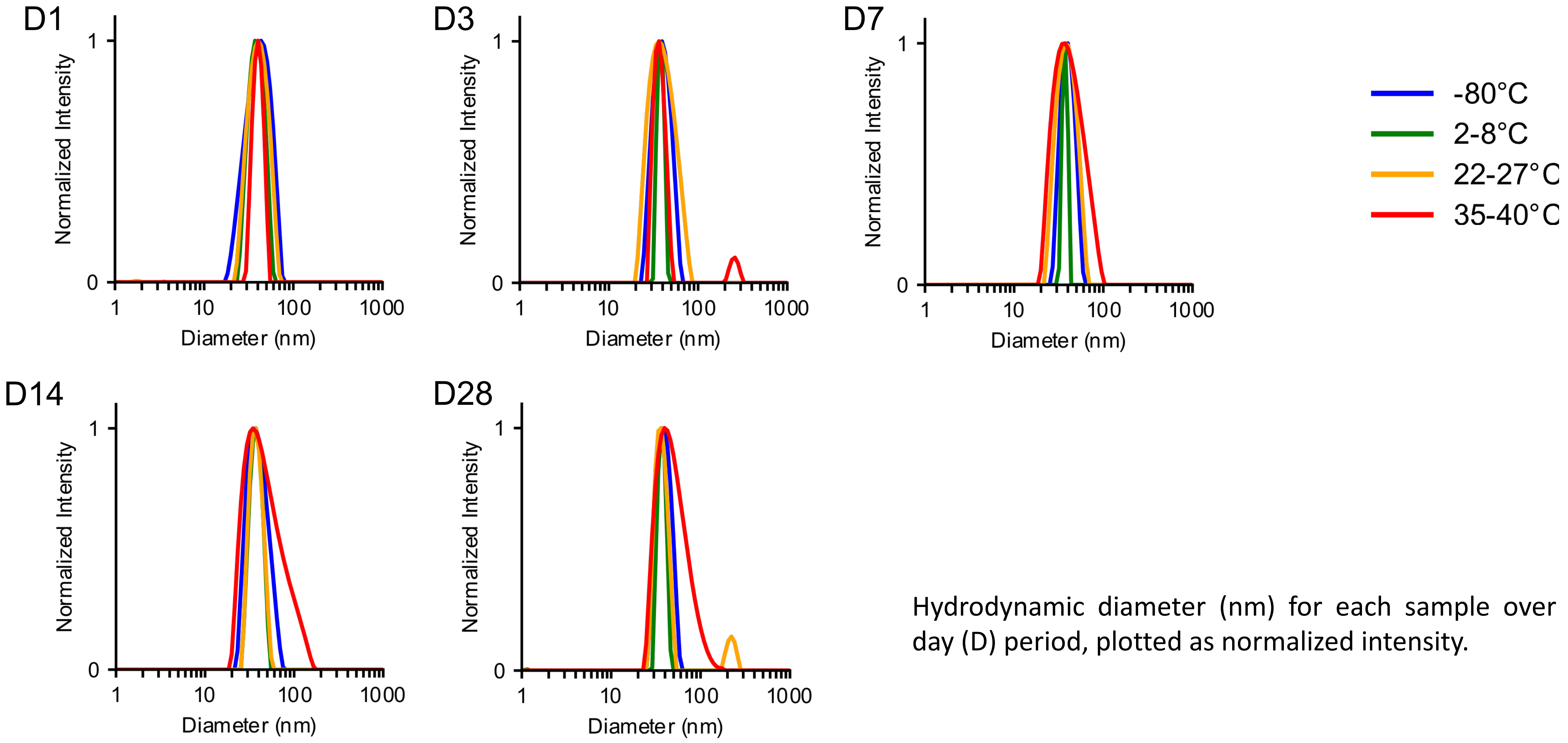


Hydrodynamic diameter (nm) for each sample over a 28 day (D) period, plotted as normalized intensity.

### SDS-PAGE for Rpk9-I53-50 nanoparticle

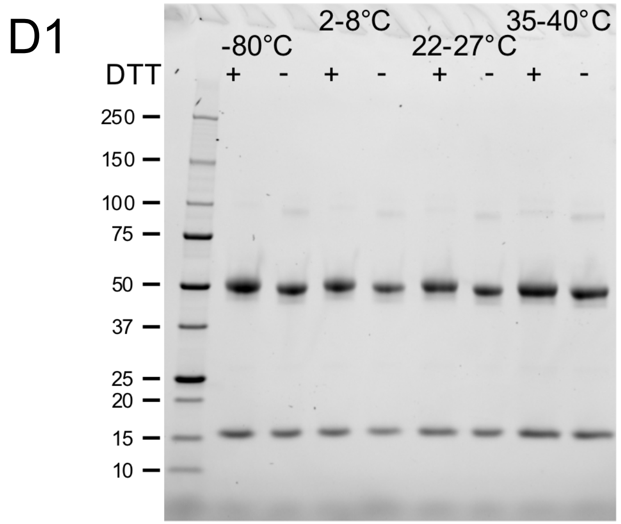


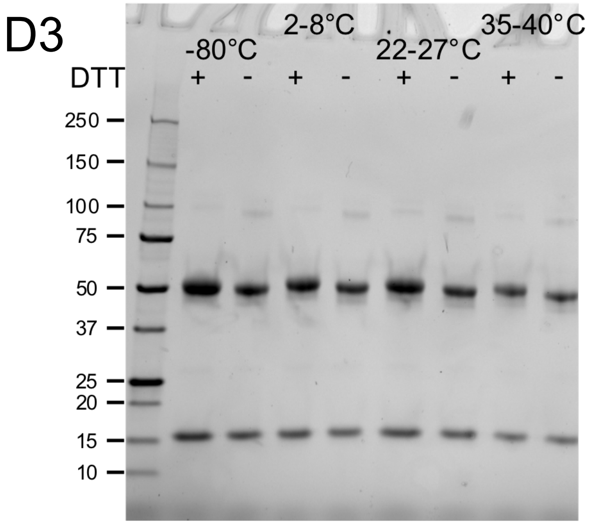


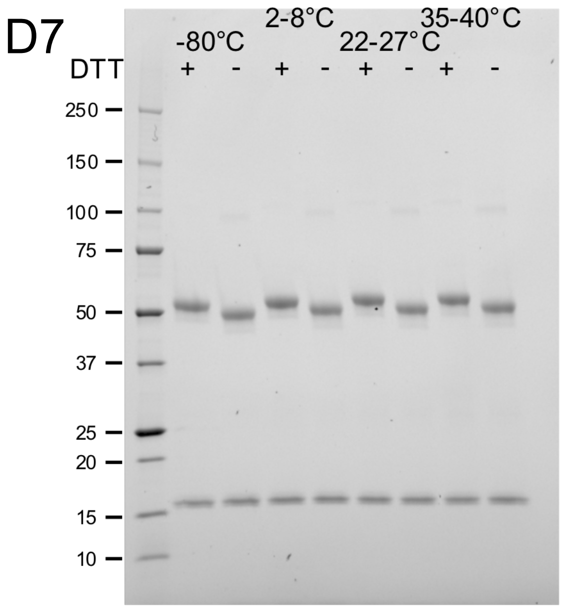


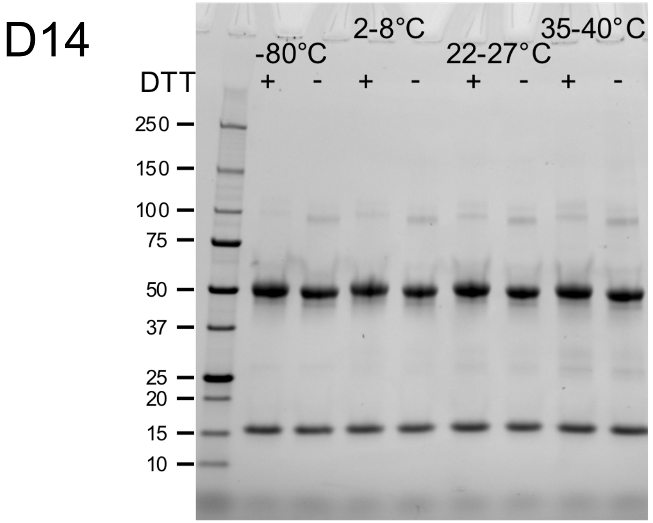


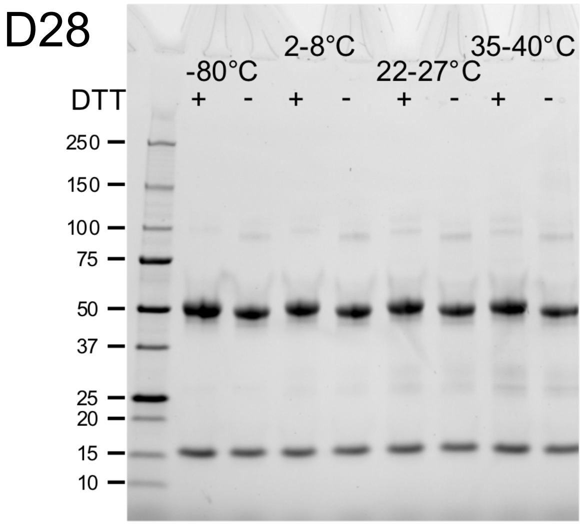


The integrity of samples after incubation at four temperatures over a 28 day (D) study was analyzed by SDS-PAGE. Molecular weights of the standard are noted in kDa. Each sample was analyzed +/- reducing agent (DTT).

### hACE2-Fc binding for Rpk9-I53-50 nanoparticle

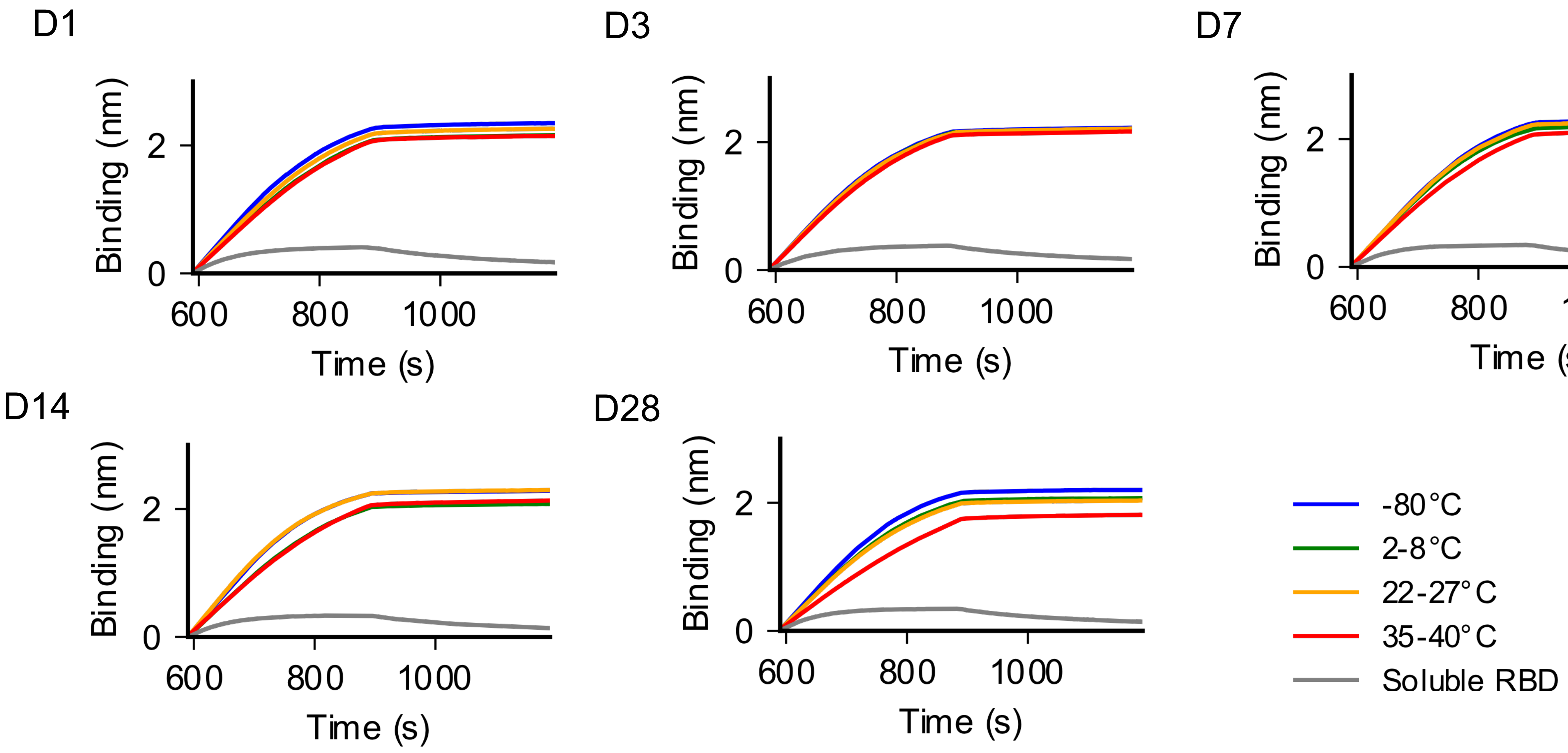


hACE2-Fc binding of antigen incubated at four different temperatures for 28 days (D) was analyzed by bio-layer Interferometry (BLI). Protein A biosensors loaded with hACE2-Fc were incubated with immunogen (association, x = 590–889 s) and then buffer (dissociation, x = 890–1190 s).

### CR3022 binding for Rpk9-I53-50 nanoparticle

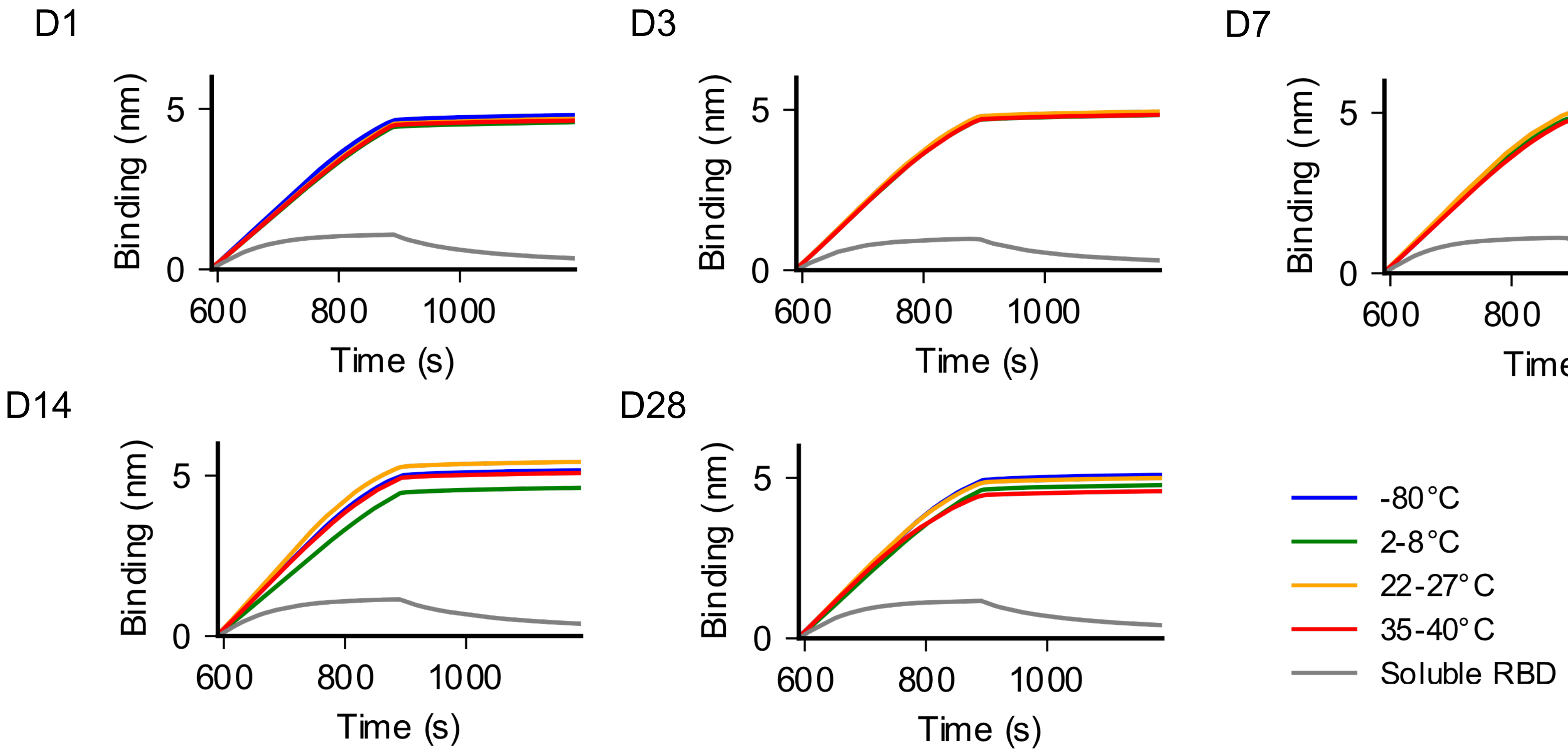


CR3022 IgG binding of antigen incubated at four different temperatures for 28 days (D) was analyzed by bio-layer Interferometry (BLI). Protein A biosensors loaded with CR3022 were incubated with immunogen (association, x = 590–889 s) and then buffer (dissociation, x = 890–1190 s).

### nsEM for Rpk9-I53-50 nanoparticle

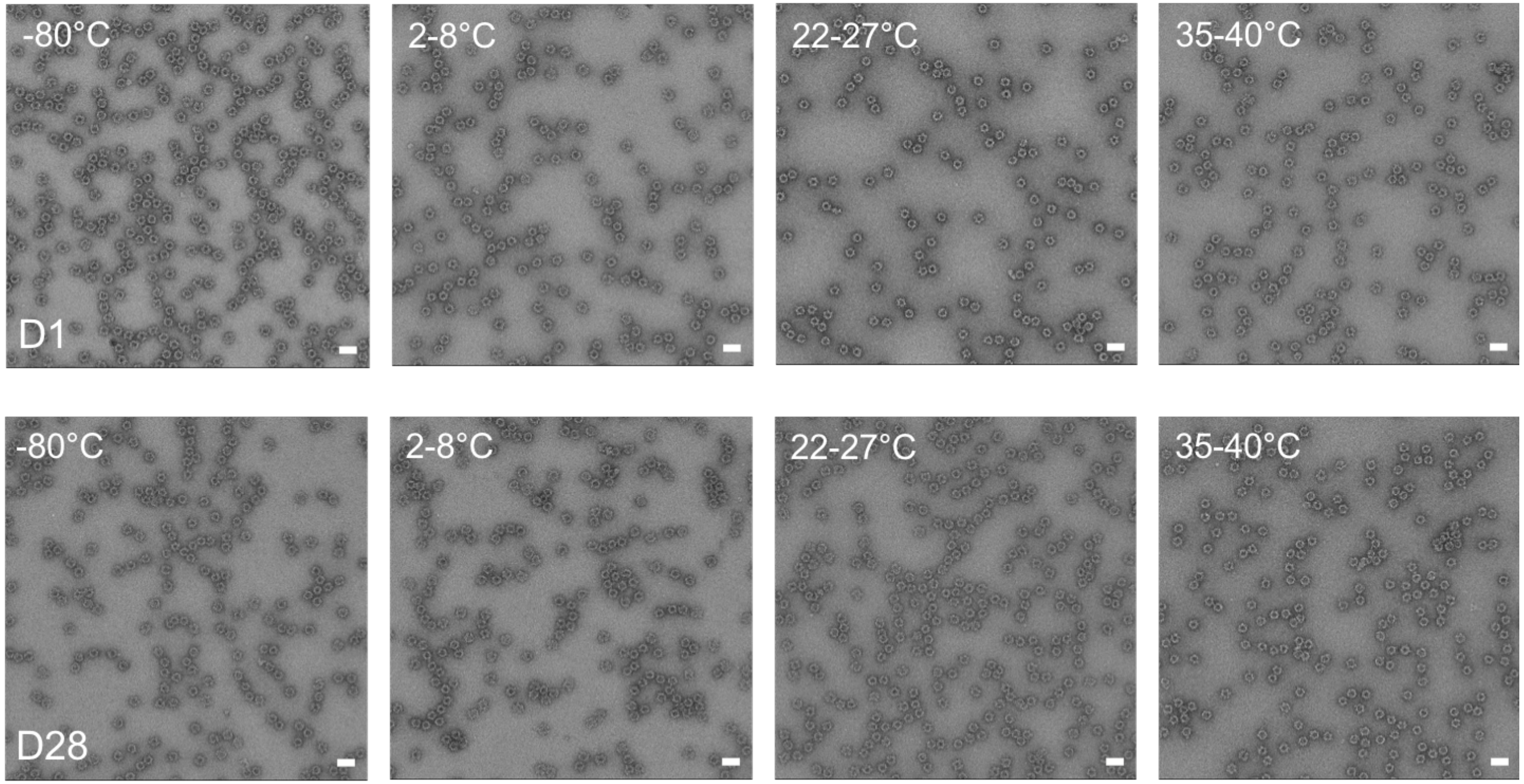


Representative negative stain electron micrographs for each sample at days (D) 1 and 28 following incubation at four temperatures. Scale bar, 50 nm.

### Dynamic light scattering for Rpk9-I53-50 nanoparticle

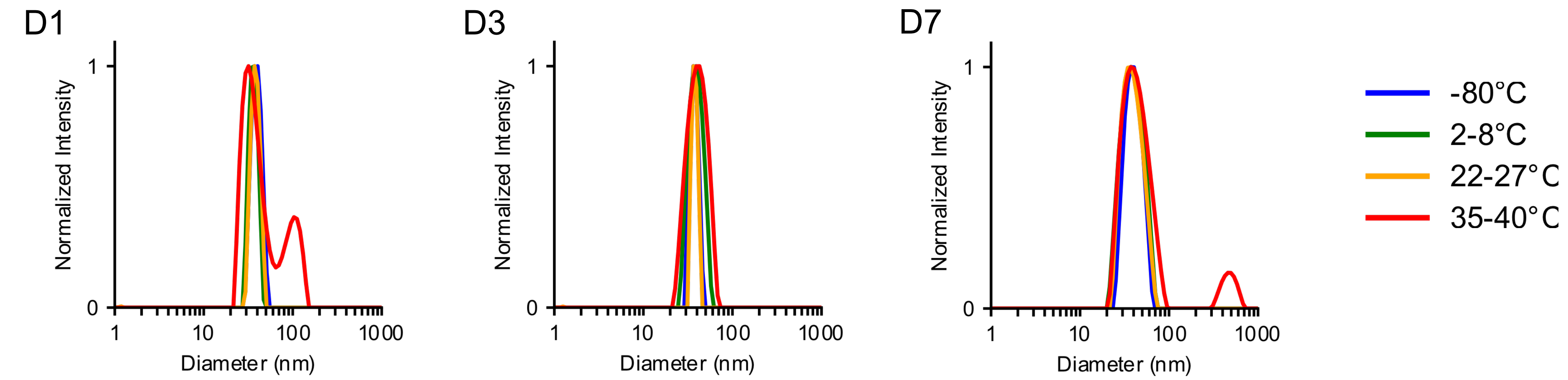


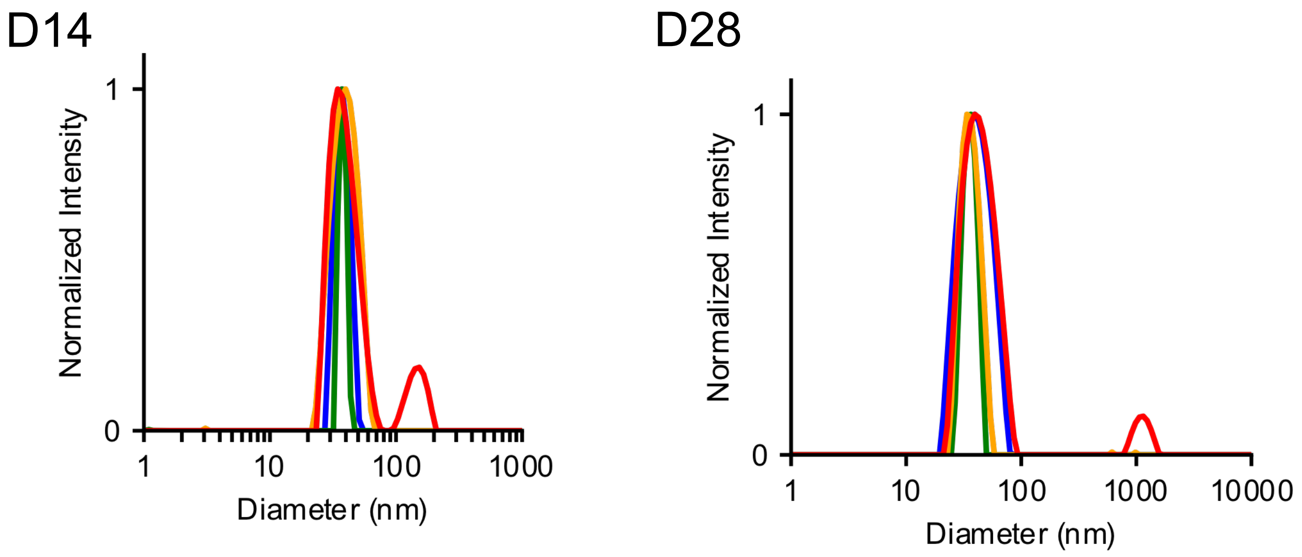


Hydrodynamic diameter (nm) for each sample over a 28 day (D) period, plotted as normalized intensity.
